# CHORUS: A proteome-scale atlas of membrane protein interaction interfaces predicted from sequence

**DOI:** 10.64898/2026.10.04.756538

**Authors:** Jingyao Zhao, Siming Huang, Jianfeng Sun

## Abstract

Membrane protein interactions underpin essential cellular processes, yet the residues mediating these interactions remain difficult to identify at scale due to the paucity of experimentally resolved protein complexes within hydrophobic bilayers. Here, we present CHORUS, a sequence-only framework for predicting interaction interfaces in membrane proteins without requiring structural coordinates, multiple sequence alignments, or known interacting partners. By synthesizing complementary latent representations from three protein language models, CHORUS outperforms 18 established sequence- and structure-based methods by a chain-macro average precision value of 0.643 versus 0.613 for ScanNet-MSA on an independent, homology-restricted test set of 4,655 protein chain contexts, with a particularly pronounced advantage within the membrane-embedded interior (0.746 versus 0.645). Using six mammalian mitochondrial respiratory supercomplexes with >100 independent chains, we show that CHORUS-identified intersubunit interaction interfaces accurately delineate the boundaries between buried protein-protein contact surfaces and surrounding lipid-facing regions in complex molecular systems (0.945-0.984). We subsequently applied CHORUS at proteome scale to chart a global landscape of membrane-protein interaction interfaces for 80,619 expert-reviewed membrane protein entries, 70,247 unique sequences, 4,918 species, and 33.7 million residues across the tree of life. The resulting open-access CHORUS Atlas (https://chorus.atlas.sunnylab.org) provides predicted residue interactions across the membrane proteome. Together, CHORUS establishes a sequence-based foundation for systematic investigation of membrane protein assembly and function, potentially facilitating interactome-scale knowledge discovery and allosteric therapeutic design.

## Introduction

Network edges in protein interactomes effectively chart cellular architecture^1^, yet they fall short of resolving mechanistic causality at the molecular scale^2^. Instead, insights into underlying mechanisms reside in the physical interfaces through which proteins recognize one another^3^, assemble into higher-order structures^4^, and propagate conformational change^5^. This distinction is particularly consequential for membrane proteins, which account for approximately one-quarter of human protein-coding genes^6^ and perform essential functions as receptors^7^, ion channels^8^, active pumps^9^, solute carriers^10^, transporters^11^, and membrane-embedded enzymes^12^, coordinating a plethora of cellular activities, such as signal transduction^13^, electrical excitability^14^, nutrient transport^15^, and energy conversion^16^. Across these classes, interaction and function are fundamentally inseparable: HRD1 recruits SEL1L to organize ER-associated degradation^17,18^. NaDC3 couples transport to asymmetric dimeric states^19^. CaSR activation is coordinated within an obligate homodimer^20^, and Kv4 channels are functionally remodeled by KChIP and DPP6 auxiliary subunits^21^. Such examples illustrate that membrane-protein interfaces are not merely sites of physical contact; they are operative engines of biological regulation.

The pervasive reliance of protein function on molecular interactions raises the dual challenge of defining cellular interactors and deciphering how their interfaces are physically encoded. This objective mirrors the steady progression of interactomics toward higher spatial resolution. Landmark screening efforts such as HuRI first established binary proteome wiring^22^, whereas platforms such as BioPlex^23^ and hu.MAP^24^ extended this view to the organization of multiprotein complexes. Yet, because co-complex membership frequently masks indirect associations, subsequent efforts such as DirectContacts have sought to disambiguate the true direct contacts within these macromolecular machines^25^. In parallel, proteome-scale deep-learning predictions and structural atlases are now translating topological networks into three-dimensional models with residue-level fidelity^26,27^. Collectively, these advances mark a transition from merely cataloging interactions to resolving the structural principles that dictate macromolecular recognition. However, this structural transition faces a formidable bottleneck in membrane proteins. Their functional interfaces are distributed across chemically antithetical compartments (i.e., spanning hydrophobic bilayers^28^ and flanking aqueous environments^29^) and the vast majority remain intractable to experimental complex determination or partner-resolved modeling. Deciphering interaction propensity directly from an isolated polypeptide sequence therefore represents an urgent yet under-addressed computational imperative, bypassing the requirement for predefined binding partners or resolved 3D complexes.

Predicting protein-protein interaction (PPI) interfaces at residue resolution provides the critical bridge between topological networks and the biophysical determinants of protein assembly^30^. Over the past decade, this field has bifurcated into structural and sequence-based computational paradigms^31,32^. Structure-based approaches have advanced rapidly through geometric deep learning^33^. MaSIF-site pioneered the extraction of surface fingerprints^34^, while frameworks such as ScanNet^35^ and PeSTo^36^ parse local atomic geometries to identify binding patches. Yet, the limited structural coverage of membrane proteins (i.e., around 3-5% of high-resolution 3D structures in the PDB^37,38^) renders sequence-based prediction an indispensable alternative. Early deep-learning frameworks, including DeepPPISP^39^, DELPHI^40^, and PIPENN^41^, demonstrated that integrating sequence-derived physicochemical and evolutionary features could effectively map interfaces without 3D coordinates. This paradigm has been further accelerated by protein language models (pLMs)^42–44^. Architectures such as ISPRED-SEQ^45^, Seq-InSite^46^, and PIPENN-EMB^47^ now exploit pretrained representations to extract rich contextual constraints directly from primary sequences. For the membrane proteome, specialized tools including MBPred^48^ and DeepTMInter^49^ further enforce topological and trans-bilayer priors. Despite these advances, predicting protein interfaces in a partner-independent manner remains a grand challenge. A sequence-only, unpaired framework must identify binding-competent residues across hydrophobic bilayers and aqueous compartments without prior knowledge of the interacting partner or quaternary context. Overcoming this limitation provides an essential, template-free foundation for deciphering the molecular wiring of uncharacterized membrane proteins.

Here, we introduce CHORUS, an end-to-end, sequence-only deep-learning framework designed to decipher residue-level interaction interfaces across membrane proteins without requiring structural coordinates, multiple sequence alignments (MSAs)^50^, or cognate partner identities. Rather than relying on a single static model, CHORUS leverages an optimal ensemble that systematically synthesizes complementary latent representations extracted across different model families (ESM-2^51^, ESM-C^52^T5^53^) and layer depths. Through rigorous retrospective benchmarking on non-redundant targets (at least <30% sequence identity to the training set) and diverse external cohorts, CHORUS consistently outperforms existing sequence-based predictors across disparate membrane topologies, interaction partner classes, and protein families. Systematic probing reveals how hierarchical pLM layers encode distinct biophysical properties of interfaces, while downstream analyses demonstrate its capacity to decouple protein interfaces from ligand-binding sites. Crucially, when applied to a mammalian respiratory supercomplex, CHORUS accurately recovers critical contact hubs across multi-subunit membrane assemblies directly based on sequences alone. Finally, to translate this computational capability into a community-wide resource, we establish the CHORUS Atlas (https://chorus.atlas.sunnylab.org), which spans 4,918 eukaryotic and prokaryotic species across the tree of life. This searchable atlas delivers precomputed, residue-level interaction landscapes for all 80,619 reviewed *α*-helical transmembrane proteins deposited in UniProt^54^, encompassing systematic inferences across more than 33.7 million individual amino acid residues. By democratizing proteome-scale interface maps without requiring structural inputs, CHORUS provides a foundational platform to potentially accelerate mechanistic discovery, functional annotation, and targeted therapeutic intervention across uncharted membrane proteomes.

## Results

### Overview of the CHORUS framework

Characterizing protein-protein interaction (PPI) sites across the membrane proteome is crucial for understanding membrane-associated signalling yet remains challenging due to the scarcity of atomic-resolution structures with interacting partners and the limited reliability of multiple sequence alignments (MSAs) in hydrophobic transmembrane domains^48^. To circumvent these limitations, we formulated the prediction of interaction sites in the partner-agnostic context (**Fig. 1**). Given the amino acid sequence *x* of a membrane protein, our proposed model, CHORUS, estimates *p*_*i*_ = Pr(*y*_*i*_ = 1|*x*), the conditional probability that each residue *i* belongs to an interaction interface across intramembrane, transmembrane, and extramembrane regions (**Fig. 1a**). This formulation computationally decouples interface prediction from explicit partner specification, *i.e.*, rather than predicting interaction sites conditional on a predefined interacting partner. The whole sequence-based inference framework therefore requires neither partner information nor complex templates, structural models, or MSAs.

**Fig. 1.**
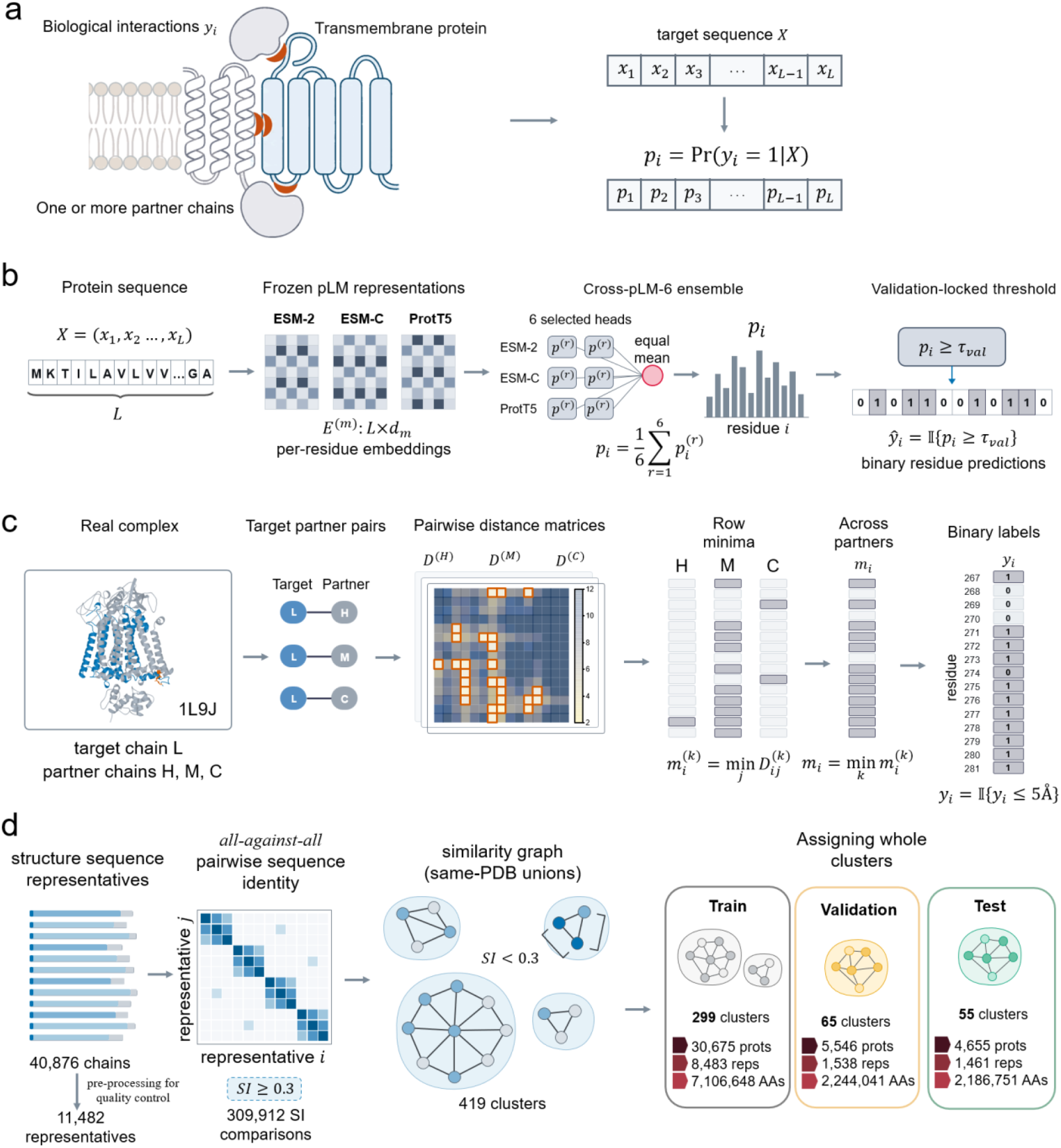
Dataset construction for predicting membrane-protein interaction sites. **a**, Illustration of mapping from the sequence of a membrane protein to probabilities of interacting or non-interacting residues. Predictions are generated without requiring interacting partners, experimental/predicted structural templates, or precalculated multiple sequence alignments (MSAs). **b**, Schematic of sequence-only inference of interaction sites. Target sequences are encoded into protein embeddings 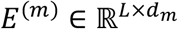 using three protein language models (pLMs), including ESM-2, ESM-C, and ProtT5. For each residue, six selected prediction heads are used to convert its representation to an unweighted mean probability indicative of interacting or non-interacting using a validation-determined cutoff (*τ*_val_ = 0.554). **c**, Structure-derived ground-truth extraction. Pairwise heavy-atom distance matrices (*D*^(*k*)^) are calculated between target chains and their interacting partner chains, e.g., target chain L interacting with chains H, M, and C in an example protein complex (PDB code: 1L9J). Per-partner row minima 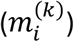 are collapsed to an overall minimal inter-chain distance per residue 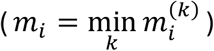. A 5Å heavy-atom cutoff defines binary ground-truth interacting residues 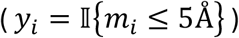. Residues 267-281 are exemplified to illustrate the resulting interface mask, with interacting residues in orange. **d**, Homology- and assembly-aware graph partitioning. A total of 40,876 protein chains are initially collapsed to 11,482 structure-sequence representatives. Pairwise comparisons identify 309,912 similarity edges at ≥ 30% sequence identity (SI ≥ 0.30). The union of this sequence similarity graph and shared intra-PDB memberships forms 419 isolated topological clusters, which are then assigned to training (299 clusters; 30,675 chain instances; 7,106,648 residues), validation (65 clusters; 5,546 chain instances; 2.24M residues), and test partitions (55 clusters; 4,655 chain instances; 2.19M residues).

As seen in **Fig. 1b**, CHORUS feeds embeddings of given residues from three complementary protein language model (pLM) families, including ESM-2^51^, ESM-C^52^, and ProtT5^53^, which are then connected to three neural network architectures (also called heads or backbones) to capture rich evolutionary and structural information. The predicted probabilities of a residue involved in protein interactions are aggregated via an unweighted ensemble mean into *p*_*i*_, which is finally converted into binary predictions (*ŷ*_*i*_ ∈ {0,1}) based on a validation-tuned decision threshold (*τ*_val_ = 0.554).

Residue interactions were derived directly from experimentally resolved multimeric complexes (**Fig. 1c**). For each membrane protein target, heavy-atom Euclidean distance matrices^55^ were computed against all interacting partners, as illustrated by chain L interacting with chains H, M, and C in an exemplified complex (PDB code: 1L9J). Residues were considered as interacting (*y*_*i*_ = 1) if their global minimum distance fell within 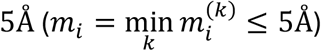. Parsing distinct experimental complex structures resolved at < 3.5 Å yielded 40,876 chains (**Fig. 1c,d**).

To remove redundancy between proteins, we instituted a graph-partitioning protocol (**Fig. 1d**) through which a total of 11,482 chains were identified to be representative of the initial set of PDBTM-derived 40,876 chains following an all-against-all pairwise alignment. This gave rise to exclusion of 309,912 edges at ≥ 30% sequence identity. Taking the connected components of this protein-similarity graph in conjunction with chains from intra-PDB co-assemblies resolved the remaining chains into 419 strictly disjoint clusters. According to a rough ratio 10:1:1, proteins of the entire 419 clusters were split to training (299 clusters, 30,675 chains, 7.11M residues), validation (65 clusters, 5,546 chain, 2.24M residues), and independent test partitions (55 clusters, 4,655 chains, 2.19M residues) (**Supplementary Table 1**). This protocol restricts protein chains such that any pair, including those within and across the training, validation, and test sets, shares less than 30% sequence identity and had no structural co-complex association. Note that identical protein sequences may exhibit different PPI sites in different structural conformations (**Supplementary Fig. 1**). Accordingly, we retained structure-specific chain contexts and their corresponding interacting residue labels.

### Orchestrating cross-pLM representation diversity into CHORUS

Our framework is designed to predict probabilities of interacting residues from sequences alone by extracting frozen embeddings from pretrained protein language models (pLMs) and routing them through downstream neural network architectures (i.e., heads/backbones) with diverse inductive biases. Established studies have suggested that embedding layers across distinct pLM families are enriched for different levels of informative biological features^56^, ranging from local physicochemical motifs and secondary structure propensities in early-to-intermediate layers to global tertiary contacts and functional site constraints in deeper layers. Our initial investigation also provides quantitative evidence that representations extracted from different pLMs exhibit distinct geometric structures, with limited overlap in their local embedding-space neighbourhoods (**Supplementary Figs. 2** and **3**; **Supplementary Note 1**). Consequently, maximizing interaction prediction requires jointly optimizing three dimensions, including *i*) cross-family pLM diversity, spanning ESM2, ESMC, and ProtT5, *ii*) pLM representation depth, sampling 19 layers across the three pLMs to capture layer-dependent biological information, and *iii*) downstream inductive bias, integrating these representations with three heads. To avoid disproportionate contributions from longer proteins and overrepresented sequence-similarity clusters, we calculated average precision (AP) separately for each chain context and averaged it either equally across contexts (chain-macro AP) or first within clusters and then equally across clusters (cluster-macro AP) (**Supplementary Fig. 4**; **Supplementary Note 2**). A systematic comparison between various deep learning architectures prioritized the use of gated Conv-BiGRU (GCB), Conv-Transformer (CT), and gated Transformer-BiGRU (GTB) as three downstream heads (**Supplementary Fig. 5**; **Supplementary Table 2**). In total, these combinations generated a pool of 57 standalone models (**Fig. 2a-c**). For each model, we retained the checkpoint achieving the highest chain-macro AP value on the validation dataset was isolated for subsequent ensemble curation.

**Fig. 2.**
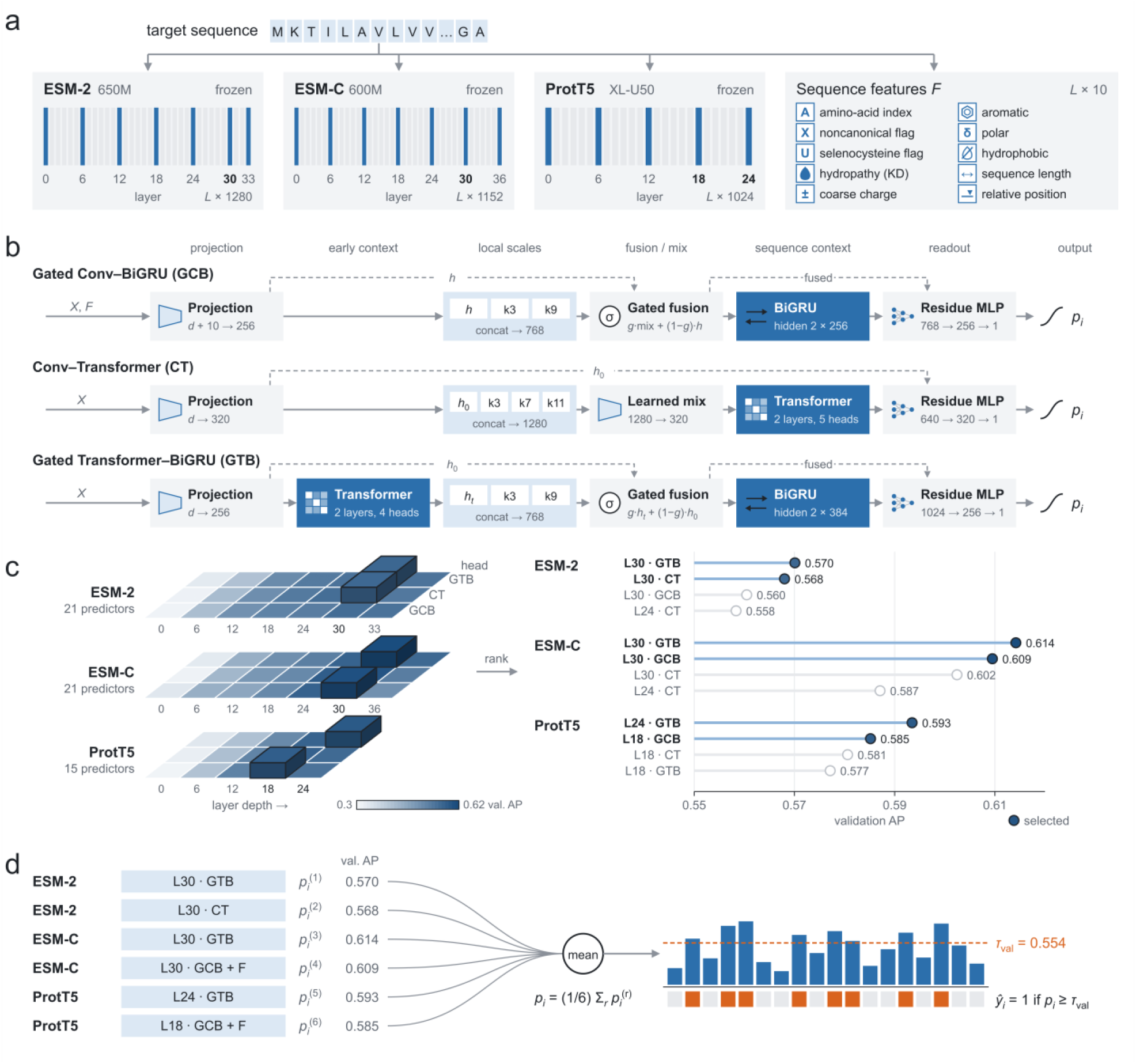
Architecture of CHORUS. a, Pretrained representation extraction and auxiliary priors. Three pLM models (ESM-2 650M, ESM-C 600M, and ProtT5-XL-U50) extract multi-layer protein embeddings per residue. For performance comparison, an auxiliary set of ten biophysical sequence properties (e.g., hydropathy, charge, polarity, and chain position) is incorporated exclusively into feature-augmented Gated Conv-BiGRU (GCB) networks. **b**, Schematic of residue-level prediction backbones. Three hybrid backbones work by combining local and sequential contexts, with Gated Conv-BiGRU (GCB, left) dynamically gating multi-scale 1D convolutions through bidirectional GRUs, Conv-Transformer (CT, middle) fusing multi-scale receptive fields through a 2-layer multi-head transformer encoder, and Gated Transformer-BiGRU (GTB, right) routing transformer-encoded sequence features through gated 1D convolutions and wide BiGRU units. All backbones conclude with a multi-layer perceptron (MLP) and sigmoid activation to yield residue-level interaction probabilities. **c**, Systematic candidate space search. A combinatorial grid of 57 predictors, which spans 19 sampled extraction layers across the above pLM families and backbone archetypes, is evaluated across validation chains. Candidates are ranked within each family by the validation chain-macro AP metric. The top two configurations per family (gold circles) are selected. **d**, Integration of the entire prediction framework. Interaction probabilities are predicted by equally averaging the decisions from six individual models spanning intermediate and deep layers of pLMs, including ESM-2 layer 30 with GTB/CT, ESM-C layer 30 with GTB/GCB, ProtT5 layers 18/24 with GCB/GTB. A validation-determined decision threshold (*τ*_val_ = 0.554) is finally used to convert the predicted probability of a residue being an interaction or non-interaction site.

To preserve foundational representation diversity, constituent model selection was conducted within, rather than across, pLM families. We obtained a six-model ensemble (dubbed CHORUS) by retaining two top-performing models per family across all three pLMs and downstream heads, including ESM2 layer 30 plugged into GTB (val AP: 0.5701) and CT (0.5680), ESMC layer 30 plugged into GTB (0.6142) and GCB (0.6095), and ProtT5 layer 24 plugged into GTB (0.5935) alongside layer 18 with GCB (0.5852) (**Extended Data Fig. 1**; **Supplementary Figs. 6** and **7**; **Supplementary Tables 3** and **4**). Our ablation experiment in a leave-one-model-out test fashion confirmed the necessity for the current choice of these models (**Supplementary Table 5**). Further analysis of the correlation between predicted probabilities of pairwise models in CHORUS highlighted the importance of the multisource, diversified predictions to elevate the ultimate performance (**Supplementary Table 6**; **Supplementary Note 3**).

To assess whether alternative model combinations could improve CHORUS, we constructed multiple strategies (**Supplementary Table 7**; **Supplementary Note 4**) and a two-layer perceptron-based neural selector for model scoring with feature ablations across 63 candidate models (**Extended Data Fig. 2**). Model combinations were selected through exhaustive search, beam search or direct rule-based selection (**Supplementary Table 8**). Our data show that selection performance was metric-dependent, and greater neural-selector complexity did not consistently improve predictive accuracy (**Supplementary Tables 9** and **10**). Next, we examined whether altering residue features and model parameters impacted the performance (**Supplementary Fig. 8**; **Supplementary Note 5**). Our investigation pinpointed that the CHORUS ensemble significantly outperformed all baseline controls constructed using multiple training seeds (0.6210), orthogonally transformed embeddings (0.6178), representation depths (0.5881), identical duplication (0.5752), or downstream heads with an enlarged parameter size (0.5567) (**Extended Data Fig. 3**; **Supplementary Tables 11** and **12**).

### Performance evaluation on the independent test set

We benchmarked CHORUS against a total of 18 established protein interaction-site predictors, encompassing sequence-based methods (DeepTMInter, DeepPPISP, EnsemPPIS^57^, PITHIA^58^, Seq-InSite^46^, PIPENN-EMB^47^, and ALLSites^59^) and structure-based methods that incorporate target-chain geometry (ScanNet^35^, GPSite^60^, PeSTo^36^, GraphPPIS^61^, EquiPPIS^62^, Gated-GPS^63^, RGN^64^, Spatom^65^, DeepProSite^66^, MaSIF-site^34^, and EGRET^67^) (**Supplementary Table 13**). Note that ScanNet was evaluated both with and without MSA features, while GraphPPIS was assessed with its fast and evolutionary-profile configurations. The benchmark was entirely evaluated on our orchestrated independent test set that comprises 4,655 chain contexts across 55 sequence-similarity clusters, with no sequence pair between partitions exceeding 30% sequence identity. All predictions were benchmarked against a standardized 5Å inter-residue contact threshold. A residue is assigned as interacting if its predicted probability is above a threshold of 0.554 determined by maximizing interacting-residue F1 across pooled validation residues and subsequently fixed for test-set evaluation (**Extended Data Fig. 4**). Protein structures required for structure-based predictors were predicted by ESMFold or ColabFold. When both predicted-structure sources were available, the source with higher test chain-macro-AP values was chosen for evaluation. DeepProSite, EGRET, and MaSIF-site were excluded from model ranking due to incomplete benchmark coverage (i.e., sequence length limit).

Overall, CHORUS outperformed all methods, achieving a chain-macro AP value of 0.643, a 0.030 absolute improvement over the best-performing complete-coverage baseline, ScanNet-MSA (0.613), and substantially outpacing leading structure-guided methods, such as GPSite (0.583) and PeSTo (0.581), and sequence-only methods, such as Seq-InSite (0.556) and ESMC-L30 (0.555) (**Fig. 3a**). Using all pooled 2.18 million residues, CHORUS reinforced its discriminative capacity by achieving a pooled-residue AP value of 0.667 and an AUROC value of 0.861, compared with 0.612 and 0.843 for ScanNet-MSA (**Fig. 3a and 3d**). Crucially, CHORUS demonstrates the strongest ability to predict the top-ranked residues as interacting (e.g., 0.826 at P@5, 0.808 at *P*@10, and 0.793 at *P*@20) (**Fig. 3b**). In stark contrast, the performance of competing methods drops off significantly, with ScanNet-MSA declining from 0.821 at *P*@5 to 0.743 at *P*@20 while GPSite and PeSTo plummeting below 0.710 (0.699 and 0.710 at *P*@20, respectively) and all other compared methods falling under 0.685. At the validation-tuned thresholds, CHORUS can strike a balance of classification between evaluation metrics over pooled residues, achieving an F1 of 0.620 and the highest Matthews correlation coefficient (MCC) of 0.509. Notably, whereas competing methods achieved apparent sensitivity by heavily sacrificing specificity, e.g., GraphPPIS-slow and PITHIA driving recall to 0.825-1.000 at the cost of precision dropping to 0.230-0.349, CHORUS balanced high precision (0.632) with robust recall (0.608). Furthermore, probability calibration assessed via the class-balanced Brier score confirmed that CHORUS produced the most reliable posterior probability estimates under class imbalance (Brier = 0.167 for CHORUS vs. 0.189 for ScanNet-MSA, 0.234 for PeSTo, and 0.291 for GPSite).

**Fig. 3.**
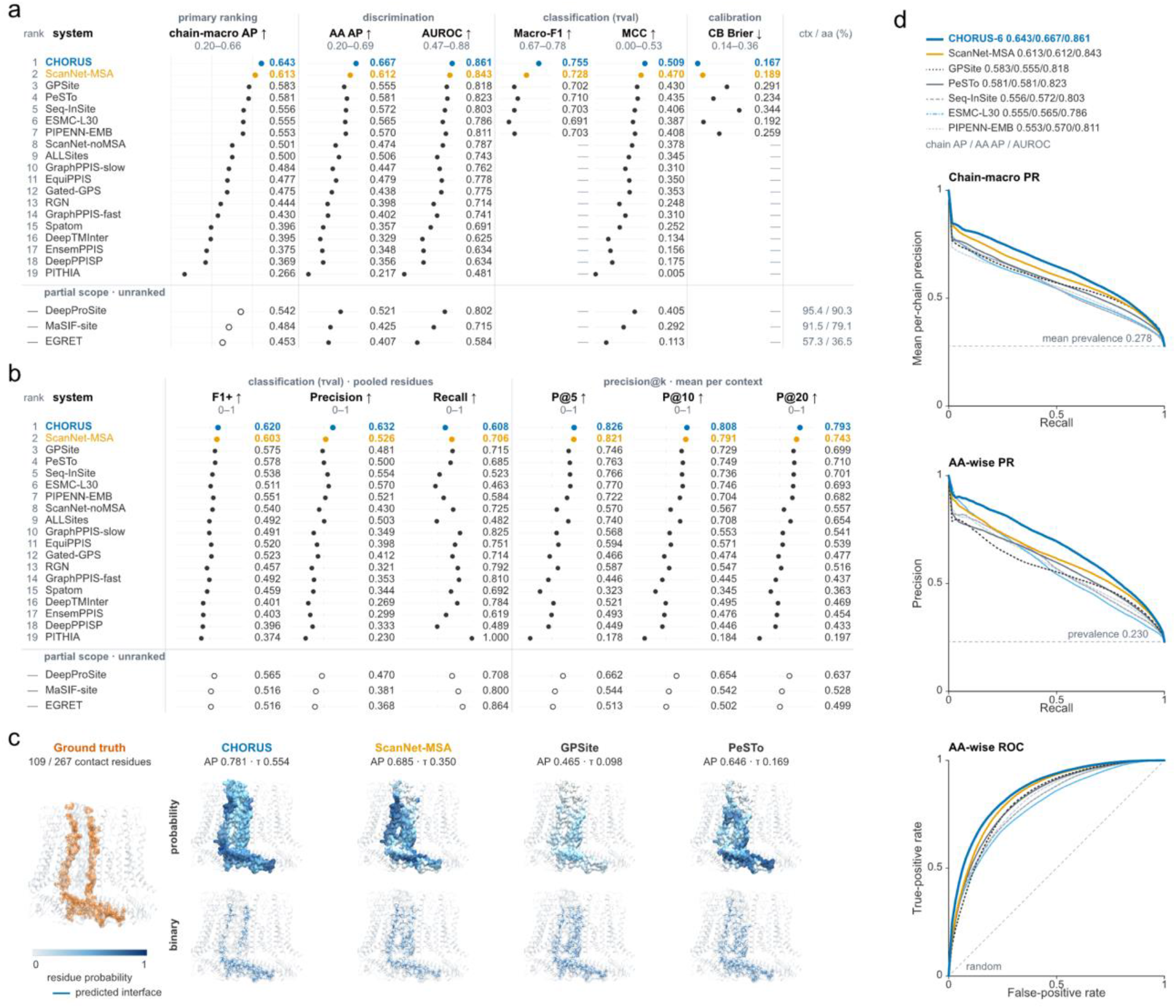
Evaluation of CHORUS against sequence- and structure-based interaction site predictors. **a**, Performance comparison between 19 complete-coverage predictors evaluated across 4,649 chain contexts in the test dataset using various assessment metrics at validation-tuned thresholds. Methods (DeepProSite, MaSIF-site, and EGRET) with limited coverage of chain contexts due to sequence-length requirements are listed separately without ranking, alongside their context/residue coverage percentages (ctx / aa (%)). **b**, Performance comparison based on the mean precision per chain context among the top-ranked 5, 10, and 20 residues with the highest probabilities (*P*@5, *P*@10, and *P*@20). Chains with length <*k* contribute all evaluated residues. Open circles denote partial-coverage methods. **c**, Case study illustrating predicted interfaces in structural contexts using human CALHM2 chain F (PDB code: 6VAI; 109 contact residues out of 267 total residues). CHORUS predictions are evaluated against ground-truth interactions of chain F. Top: residue-level predicted interaction probabilities (blue intensity gradient). Bottom: binary interface predictions (thresholded at method-specific optimal cutoffs, *τ*) against ground-truth interactions (orange silhouette). **d**, Precision-recall curves (chain-macro and pooled amino acids/residues) and receiver operating characteristic (ROC) curves comparing CHORUS with top-performing baselines. Dashed horizontal/diagonal reference lines denote class-positive prevalence, i.e., the fraction of interacting residues and random classification performance. The full test set encompasses 4,655 chain contexts and 2,186,751 residues (502,137 interacting residues). Six contexts devoid of interacting residues were excluded from AP calculations. Binary cutoffs were optimized on validation data via positive-class F1 and held fixed during testing.

This quantitative advantage translated directly into spatially coherent interface delineation on a representative benchmark target, the Cryogenic electron microscopy (cryo-EM) structure of a dimer of the undecameric human CALHM2 (PDB code: 6VAI, chain F), which comprises 109 interacting residues out of 267 total residues (**Fig. 3c**). CHORUS achieved a chain-macro AP value of 0.781, accurately delineating the contiguous interaction patch. By contrast, other approaches failed to fully capture interface geometry or suffered widespread false positives, with lower performance, e.g., a chain-macro AP value of 0.685 for ScanNet-MSA, 0.646 for PeSTo, and 0.465, for GPSite. Furthermore, precision-recall and ROC curves collectively demonstrated consistent superiority of CHORUS across decision thresholds (**Fig. 3d**).

### Performance evaluation on external datasets

To rigorously assess model generalizability beyond our internal splits, we first evaluated CHORUS on three established benchmarking datasets specialized for transmembrane proteins, including CompData101 (101 chains)^68^, TestData36 (36 chains)^48^, and IndepData30 (30 chains)^49^ (**Supplementary Table 14**). To prevent data leakage from sequence homology, we retrained CHORUS on protein chains filtered at < 30% sequence identity against the external datasets (14,625 chains for training and 4,112 chains for validation). Evaluated across all 167 non-redundant external membrane protein targets, our ensemble model retained the most robust predictive ability, with chain-macro-AP values of 0.770, 0.741, and 0.817 on CompData101, IndepData30, and TestData36, respectively (**Supplementary Table 15**). These results show that CHORUS captures interface physics rather than relying on sequence homology.

### Sensitivity analysis of prediction performance across pLM layers

To investigate how biological features encoded at different pLM layers contribute to predicting interacting residues, we benchmarked the chain-macro AP performance of 57 models in multiple scenarios (**Fig. 4**; **Supplementary Note 6**). Notably, the terminal layer of pLMs was found to be less informative for identifying interaction sites in membrane proteins (left panel, **Fig. 4a**). But the predictive performance of almost all models were shown to peak at later-stage layers, e.g., layer 30 for ESM2 as well as ESMC and layer 18 for ProtT5. Meanwhile, it was observed that none of prediction heads consistently dominated across pLM representations. This heterogeneity underscores the importance of jointly optimizing the pLM representations and downstream heads to translate protein embeddings into accurate likelihoods of residues being interaction sites. In addition, we found that imposing orthogonal transformation on off-the-shelf protein embeddings to train those constituent models in CHORUS was significantly detrimental to the ultimate predictive performance (0.618 versus 0.643; right panel, **Fig. 4a**). The contribution of protein embeddings to the CHORUS performance is contextualized within a broader comparative framework (**Supplementary Tables 11** and **12**; **Supplementary Note**).

**Fig. 4.**
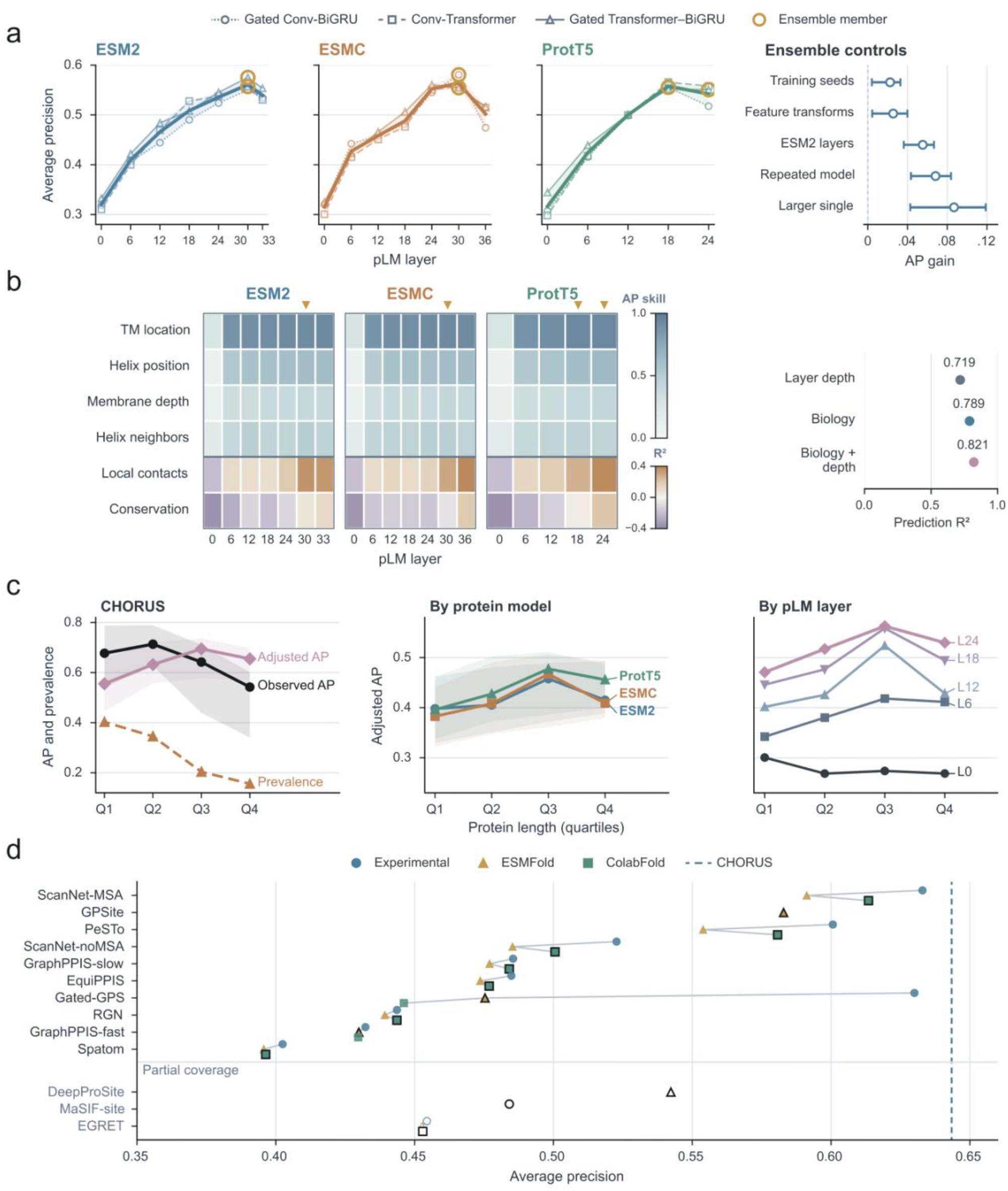
Sensitivity analysis of predictions using pLM representations with respect to biological properties. **a,** Representation benchmarks and ensemble controls. Left: chain-macro AP performance across 57 predictors combining three pLMs, 19 layer representations, and three downstream heads). Curves denote head means, shading shows ranges across heads, and amber circles highlight the six CHORUS members. Right: chain-macro AP performance versus five controls (different training seeds, orthogonally transformed protein embeddings, ESM2 layer combinations, identical replicas, and an enlarged head). Error bars indicate 95% bootstrap confidence intervals (10,000 resamples across 54 test clusters; paired differences lie strictly above zero). **b,** Biological probing and performance comparison. Left: biological information recovered from frozen pLM representations across layer depth. The top four rows depict AP skill, calculated as 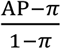 where *π* indicates the fraction of interacting residues for predicted properties related to transmembrane (TM) topologies and depths. The bottom two rows report *R*^2^ for predicted local contacts and evolutionary conservation. Amber markers denote CHORUS layers. Right: leave-one-pLM-out ridge regression predicting AP gains over layer 0 using depth, biological probe scores, or both (*R*^2^ shown). **c,** Performance across protein length quartiles. Left: observed AP, positive prevalence, and prevalence-adjusted AP (standardized to validation prevalence *π* = 0.2332) for CHORUS across sequence length quartiles (Q1-Q4: 20-288, 289-424, 426-614, and 615-1,380 residues). Middle and right: adjusted AP for a balanced set of 45 predictors, broken down by pLM family and layer depth. Shading indicates 95% cluster-bootstrap intervals (2,000 resamples). **d,** Sensitivity of structure-based methods to the source of 3D structures (experimental, ESMFold, or ColabFold). Solid lines denote the three structures as inputs for each method. Black borders mark the optimal benchmark input. Dashed vertical lines mark CHORUS. Methods below the horizontal grey line exhibit partial coverage of the test chain contexts (DeepProSite: 95.4%; MaSIF-site: 91.5%; EGRET: 57.3%).

Next, we investigated how a variety of membrane protein-associated biological properties are represented across pLM layers (**Fig. 4b**; **Supplementary Fig. 9**). We first found that the predictability of geometric and topological features, including transmembrane (TM) location (interior vs. flanking), helix position, and membrane depth, was largely insensitive to pLM layer depths, while local contacts and evolutionary conservation progressively became informative as pLM layers deepened (left panel, **Fig. 4b**; **Extended Data Fig. 5**). Using the L2-regularized ridge regressor, we then explored how these biological properties embedded in pLM layers positively impacted the chain-macro AP performance of the interaction-site prediction. The results showed that probe scores gained from predicting these biological properties (i.e., AP skill and goodness of fit) were correlated strongly with pLM layer depths, together contributing to the AP gain of CHORUS (*R*^2^ = 0.821) (right panel, **Fig. 4b**). In line with these findings, our additional studies revealed that biological properties and pLM layer depth explained largely overlapping variation in AP gains, whereas representation diversity alone explained little of the ensemble improvement (**Supplementary Figs. 10 and 11**). As the raw chain-macro AP performance was rendered to vary largely with the percentage of resides annotated as interacting (leftmost panel, **Fig. 4c**), we adjusted AP to a common percentage of 23.32% (*π* = 0.2332) based on the validation dataset to enable fairer comparisons across protein lengths. The analysis of predictive performance regarding the length of protein sequences and the percentage of interacting residues unravelled that utilizing protein embeddings from deeper layers was consistently beneficial to achieve high performance across all length settings (from Q1 to Q4 quartiles) (middle and rightmost panels, **Fig. 4c**).

Finally, further benchmarking highlighted that methods requiring explicit 3D conformations suffered a drop in terms of the chain-macro AP performance when moving from experimental coordinates to ESMFold or ColabFold models, e.g., Gated-GPS declining significantly from 0.630 to 0.475 and 0.446, respectively, while GraphPPIS-slow declining modestly from 0.485 to 0.477 and 0.484, respectively (**Fig. 4d**). Several structure-dependent tools also had limited coverage of the tested chain contexts (e.g., 57.3% for EGRET). In contrast, sequence-only CHORUS worked on full coverage without requiring structural preprocessing.

### Performance evaluation across membrane regions and biological contexts

To determine the structural and evolutionary factors underlying interaction site prediction methods, we systematically evaluated their performance across a multitude of contexts, including sub-cellular microenvironments, interaction partners, topologies, and protein families (**Fig. 5**). To prevent over-represented protein families from biasing the evaluation, metrics across biological subgroups were macro-averaged within homology clusters (30% sequence identity, SI30) and reported as median cluster average precision (mAP) alongside interquartile ranges (IQR).

**Fig. 5.**
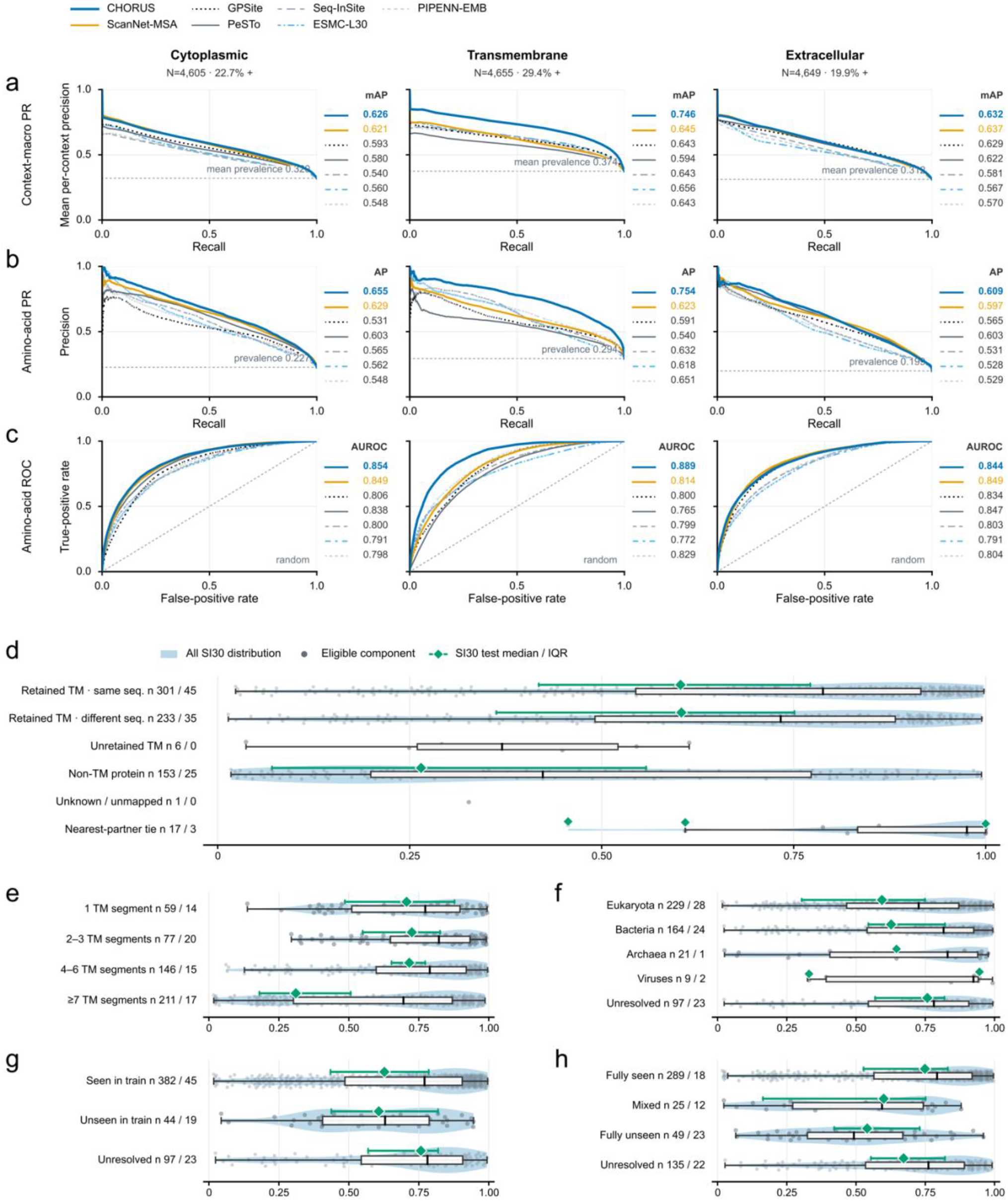
Analysis of predicted interaction sites across membrane topologies, biological constraints, taxonomic classes, and protein families. **a-c,** Performance of predicted interaction sites stratified by membrane microenvironments. Precision-recall (context/chain-macro, **a**; pooled residues, **b**) and ROC (pooled residue, **c**) curves for cytoplasmic, transmembrane, and extracellular regions, respectively, evaluated on the independent test set (*N* = 4,655 contexts). CHORUS is benchmarked against five external predictors and the internal ESMC-L30 baseline. Dashed lines denote mean baseline prevalences or random discrimination. **d-h,** Performance across protein classes (**d**), target transmembrane segment counts (**e**), taxonomic domains (**f**), species-level presence/absence of proteins during training (**g**), and Pfam-derived homology-level presence/absence of proteins during training (**h**). In all panels, violins illustrate distributions across the entire dataset (muted blue), while green diamonds and error bars represent medians and interquartile ranges (IQR) across independent-test clusters (shown for categories with ≥ 4 test clusters).

We first assessed whether different methods displayed a region-specific difference in predictability across the lipid bilayer (**Fig. 5a-c**; **Supplementary Figs. 12 and 13**). Our assessment showed that while two top-performing groups, ScanNet-MSA and CHORUS, delivered comparable precision in aqueous environments, with a mAP of 0.626 versus 0.621 at the cytoplasmic region and 0.632 versus 0.637 at the extracellular region, the CHORUS performance stayed ahead way more than ScanNet-MSA within the intramembrane core in terms of a chain-macro AP value of 0.746 versus 0.645 (pooled AP: 0.754 versus 0.623, AUROC: 0.889 versus 0.814). This pronounced advantage demonstrates that sequence-derived representations from diverse pLMs can effectively capture the tightly packed characteristics of transmembrane helices in a low-dielectric environment, negating the need for 3D atomic coordinates.

We also dissected residue interactions that occur between membrane and soluble proteins (**Fig. 5d**). If interacting residues lie at membrane protein sites, CHORUS maintained consistently high precision across permanent interfaces in both homo-oligomeric protein complexes (median AP: 0.603) and hetero-oligomeric protein complexes (different-sequence partners: median AP: 0.604) on 45 and 35 independent test clusters, respectively. In stark contrast, predictions degraded substantially for transient interfaces formed with soluble proteins (median AP = 0.265 across 25 clusters). This divergence indicates that CHORUS has benefitted deeply from learning lipid-embedded helix-helix packing motifs based on a miscellaneous array of membrane proteins using protein embeddings across pLM families.

Yet another analyses of prediction accuracy across TM segments, taxonomic groups, species-level generalizability, and protein-family homology were carried out to identify the biological contexts in which CHORUS performs most reliably (**Fig. 5e-h**). CHORUS was shown to remain robust for bitopic and oligotopic proteins containing 1 to 6 TM segments, with median cluster AP values of 0.706-0.725 (**Fig. 5e**). However, it dropped sharply for polytopic membrane proteins characterized by ≥ 7 TM segments, with median AP value of 0.311 across 17 test clusters. Such polytopic bundles are inextricably intertwined with intricate intramolecular contacts that often confound the identification of intermolecular interactions. Moreover, we discovered that although CHORUS predictions generalized seamlessly across major kingdoms, e.g., mAP 0.627 for Bacteria and 0.594 for Eukaryota (**Fig. 5f**) and between species seen or unseen during training (i.e., 0.627 versus 0.607, **Fig. 5g**), the performance was strongly modulated by family-level homology (**Fig. 5h**). Membrane proteins whose SIFTS-Pfam domain families were fully represented in training reached a median AP value of 0.749, dropping dramatically to 0.600 for partially represented families and 0.540 for orphan families unseen during training (23 test clusters). Thus, while taxonomic origins do not restrict model utility, structural family coverage remains a key governing factor.

### Selective generalization of CHORUS to soluble protein interfaces

We applied CHORUS to identify interaction interfaces in soluble assemblies to investigate whether membrane protein interfaces share common features with these proteins (**Supplementary Table 16**). Our evaluation results illuminated that CHORUS generalized well to three soluble coiled-coil oligomers composed of tightly packed assemblies of 2-5 *α*-helices, achieving 0.943-0.960 at AUROC (**Supplementary Tables 17 and 18**). One line of reasoning that could explain the results is that *α* -helical transmembrane proteins are arranged in a tightly packed bundle fashion structurally similar to coiled coils, thereby enhancing the ability of generalization. Meanwhile, it accurately identified interfaces in two protein-inhibitor complexes, eglin C and stefin B, with AUROC values of 0.942 and 0.940. However, the performance became less consistent on enzyme-partner interfaces, with AUROC values ranging widely from 0.469 to 0.869. Meanwhile, we observed that predicted interaction interfaces at our prespecified threshold were saturated with false positives on tetrabrachion and no true positives on ribonuclease inhibitor. These findings substantiate that geometric interaction features learned from membrane environments have helped to generalize to specific soluble protein targets.

### Identifying interaction landscapes across large respiratory supercomplexes

Mitochondrial respiratory supercomplexes are large assemblies of membrane proteins that modulate energy transduction. These higher-order, membrane-embedded supramolecular machineries are assembled with individual respiratory complexes through residue interactions to optimize electron flux. To facilitate the understanding of whether CHORUS can precisely delineate interaction interfaces within a large, physiologically relevant membrane-protein assembly, we pick the circular supercomplex assembly (PDB code: 8UGN) as our primary biological case study (**Fig. 6**). The cryo-electron microscopy structure resolves this massive assembly into 140 independent chains in which 78 are membrane-associated proteins containing 38 unique chains. We applied CHORUS to identify interfaces in a per-chain manner without access to the quaternary assembly context. Across 12,337 residues, CHORUS accurately delineated inter-subunit interacting boundaries, achieving an chain-macro AP value of 0.984, an AUROC value of 0.970, and an MCC value of 0.817 at the 5Å threshold (**Fig. 6a**).

**Fig. 6.**
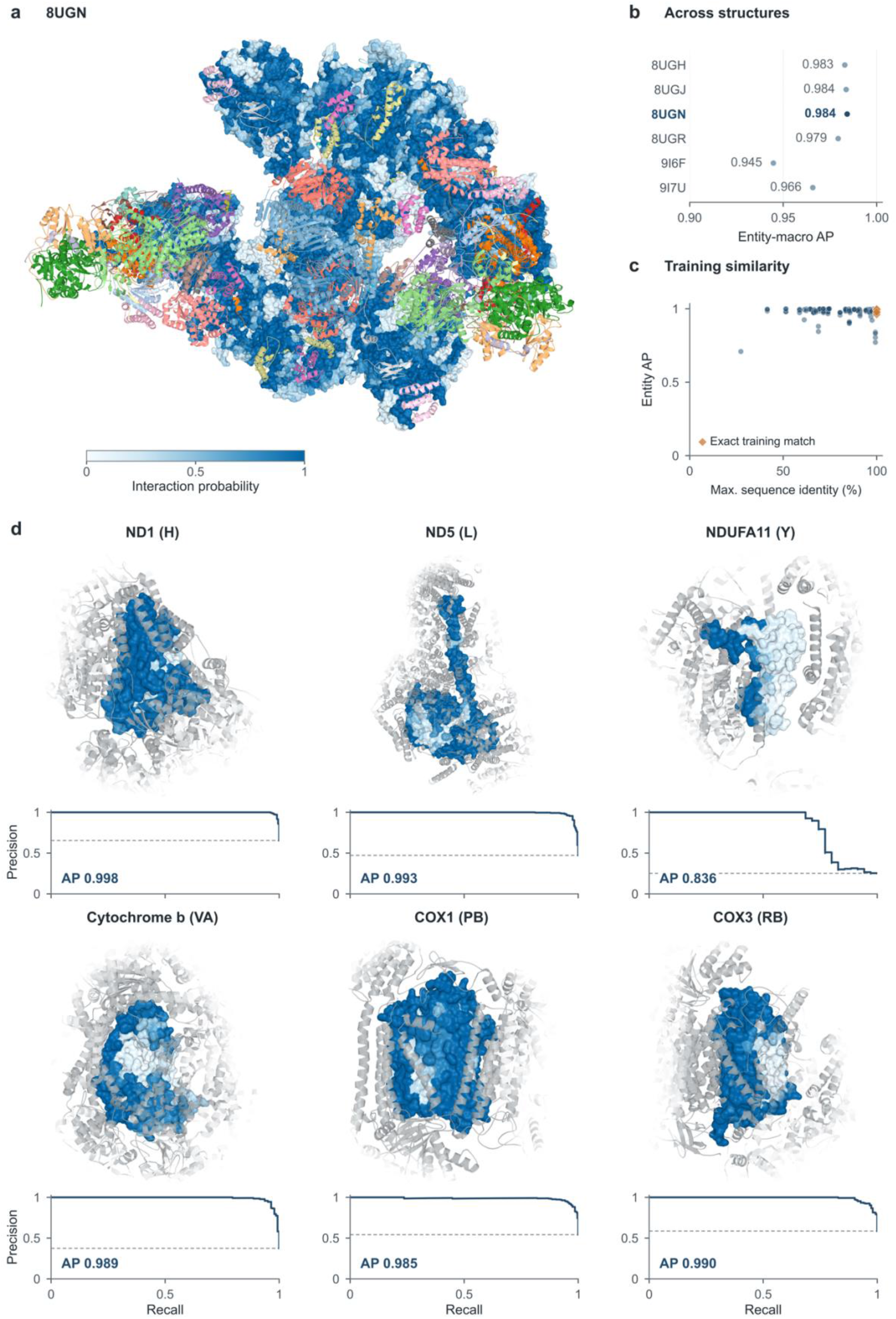
Predicted interfaces across mitochondrial respiratory supercomplexes. **a,** Quaternary interaction landscape of the porcine respirasome. Biological assembly of the *I*_2_*III*_2_*IV*_2_ supercomplex (PDB code: 8UGN, 140 protein chains). The 78 physical copies of 38 target transmembrane protein entities are rendered as molecular surfaces colored by predicted interaction probability (0 to 1, white to blue; six-model uncalibrated ensemble mean). The remaining 62 non-target chains are displayed as ribbons colored by subunit identity. **b,** Consistent accuracy across macromolecular assemblies. Entity-macro AP across six cryo-EM respiratory reconstructions, including four porcine supercomplex states (PDB codes: 8UGH, 8UGJ, 8UGN, 8UGR) and two human complex-IV maturation intermediates (PDB codes: 9I6F, 9I7U). 8UGN is highlighted in bold blue. **c,** Performance across homology strata. Entity-level AP versus maximum sequence identity to the training set across 174 entity-structure instances. Orange diamonds denote exact sequence matches to training entries. **d,** Subunit-resolved interfaces and precision-recall profiles. Structural close-ups and corresponding precision-recall curves for six key core subunits: Complex I (ND1, ND5, NDUFA11), Complex III (Cytochrome b), and Complex IV (COX1, COX3). Target surfaces are colored by predicted probability. Interacting partners are visualized as grey ribbons fading with distance (12-28Å). Horizontal dashed lines mark baseline interacting prevalence. Residue interactions are defined at a 5Å heavy-atom inter-chain cutoff.

To assess reproducibility across distinct quaternary states and species, we evaluated five additional cryo-EM-determined assemblies, spanning three sus scrofa-derived supercomplexes (8UGH, 8UGJ, and 8UGR) and two human-derived supercomplexes (9I6F and 9I7U). Still, CHORUS attained high chain-macro AP values ranging from 0.945 to 0.984 (**Supplementary Table 19**). As seen in **Fig. 6c**, the high accuracy was shown to maintain robustly for most of their subunits across varying degrees of homology to training sequences and was like to arise from high sequence identity (median 90.8%). Nevertheless, some subunits divergent with our training cohort achieved high AP performance. Using structural cutouts displayed in **Fig. 6d**, we showed that the predicted probabilities avail of themselves of separating the buried interaction surfaces from the surrounding membrane-facing lipidic perimeters. These results demonstrate that CHORUS-predicted interaction landscapes aid accurately explain how these native macromolecular supercomplexes might be assembled directly from primary sequence.

### Generalizability across diverse membrane-protein systems

Beyond respiratory supercomplexes, we further evaluated CHORUS across diverse membrane-protein systems involved in photosynthesis, innate immune signaling, mitochondrial protein import, endoplasmic reticulum (ER) translocation, and ion-channel regulation (**Supplementary Table 20**). CHORUS achieved high AP values for PSII–LHCII assemblies (0.900-0.950; **Extended Data Fig. 6**), STING complexes (0.827-0.919; **Supplementary Fig. 14**), and PINK1–TOM–VDAC assemblies (0.753–0.801), whereas performance was more variable for ER translocons (0.650-0.687; **Extended Data Fig. 7**) and SCN5A C-terminal domain complexes (0.480-0.737; **Supplementary Fig. 15**). Together, these case studies demonstrate the applicability of CHORUS across biologically diverse membrane-protein assemblies.

### Proteome-scale landscape of residue interactions on membrane proteins

Ultimately, we capitalized on the CHORUS predictions to generate an atlas of interaction interfaces across membrane proteins to bridge the gap between sparse 3D protein complexes and the vast expanse of uncharacterized membrane interactomes (**Fig. 7**; **Supplementary Table 21**). It spans 80,619 expert-curated protein entries deposited in UniProt corresponding to 70,247 non-redundant sequences, 4,918 species across all domains of life, and over 33.7 million residues (**Fig. 7a**). Quantitative mapping across the taxonomic cladogram uncovered substantial historical sampling asymmetries across evolutionary lineages (**Fig. 7a,b**). The pronounced taxonomic imbalance (Gini coefficient: 0.865) revealed that membrane-protein records were disproportionately concentrated in a small number of well-characterized species, highlighting substantial gaps in proteome coverage across the tree of life. CHORUS achieved near-complete membrane proteome-wide prediction coverage, with 99.21% of sequences processed directly and the remaining long and ultra-long proteins accommodated through overlapping windows (**Fig. 7c**). Predicted interface densities varied with sequence length, with most proteins exhibiting 15-40% interacting residues. To facilitate the exploration, we packaged this pan-proteomic landscape into an open-access web resource (**CHORUS Atlas**, https://chorus.atlas.sunnylab.org), which seamlessly integrates cross-species accession indexing, per-residue interaction probability profiles, and automatic topological alignment onto experimental structures or homology models, e.g., PDB 8V00 for olfactory receptor coreceptors (**Fig. 7d**). To assess the functional coherence of this proteome-wide catalog against orthogonal biochemical evidence, we systematically cross-referenced predictions against BioLiP ligand-binding annotations and CATH FunFam functional conservation scores, establishing that predicted inter-chain contact hot-spots systematically correlate with known catalytic and evolutionary centers.

**Fig. 7.**
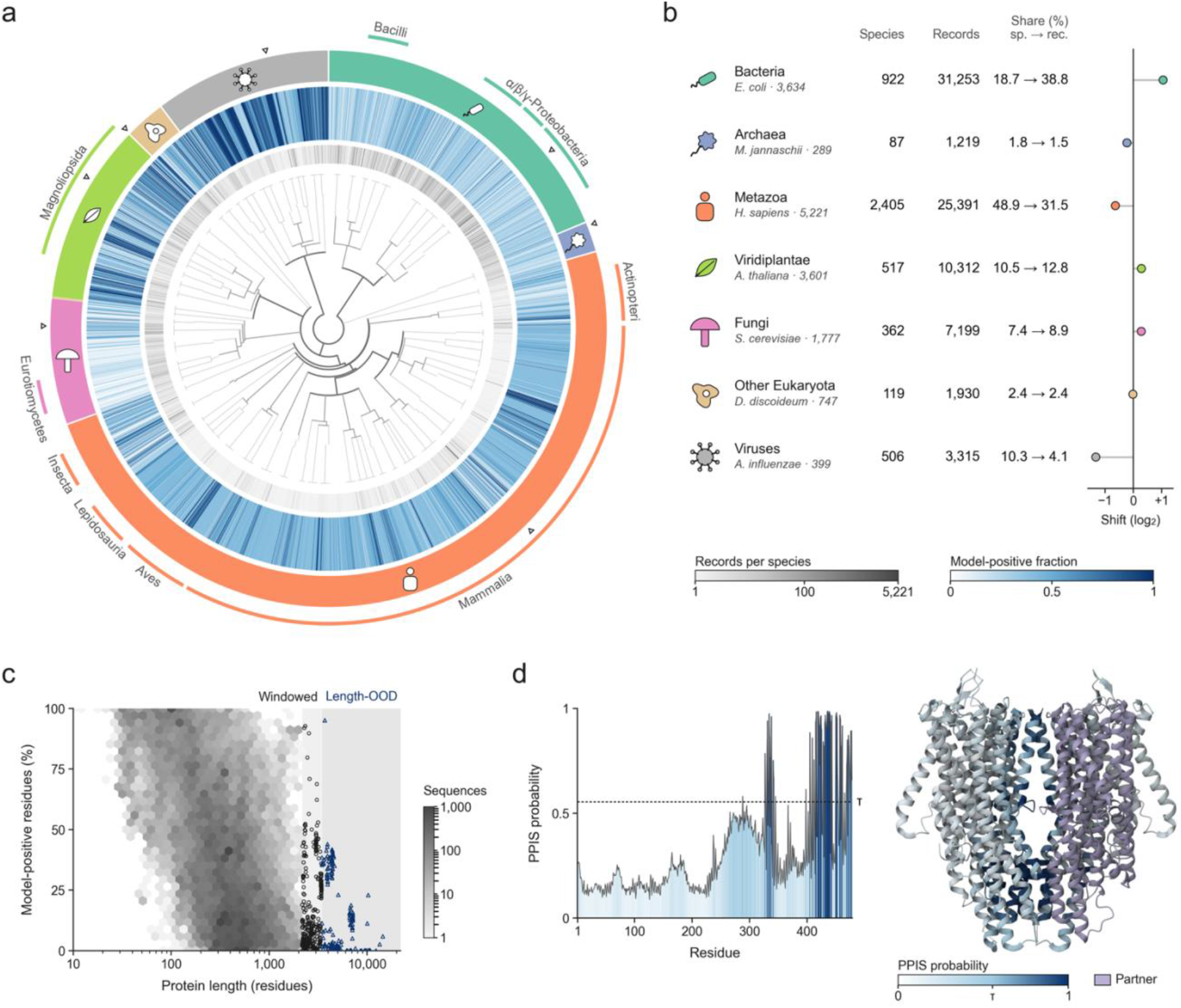
Atlas of CHORUS-predicted interaction interfaces in the membrane proteome. **a,** Global taxonomic view across 4,918 species. The cladogram represents 80,619 expert-reviewed UniProt membrane proteins resolving into 70,247 unique sequences and 33.7M residues. Radial clades depict non-hidden lineages with ≥ 25 species. From the inside out, the concentric rings display the volume of protein records per species (grey, log_10_ scale), the fraction of CHORUS-predicted interacting residues, and the broad lineage group (colored band, marked with its glyph). Open triangles mark the record-richest species in each lineage. Outer arcs delineate dominant clades (≥ 50 species), with classes ≥100 species labelled. α-, β- and γ-Proteobacteria share one label. **b,** Composition of each broad lineage showing species and record counts, landmark species with their record counts, and the sampling shift, log_2_ (record share/species share). Sampling is concentrated, with 2,537 of 4,918 species (51.6%) being singletons, the top-ranked 10 and top-ranked 100 species contributing 33.4% and 61.0% of records, and the species record-count Gini coefficient being 0.865. **c,** Protein length plotted against the fraction of CHORUS-predicted interacting residues per protein at the validation-selected threshold *p* ≥ 0.554. Proteins inferred at exact length (69,694; 99.2%) are shown as a log-scaled hexbin density, while proteins inferred with overlapping windows (368; 0.5%) are open circles. Length-OOD proteins (185; 0.3%) are triangles and use overlapping windows. The three groups are mutually exclusive. **d,** Example query for the odorant receptor coreceptor Orco (A0A026W182, Ooceraea biroi). Left: predicted interaction probabilities, with the dashed line at *τ* = 0.554 and 12.3% of 478 residues classified as interacting. Right: the probabilities projected onto its 3D structure (PDB code: 8V00), a homolog with 77.1% sequence identity (469 of 478 residues mapped), as rendered by the CHORUS Atlas. Partner chains and unmapped residues are shown tan and grey, respectively. Panel **d** and the outer ring of panel **a** share the atlas probability spectrum, namely, white below *τ*, slate at *τ*, and navy at 1.

### CHORUS-predicted interaction interfaces are independent of protein-ligand binding sites

We mapped experimentally curated small-molecule and ion-binding sites from BioLiP onto CHORUS-predicted interaction sites in membrane proteins. This mapping identified a total of 100,003 protein-ligand-binding (PLB) residues across 2,994 proteins. We then compared the observed overlap between CHORUS-predicted interacting residues and PLB residues with that expected using a sequence-specific model (see ***Methods*** for details) (**Supplementary Fig. 16a-c**). The co-occurrence analysis identified 19.1% of PLB residues falling within predicted interaction sites, closely matching the expected overlap (observed-to-expected ratio, O/E = 0.979). This was consistently observated across cytoplasmic (0.967), extracellular/luminal (1.019) and transmembrane (0.983) regions, whereas intramembrane sites showed only a modest positive shift (1.094) (**Supplementary Fig. 16c**). The observed-minus-expected differences were likewise centered close to zero at the sequence level (**Supplementary Fig. 16d**). The comparison samples only ligand contacts captured in experimentally resolved structures and should, in principle, represent a restricted view of the accessible ligand-binding landscape. Indeed, residues lacking BioLiP annotations constitute an annotation background rather than a verified non-binding class. Thus, our analysis establishes that the lack of systematic overlap between BioLiP-annotated ligand contacts and CHORUS-predicted PPI interfaces does not preclude small-molecule binding or modulation at individual interfaces.

We next assessed whether this relationship was consistent across individual proteins and evolutionary lineages by quantifying the difference in PLB-site occupancy between our predicted interacting and non-interacting residues. Values near zero therefore indicate comparable PLB-site occupancy in the two residue classes. Differential PLB-site occupancy remained centred close to zero across major taxonomic strata (**Supplementary Fig. 16e**) and representative species (**Supplementary Fig. 16f**), with no consistent directional shift towards either predicted PPI or non-PPI regions. Together, these analyses show that CHORUS-predicted interfaces do not preferentially recapitulate known PLB sites, providing a cross-functional assessment of model specificity and indicating that CHORUS may capture residue-level information complementary to established ligand-binding annotations.

### Association with FunFam conservation

To determine whether CHORUS-predicted interaction sites preferentially occupy evolutionarily constrained positions, we mapped predictions onto CATH FunFam conservation profiles (**Fig. 8**). The high-confidence analysis encompassed 2,739,713 Scorecons-mapped residues from 9,529 exact sequences distributed across 511 FunFams. Residues with Scorecons ≥0.7 were designated as highly conserved according to the established FunFam criterion^69^, and their overlap with predicted PPI sites was compared with the corresponding expectation, which thus account for both the prevalence of conserved residues and the fraction of residues classified as interaction sites within each sequence (**Fig. 8a-c**). This analysis identified that 23.0% of highly conserved residues coincided with predicted PPI sites, corresponding to an observed-to-expected ratio of 1.014 and indicating only limited overall enrichment. The association varied across membrane topologies. The strongest enrichment was observed in cytoplasmic regions, with an observed-to-expected ratio of 1.127, whereas extracellular, transmembrane, and intramembrane regions showed ratios of 1.013, 0.980, and 1.009, respectively (**Fig. 8b,c**). Thus, our predicted protein interaction sites were enriched modestly for highly conserved residues overall, with most evident in cytoplasmic regions. We therefore complemented this threshold-based analysis by treating Scorecons as a continuous measure of evolutionary conservation. Within each protein, mean Scorecons was compared between predicted PPI sites and non-PPI sites (**Fig. 8d**). Predicted PPI sites showed higher mean conservation within cytoplasmic and transmembrane regions, whereas no clear difference was observed in extracellular regions. Notably, the positive conservation shift within transmembrane regions was detectable in the continuous analysis despite the absence of enrichment under the binary Scorecons ≥ 0.7 criterion, indicating that the association with evolutionary constraint was not restricted to the most highly conserved residues.

**Fig. 8.**
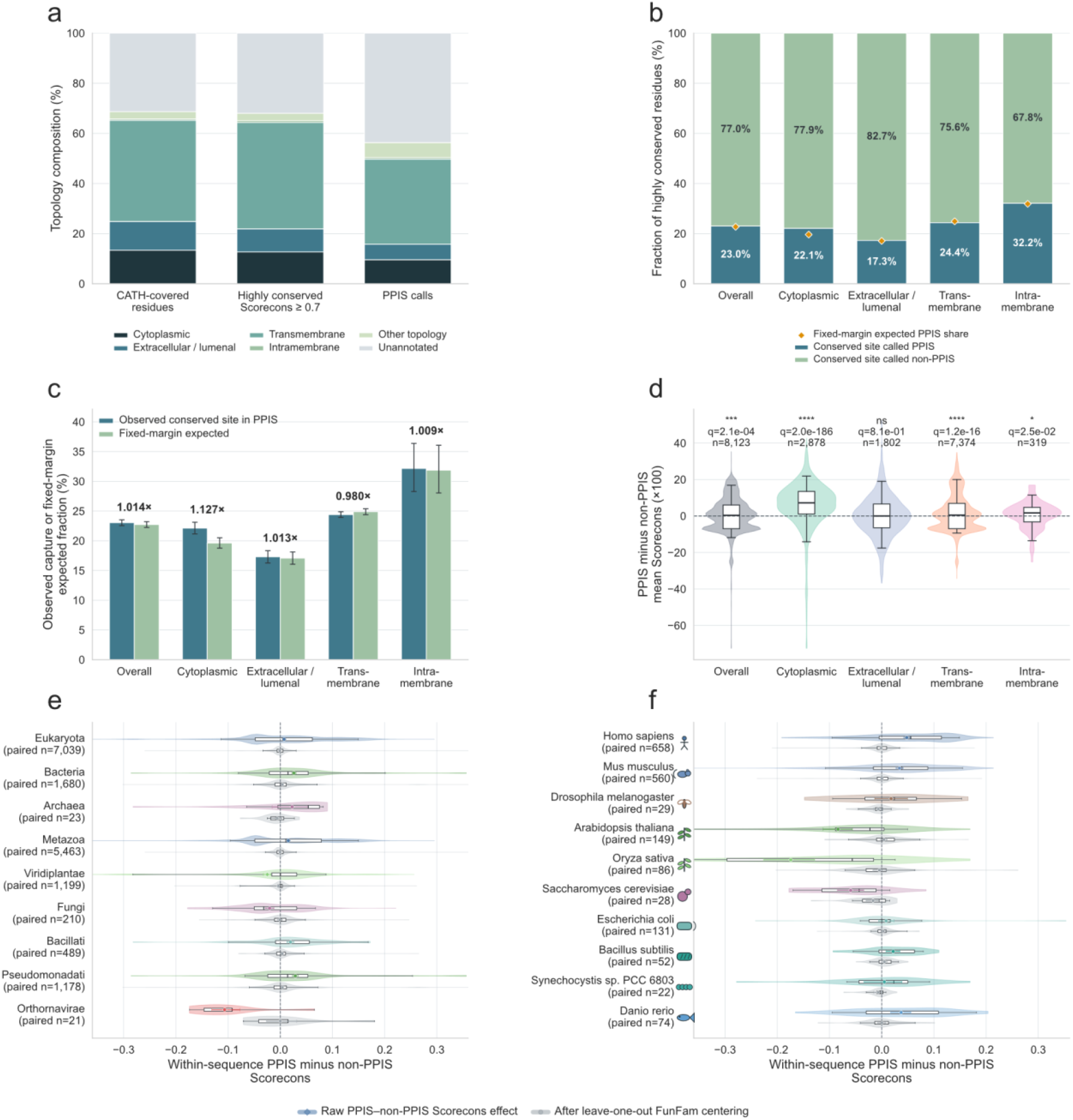
Relationship between predictions from the family/depth-controlled cross-pLM ensemble and CATH FunFam conservation. **a,** Topological composition of CATH-annotated residues, evolutionarily conserved residues, and predicted protein-protein interaction sites (PPIS). **b,** Fraction of conserved residues overlapped with PPIS. **c,** Fraction of observed and fixed-margin expected conserved residues in PPIS across full-length (overall), cytoplasmic, extracellular, transmembrane, and intramembrane regions. **d,** Distribution of Scorecons between predicted PPIS and non-PPIS residues. **e,f,** Distribution of Scorecons across taxonomic strata (**e**) and representative species (**f**).

The magnitude of these within-sequence conservation differences varied across taxonomic groups (**Fig. 8e**) and representative species (**Fig. 8f**) and was generally attenuated after leave-one-out FunFam centering, suggesting that part of the apparent taxonomic variation reflected differences in the FunFam composition represented across taxa. Collectively, these analyses indicate a modest but reproducible association between predicted PPI sites and evolutionary conservation, with the clearest signal observed in cytoplasmic regions and additional evidence of elevated conservation within transmembrane PPI sites.

## Discussion

In this work, complementary representations from diverse pLMs have been harnessed to predict interaction interfaces on membrane proteins directly from sequences without requiring structural coordinates, MSAs, or nominated interacting partners. We carried out a systematic benchmark for the optimal choice of pLM layers across multiple biological properties specific to membrane proteins, prioritizing the use of deeper layers to decipher residue-level interaction patterns. We applied CHORUS to generate a membrane proteome-scale atlas of interaction sites, which we have used to uncover a largely unexplored landscape distinct from known ligand-binding annotations and shaped by different evolutionary constraints across membrane environments.

However, several limitations of CHORUS warrant consideration. First, the knowledge about physical interactions used to train CHORUS was gained from known experimentally resolved protein assemblies, which represent a fraction of the membrane interactome. This may give rise to an issue that residues without experimentally observed contacts cannot necessarily be regarded as non-interacting, particularly for transient interactions. Second, the current version of CHORUS does not identify specific interaction partners or resolve the geometry, affinity, or conformational dynamics of protein-protein interactions. Third, the use of predictions from the CHORUS atlas necessitates further experimental validation, particularly through site-directed mutagenesis and protein interaction assays to further elucidate their underlying molecular mechanisms.

Together, CHORUS establishes a scalable, sequence-only framework for exploring membrane protein interaction interfaces at proteome scale. By extending interface predictions across 80,619 membrane proteins from 4,918 species, this work provides a foundation for systematic investigation of membrane protein interactions, functional characterization, and future therapeutic exploration.

## Methods

### Sequence-only prediction across a family-layer-backbone candidate space

To capture local biochemical surroundings and long-range structural dependencies from frozen pLM representations, we engineered three hybrid backbone architectures for residue-level interface prediction, including Gated Conv-BiGRU (GCB), Conv-Transformer (CT), and Gated Transformer-BiGRU (GTB) (**Fig. 2a and 2b**). Across all architectures, parallel multi-scale 1D convolutions extract localized motif patterns, which are integrated with contextual representations derived from multi-head self-attention or bidirectional recurrent units. In GCB and GTB, dynamic gating mechanisms adaptively balance local convolutional signals against sequential contexts. GCB incorporates ten auxiliary residue-level biophysical priors, including amino-acid (AA) index^70^, sequence length, noncanonical residue indicator, selenocysteine (U) indicator, polarity, hydrophobicity, charge, hydropathy, aromaticity, and relative position, while CT and GTB operate solely on foundation embeddings 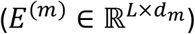.

Different pLM families and transformer layers capture distinct hierarchies of protein structures, from local secondary-structure motifs to global fold topologies. To identify optimal representational depths, we systematically sampled 19 layers from pLMs, with 7 for ESM-2 (650M), 7 for ESM-C (600M), and 5 for ProtT5-XL-U50. Pairing each layer with the three candidate backbones led to a comprehensive pool of 57 prediction architectures (**Fig. 2a,c**). Candidates within each pLM family were ranked by chain-macro average precision (AP) values across proteins from the validation dataset. Rather than relying on a single architecture, two top-performing models per pLM family were selected to construct the final ensemble model, dubbed Cross-pLM-6 (**Fig. 2c,d**).

The resultant ensemble model strategically traverses structural depths and architectural archetypes, including two configurations utilizing ESM-2 layer 30 (coupled with GTB and CT), two exploiting ESM-C layer 30 (coupled with GTB and GCB), and the remaining two deploying intermediate representations from ProtT5 layers 18 and 24 (paired with GTB and GCB, respectively, **Fig. 2d**). Finally, individual residue interaction probabilities are pooled via unweighted averaging across the six complementary predictors. Binary interface calls were assigned using a decision threshold determined exclusively on the validation set and subsequently fixed for all test evaluations (*τ*_val_ = 0.554). This design allows the final classification of interfacial residues to benefit from complementary representational perspectives, thereby reducing single-model bias.

### Model selection

#### Random search with exchange refinement

To investigate whether alternative combinations of individual models could improve ensemble performance, we implemented a random search followed by iterative exchange refinement, drawing on established stochastic optimization and ensemble-selection strategies^71^. For each of three independent random seeds, 299 combinations of six individual models were randomly sampled from the pool of 57 candidate predictors and evaluated using the chain-macro AP metric on the validation data. Ensemble predictions were obtained by averaging the residue-level interaction probabilities across the six individual models. For each seed, the combination achieving the highest validation chain-macro AP value was subjected to iterative refinement, whereby one selected individual model was replaced with an unselected candidate. Each exchange was retained only if it improved the chain-macro AP metric, and the procedure continued until no further improvement could be achieved through individual-model replacement. The resulting combinations were compared with the original CHORUS ensemble to assess whether random exploration followed by local optimization could yield improved predictive performance (**Extended Data Fig. 1a**).

#### Cluster-robust single exchange

To examine whether the CHORUS ensemble could be improved through individual-model replacement, we implemented a cluster-robust exchange strategy based on the maximin principle of robust optimization^72^. Starting from the original CHORUS ensemble comprising six individual models, we systematically evaluated all 306 possible exchanges, each replacing one selected model with one of the remaining 51 candidates. For each exchange, predictive improvement relative to CHORUS was assessed across three cluster-weighting schemes and additional perturbations involving the exclusion of individual clusters or cluster groups. The minimum AP gain across these conditions was defined as the worst-case gain, and the exchange maximizing this criterion was selected to prioritize improvements that remained robust to changes in cluster composition and weighting. Among the 306 exchanges, only 15 yielded positive worst-case gains. The highest worst-case gain was achieved by replacing the ESM-2 layer-30 CT model with the ESM-C layer-36 GTB model, while retaining the other five individual models (**Extended Data Fig. 1b**).

#### Restricted exhaustive search

To investigate whether systematic optimization of ensemble composition could improve predictive performance, we implemented a two-stage restricted exhaustive search, following established principles of performance-guided ensemble selection^73^. Given the computational complexity of evaluating all 36,288,252 possible combinations (i.e., 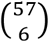 of six individual models from the 57 candidate predictors, we first reduced the search space through pLM-family-stratified screening. Within each of the three pLM families (ESM-2, ESM-C and ProtT5), we retained the two individual models achieving the highest chain-macro AP value on the validation data, together with the four highest-ranked remaining candidates according to cluster-macro AP values, yielding a pool of 18 individual models. We subsequently exhaustively evaluated all 18,564 possible combinations (i.e., 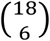 of six individual models from this pool. For each combination, residue-level interaction probabilities were aggregated by unweighted averaging, and ensemble performance was assessed using the cluster-averaged AP to assign equal weight to individual protein clusters irrespective of their respective chain counts. The combination achieving the highest validation cluster-averaged AP was retained for subsequent comparison with the original CHORUS ensemble (**Extended Data Fig. 1**).

### Dataset construction and structural quality control

To develop and evaluate CHORUS, transmembrane protein complexes were retrieved from the protein data bank of transmembrane proteins (PDBTM)^74^, which provides an exhaustive cohort of annotations of protein locations (e.g., transmembrane segments). We restricted the dataset to multimeric assemblies with at least one interacting partner, which yielded an initial pool of 45,750 chains across 11,469 entries.

To resolve discrepancies with unresolved regions in canonical records, structure-derived sequences were aligned against canonical PDBTM sequences using the Python built-in class, difflib.SequenceMatcher, with heuristics disabled. Chains were excluded if structural sequence coverage fell below 70% of the canonical record, or if sequence identity was < 99% with more than 3 mismatches. Entries with empty sequences, index corruptions, or coordinate-record inconsistencies were discarded. This filter eliminated nine chains, ensuring strict one-to-one mapping between sequence positions and coordinate-derived labels. In total, the final dataset comprises 40,876 chains across 8,363 entries, encompassing 11,482 distinct structure-derived sequences and 11,537,440 residue-level contact labels. Based on PDBTM classifications, these correspond to 40,849 *α*-helical and 27 *β*-strand target chains.

### Definition of interacting residues

Interacting residues between protein chains were identified using spatial proximity between their respective heavy atoms, following distance-based criteria established for interaction interfaces in membrane proteins^48,49^. For each multimeric complex, the target chain was evaluated against all co-assembled partner chains, which were not restricted to transmembrane proteins. All hydrogen atoms, crystallographic waters, and heteroatoms (HETATM records) were removed. For each target residue at sequence position *i*, the minimum Euclidean distance across all heavy-atom pairs belonging to any partner chain was denoted as *d*_*i*_. In accordance with conventional interface definitions (4-6Å), a residue was designated as an interacting residue (*y*_*i*_ = 1) if it fell within 5Å of any partner chain, and non-interacting (*y*_*i*_ = 0) otherwise:

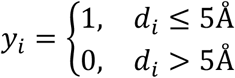

Residues lacking resolved coordinates in the structure were excluded from the analysis rather than treated as non-interacting. Crucially, designating a residue as interacting denotes geometric contact within a specific structural snapshot rather than a direct thermodynamic contribution to binding affinity, whereas a non-interacting assignment reflects the absence of contact within that observed conformation. This criterion was applied uniformly across training, validation, and test sets.

### Sequence redundancy control and dataset partitioning

To avoid sampling bias and further ensure a fair comparison for evaluating generalization ability, we eliminated redundancy between proteins both within and across datasets. Pairwise sequence identity was computed across all sequence pairs via Needleman-Wunsch global alignment using the BLOSUM62 matrix^75^ with affine gap penalties (gap opening set as -10 and gap extension set as -0.5). Sequence identity *SI*(*A,B*) between sequences *A* and *B* of lengths *L*_*A*_ and *L*_*B*_ was defined over the longer sequence to avoid inflating local alignment fragments, such that

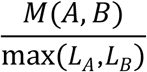

Sequences were organized into an undirected graph, connected by edges if they shared ≥ 30% sequence identity (in line with established studies^49,76^) or co-assembled within the same PDB entry. Resolving the connected components of this graph yielded 419 strictly distinct clusters, which were then allocated, with a deterministic greedy approach that balance chain contexts, residue and interacting-residue counts, to training, validation, and testing splits in an approximate 70:15:15 ratio, respectively. The resulting splits contain 299 clusters (30,675 chain contexts) for training, 65 clusters (5,546 contexts) for validation, and 55 clusters (4,655 contexts) for testing, respectively.

### External datasets and evaluation

CHORUS was evaluated externally in two settings: *i*) direct inference using the released six-member ensemble and *ii*) a separate retraining experiment following sequence-similarity exclusion. We collected 3 datasets specialized for membrane proteins, including CompData (101 chains)^68^, TestData (36 chains)^48^, and IndepData (30 chains)^49^, alongside 15 general-protein datasets or docking views from literature^34,35,58,77–83^ (**Supplementary Table 10**). Each external benchmark retained its native interaction criteria and residue masks. For membrane protein targets, native 6Å heavy-atom criteria were maintained. For unbound docking targets, bound interfaces (≤ 6Å) were mapped via sequence alignment. For proteins used in MaSIF, residues were defined as interacting if their interface areas exceeded 5Å^2^ and > 4% of total surface area (14,227 evaluated, 2,720 masked residues).

Predictions were generated by unweighted probability averaging across the six ensemble backbones and scored by chain-macro and pooled AP/AUROC values. Overlapping cohorts were evaluated individually. Since the collected external membrane protein targets share substantial sequence identity with those compiled in our datasets, evaluating the released ensemble model of CHORUS serves as a reference benchmark. In the separate retraining benchmark, any chain from our raw training dataset sharing ≥ 30% identity with the 167 external membrane chains was excluded prior to model fitting, which resulted in removal of 16,050 training and 1,434 validation contexts and retaining 14,625 and 4,112). Backbones were retrained on frozen pLM embeddings (seed 0) and evaluated against native external definitions.

### Membrane protein-specific properties

#### Helix connectivity

Numerous transmembrane proteins adopt conformations in which transmembrane helices are embedded within densely packed multi-helix bundles^84^. Local packing around transmembrane helices serves as a primary structural constraint shaping their tertiary structures^85^. We used connectivity between helices as a structural probe to determine whether 3D packing environments have been encoded by frozen pLM representations rather than merely local physicochemical properties.

The helix-neighbor connectivity was defined as a helix-level descriptor of intrachain transmembrane packing. Within a target chain, each transmembrane helix annotated in PDBTM was represented as a node in a graph where an edge can be represented as two helices being in contact. Residues belonging to distinct helices were considered to be in contact when their minimum heavy-atom distance was ≤5.5 Å as reported by Fuchs et al.^55^. A helix-helix edge was retained only when at least three residues from each of the two helices participated in such inter-helical contacts. This requirement suppresses edges arising from isolated atomic proximities, and, instead, captures sustained helix-helix packing. Helix connectivity was quantified by node degree and grouped into three packing states: isolated (0), paired (1), or bundled (≥ 2 neighboring helices). Residue embeddings were mean-pooled across each helix to yield a single representation, which was then evaluated for its predictive capacity over these connectivity classes. This task, therefore, tested whether frozen sequence-derived representations encoded higher-order transmembrane architecture sufficient to distinguish relatively isolated helices from helices participating in pairwise or multi-helix packing arrangements.

#### Transmembrane location

The lipid bilayer imposes stringent physicochemical constraints on α-helical transmembrane proteins, which require membrane-spanning segments to partition into a highly hydrophobic and anisotropic environment^86^. Residues buried within the bilayer experience a different physicochemical environment from residues located in membrane-flanking regions, and this distinction is reflected most likely in amino-acid composition, local sequence context, and helix organization. We therefore used transmembrane location as a structural probe to test whether frozen pLM representations encode the membrane-embedded state of individual residues. Let *T* denote the retained interior residues of all annotated helical segments and *S* the union of residues assigned to Side 1 or Side 2 as defined by PDBTM. The binary target was

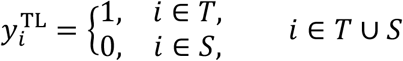

The first and last two residues of each helical segment were excluded to reduce boundary ambiguity, the two side classes were pooled, and unannotated residues were omitted.

#### Helix position

The intra-helical residue position is informative since amino-acid preferences and insertion energetics vary systematically across the membrane^87^. Residues near the center of a membrane-spanning segment occupy a different structural context from residues approaching the helix boundaries, where the polypeptide transitions toward membrane-flanking regions. We therefore used helix position as a residue-level probe to test whether frozen pLM representations encode this positional organization within annotated transmembrane segments.

For a PDBTM-annotated helical segment extending from residue *a* to *b*, the normalized position of residue *i* was defined as

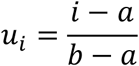

Residues satisfying 0.25 ≤*u*_*i*_ ≤ 0.75 were assigned to the helix-center class, whereas the remaining residues within the same helical segment were assigned to the helix-boundary class, given below

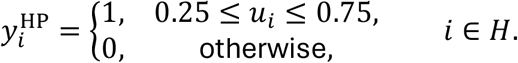

Thus, this task sets out to understand whether pLM representations can distinguish the central portion of a transmembrane helix from its terminal regions, independently of whether the helix lies on the cytoplasmic or extracellular side of the membrane.

#### Membrane depth

Membrane depth provides a direct geometric measure of where a residue is positioned across the bilayer and therefore complements sequence-based descriptors of transmembrane topology. Whereas helix position describes a residue relative to the linear boundaries of an annotated transmembrane segment, membrane depth captures its physical placement within a membrane-oriented 3D structure. We therefore used it to test whether frozen pLM representations encode information about the spatial organization of residues across the bilayer, beyond what can be inferred from sequence position alone. PDBTM membrane-oriented coordinates were used for this purpose, with the membrane midplane defined by *z* = 0.

For an eligible residue *i* within a PDBTM H-labelled transmembrane segment, we first calculated the mean *z*-coordinate of its non-hydrogen atoms,

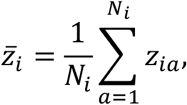

where *N*_*i*_ is the number of non-hydrogen atoms represented for residue *i*. Its membrane depth was then defined as the absolute distance from the membrane midplane,

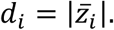

Residues were assigned to three depth classes,

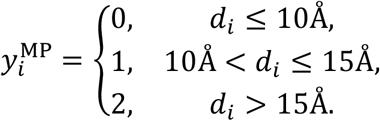

Note that using the absolute *z*-coordinate makes membrane depth a measure of distance from the bilayer midplane, independent of which side of the membrane the residue occupies.

### Learning architecture

CHORUS predicts residue-level interactions directly from amino-acid sequences without structural inputs or evolutionary alignments. Feature vectors were extracted from frozen pLMs, including ESM-2^51^, ESM-C^52^, and ProtT5^53^ [12, 19, 20], sampling 19 distinct hidden layers from pre-Transformer inputs (Layer 0) to final layers. Non-canonical symbols (U, Z, O, B) were unified as X, and sequence delimiters were stripped. Protein sequences exceeding maximum context lengths were processed via sliding windows (window size: 1,022 residues yielding 128-residue overlaps for ESM-2/ESM-C, 1,024 yielding a 128-residue overlap for ProtT5), followed by linear averaging across intersecting segments. The pLMs and chosen layers are summarized in **Table 1**.

**Table 1.**
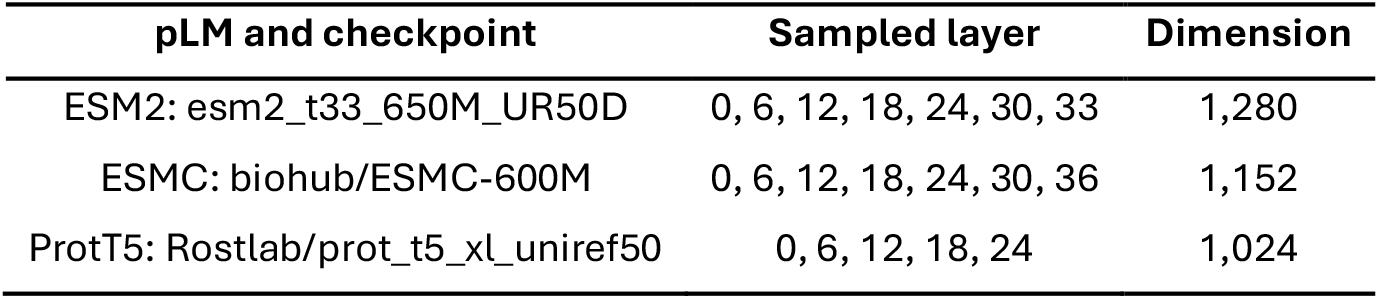
Summary of pLMs and sampled layers.

We also calculated ten features directly from the input sequence, including a numerical amino-acid identity code, labels identifying noncanonical or unknown residues and selenocysteine (U), hydropathy, a coarse charge value, labels for aromatic, polar and hydrophobic residues, sequence length, and normalized residue position. The charge property was encoded as +1 for *K, R* and *H*, -1 for *D* and *E*, and 0, otherwise. For residue position *i* in a sequence of length *L*, normalized position was 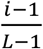, ranging from 0 at the first residue to 1 at the last. The GCB backbone received these ten features in addition to the pLM embeddings, while the CT and GTB backbones received pLM embeddings alone. All ten additional features were derived from sequence without structural or alignment information. Evaluating all combinations of the 19 representation layers and three backbones established a structured space of 57 candidate predictors (19 × 3).

Extracted representations were mapped to interaction probabilities using **additional neural networks.** To systematically evaluate the representational capacity across pLM families and layers, we designed three complementary backbones of neural networks as follows.

- **Gated Conv-BiGRU (GCB)**: Parallel 1D convolutions (kernel sizes *k* ∈ {3,9}) capture sequentially contextual features. Concatenating the projected input with both convolutional representations feeds a gating sub-network that computes learned interpolation weights between 0 and 1, which blends local convolutions with the original projection. A single bidirectional GRU (BiGRU)^88^ processes this blended sequence, and its hidden states are concatenated with the gated features before classification.
- **Conv-Transformer (CT)**: Parallel convolutions (*k* ∈ {3,7,11}) capture multi-scale local neighborhoods, which are concatenated with the projected features and linearly compressed to the backbone width. Two Transformer encoder layers^51^ model long-range residue dependencies. The classifier receives the concatenation of the Transformer output and the initial projection, preserving both localized and global context.
- **Gated Transformer-BiGRU (GTB)**: Two initial Transformer encoder layers process the projected features, followed by parallel convolutions (*k* ∈ {3,9}) that inform a gating mechanism. This gate balances the Transformer output against the input projection. A single BiGRU processes the resulting mixture, with final classification performed on the concatenated BiGRU and mixed features.

Input features for each residue were standardized via layer normalization^89^, projected to the backbone hidden dimension (**Table 2**), and activated by Gaussian error linear units (GELUs)^90^. For GCB, the ten engineered sequence features were concatenated with the pLM embedding prior to normalization. A linear projection produced an unconstrained logit *z*_*i*_ for residue *i*, mapped to an interaction probability via the standard sigmoid function (1 + exp(−*z*_*i*_))^−1^ . Backbones were optimized jointly while pLM weights remained frozen.

**Table 2.** Settings of downstream neural network backbones.

| **Setting** | **Gated Conv-BiGRU** | **Conv-Transformer** | **Gated Transformer-BiGRU** |
|---|---|---|---|
| Projection width | 256 | 320 | 256 |
| GRU units per direction | 256 | - | 384 |
| Transformer layers/attention heads | - | 2/5 | 2/4 |
| Transformer feedforward width | - | 1,280 | 1,024 |
| Learning rate | 10<sup>-3</sup> | 2.5×10<sup>-4</sup> | 3×10<sup>-4</sup> |
| Residue-pair ranking weight | 0 | 0.05 | 0.03 |

### Training objectives and loss formulation

Models were supervised against binary interaction states, where *y*_*i*_ = 1 denotes an interacting residue and *y*_*i*_ = 0 denotes a non-interacting residue. To mitigate the class imbalance problem^91^, i.e., under-representative class for interfacial residues, positive examples were weighted by

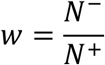

where *N*^−^ and *N*^+^ denote total non-interacting and interacting residues in the training set, respectively. The per-residue weighted cross-entropy *b*_*i*_ was defined as:

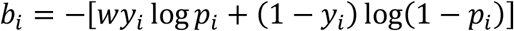

For GCB and GTB, we additionally followed the principles of focal loss^92^ to suppress the loss contribution from confidently predicted non-interacting positions while leaving interacting residues unscaled. Across a batch of *n* resolved sequence positions, the focal objective was computed as

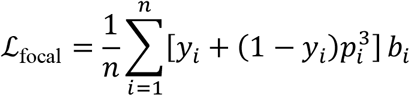

For CT, we used unmodulated weighted cross-entropy averaged at the chain-context level to prevent longer protein chains from dominating the gradient update. For a batch of *C* chain contexts with respective lengths *L*_*c*_:

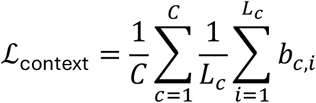

To enhance the relative scoring contrast of interacting sites within each protein sequence, CT and GTB were additionally trained with an intra-chain ranking loss. For an interacting residue *i* and a non-interacting residue *j* belonging to the same chain context, the pairwise margin loss was log(1 + exp(*z*_*j*_ −*z*_*i*_)). For contexts exhibiting both classes, *m* pairs (min(192,*N*_pos_ ×*N*_neg_)) were sampled with replacement, that is, interacting residues were sampled uniformly, whereas non-interacting residues were sampled from the hardest negatives (the top ⌊*m*⌋ highest scoring non-interacting residues). The resulting mean ranking loss ℒ_rank_ yielded the overall objective, given by

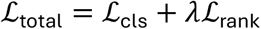

where ℒ_cls_ corresponds to ℒ_context_ for CT and ℒ_focal_ for the two gated architectures, with *λ* configured according to the hyperparameter table.

### Optimization and checkpoint selection

Trainable parameters were optimized using AdamW^93^, with a weight decay of 10^−4^, a constant learning rate, a dropout proportion of 0.15, and gradient norm clipping of 5. Protein chains used for training were dynamically batched and sorted by length (maximum 12 contexts per batch), which restricted total padded tokens to ≤ 14,000 during training and ≤ 18,000 during validation. Zero-padded positions were masked out of attention operations and excluded from all loss and metric evaluations. Training proceeded for up to 16 epochs with an initial random seed of 0. Model checkpoints were monitored after each epoch using the chain-macro-AP metric on the validation split, and training was terminated after four consecutive epochs without performance gain. The checkpoint achieving peak validation chain-macro-AP value was retained for ensemble integration and threshold selection.

### Ensemble construction and inference

CHORUS aggregates six trained models into an ensemble, selected by ranking all 57 candidates within each pLM family by validation chain-macro-AP values. This retains the top two per family at their optimal checkpoints, that is, layer 30 with GTB and CT for ESM-2, layer 30 with GTB and GCB for ESM-C, and layer 24 with GTB and layer 18 with GCB for ProtT5. A separate ablation ensemble (*primary_family_depth_x6*) obtained by fixing the GTB backbone across paired depths (ESM-2: 24/30, ESM-C: 24/30, and ProtT5: 18/24) was used for BioLiP- and FunFam-related analysis. The final predicted interaction probability *p*_*i*_ was derived by unweighted probability averaging across six selected models, expressed as

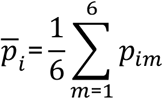

where *p*_*im*_ denote the probability assigned to residue *i* by individual model *m*.

### Metrics for performance assessment

Model performance was comprehensively evaluated based on a variety of metrics (see **Supplementary Note 2**). For residue *i*, let *p*_*i*_ ∈ [0,1] denote its predicted interaction probability and *y*_*i*_ ∈ {0,1} its reference interaction state (1 for interacting, 0 for non-interacting). Metrics were restricted strictly to structurally resolved and evaluated positions. The predicted probability *p*_*i*_ were converted to an interaction state based on a threshold *t*:

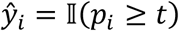

where I(·) equals 1 when its condition is satisfied and 0 otherwise. Precision (*P*), recall (*R*), and interfacial *F*_1_ (*F*_1,+_) are defined as

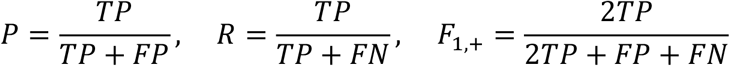

where *TP* stands for true positives, *FP* stands for false positives, and *FN* stands for false negatives. To account for class imbalance without bias toward majority non-interfacial residues, we also evaluated class-macro *F*_1_ and the Matthews correlation coefficient (MCC), such that

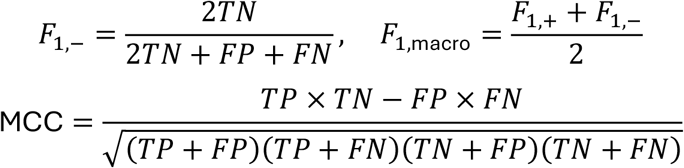

where *TN* stands for true negatives.

The threshold in support of residue interactions was calibrated independently for each model configuration by maximizing interfacial *F*_1_ across all pooled validation residues, resolved by selecting the highest threshold *t*^∗^, denoted as

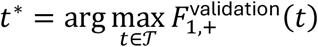

In this study, this procedure established *t*^∗^ = 0.5542 for the final ensemble model of CHORUS, which was held strictly fixed for test and external benchmarks without post-hoc adjustment.

In addition, two threshold-free metrics, the average precision (AP) and the area under the receiver operating characteristic curve (AUROC), are also used. For *J* distinct score cutoffs ordered descendingly, AP evaluates precision across incremental recall steps, calculated as

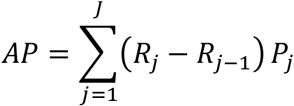

AUROC quantifies the probability that a randomly chosen interacting residue receives a higher predicted score than a non-interacting residue.

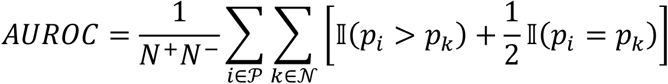

where *P* and *N* represent the sets of interacting (*N*^+^) and non-interacting (*N*^−^) residues, respectively. AP was evaluated for contexts containing ≥ 1 interacting residue, and AUROC required both classes.

To assess utility in targeted experimental mutagenesis, we evaluated top-*k* precision under both fixed (*k* ∈ {5,10,20}) and length-proportional budgets (Top-*L*/20, Top-*L*/10, Top-*L*/5, selecting *m*_*c*_ = max(1, ⌈*L*_*c*_/*d*⌉) residues for chain length *L*_*c*_). Precision within selected subset *S*_*c*_ was defined as

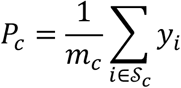

Probability calibration was measured using the standard Brier score^94^ and a class-balanced Brier score (*B*_balanced_)^95^ as follows.

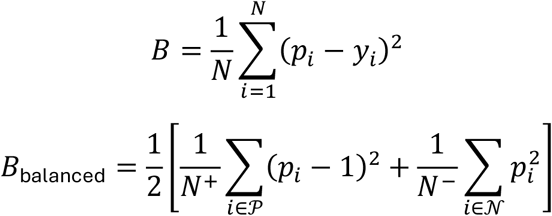

The primary benchmark endpoint was the chain-macro-AP metric (*AP*_*chain*_), which assigns equal weight to each eligible target-chain context

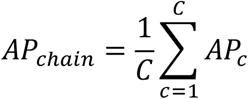

Complementary analyses examined pooled-residue metrics and cluster-level summaries. For sequence-similarity cluster *g* containing *C*_*g*_ eligible contexts, cluster performance *A*_*g*_ was computed as

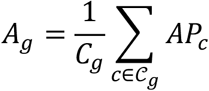

### Comparison of CHORUS with established methods

To benchmark CHORUS against established interface predictors, we evaluated published methods on the independent test partition (4,655 chain contexts, 2,186,751 residues) using unified reference interaction criteria. Evaluated frameworks comprised sequence-only models, evolutionary-profile/MSA-dependent methods, and structure-dependent architectures. Where official public checkpoints were available, these methods were deployed directly, including ScanNet^35^ (with and without MSAs), GraphPPIS (fast and evolutionary-profile-based), and five-fold averages of GPSite^60^, DeepProSite^66^, and Gated-GPS^63^. Models lacking standalone complex-prediction checkpoints or accessible pipelines were retrained on our training splits following published specifications, including PIPENN-EMB^47^ (embedding-based implementation), ALLSites^59^ (ESM-2-3B layer-36 embeddings with validation AP checkpointing), and DeepPPISP^39^ (re-implemented architecture). For DeepTMInter^49^, PDBTM-derived membrane topologies were supplied using Phobius predictions^96^.

### Predicted structural coordinates

For structure-dependent predictors, 3D structures were predicted *de novo* from their sequences using ESMFold^51^ (v1, chunk size 64) or ColabFold^77^ (AlphaFold2-ptm). Derived coordinates supplied information about structural features, solvent accessibility, and secondary structure elements without falling back to experimental coordinates upon structural failure. Predicted interacting or non-interacting residues were aligned to ground-truth labels based on chain contexts. Failed predictions were penalized via coverage metrics rather than imputed as zeros. The best performance of these structure-dependent methods, evaluated on ESMFold- or ColabFold-predicted structures, are reported.

### Representation ablations, ensemble controls, and error independence

To systematically investigate the determinants of model performance, we constructed three schemes for ablation studies as shown below.

1. **Exploratory architecture screening**: Initial tests on sets of ∼1,400 training and ∼300 validation chains were conducted for choosing three backbones from linear projections, MLPs, residual/dilated CNNs, U-Nets, BiGRUs, and Transformers.
2. **Ensemble diversity controls**: CHORUS was benchmarked against five controlled six-member variants: *i*) variation of random seeds from 0 to 5 using ESM-2 layer 30, *ii*) coordinate permutations, *iii*) within-family depths (ESM-2 layers 0, 6, 12, 18, 24, 30), *iv*) identical duplicates, and *v*) a single-model baseline scaled to match the total number of parameters of our ensemble model (27.7M parameters vs. ∼24.4M across CHORUS backbones).
3. **Synergy and prediction decorrelation**: To assess the complementarity of individual models, we evaluated all 15 pairwise model combinations and six leave-one-out ensembles. Pairwise Pearson correlations between individual model predictions were computed, relating lower correlation to performance gains over the reference ensemble.

### Representation, ensemble and length-sensitivity analyses

Test contexts were partitioned into four sequence-length groups (20-288, 289-424, 426-614, and 615-1,380) across CHORUS and 45 balanced candidates (i.e., 3 families × 5 shared layers × 3 backbones). Because empirical AP values covary with the baseline fraction of interacting residues, observed AP values were complemented by standardized AP values adjusted to a constant reference prevalence (*π*_0_ = 523,268/2,244,041 ≈ 0.233). For a context with local interacting fraction *π*_*c*_, the precision at given true-positive (TPR) and false-positive rates (FPR) was standardized as

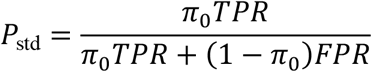

Moreover, to determine how the structure quality affects prediction performance, structure-dependent predictors for interacting residues were evaluated based on structures derived from PDBTM alongside those predicted by ESMFold and ColabFold.

### Comparison of candidate-selection and ensemble strategies

To evaluate alternatives to the validation-AP criterion used in CHORUS, we benchmarked ensemble strategies over an expanded candidate space of 63 predictors, comprising the 57 single-layer models and six multi-layer ProtT5 variants (concatenating layers 6-24 or 0-24 across the three backbones). We evaluated all 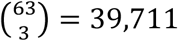 possible triplets from the 63-candidate pool to determine whether minimizing probability calibration error could identify compact, three-model ensembles without compromising ranking precision. The cluster-averaged Brier score (*B*_*S*_) was computed as

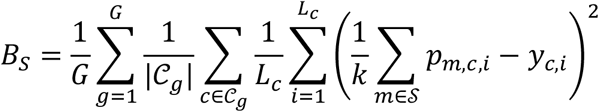

where *G* is the number of clusters, *C*_*g*_ denotes the chain contexts in cluster *g, L_c_* is sequence length, and *y*_*c,i*_ ∈ {0,1} represents the reference interaction state. Selection rules evaluated individual minimum Brier scores (with and without language-model family constraints) and exhaustive triplet optimization.

### Biological probing and representation-performance association

To assess whether the structural and evolutionary properties have been captured by frozen pLM embeddings, we trained linear probes following established representation-probing protocols. Evolutionary conservation was derived from MSAs using normalized Shannon entropy in consistency with our previous research^49^, such that,

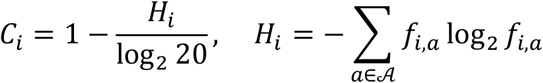

where *f*_*i,a*_ is the frequency of amino acid *a* at alignment position *i*. Probes were trained using class-weighted cross-entropy or mean squared error (AdamW^93^, 12 epochs) and scored by *R*^2^ or the chance-adjusted AP metric, given by

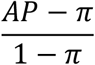

This adjustment rescales performance relative to the random expectation *π*, ensuring that probe accuracies can be directly compared across biological properties with widely differing positive frequencies. The independent-test chain-macro-AP gain (Δ*AP*) over layer 0 was measured against relative probe performance and network depth using ridge regression (regularization penalty 1). We compared depth-only, biology-only, and combined feature sets. To test generalizability across pLM families, regressions were fitted on two families and evaluated on the omitted third family, with feature standardization restricted strictly to the training families. Across the 16 held-out family-layer observations (observed gains *v*_*j*_, predictions 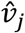, and mean gain 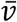), predictive accuracy was quantified by *R*^2^ and mean absolute error (MAE), given by

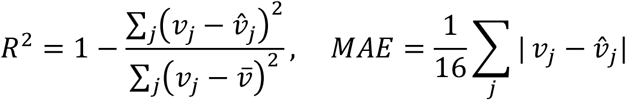

### Evaluation across membrane and biological contexts

To dissect performance across diverse biological environments, predictions from the final CHORUS ensemble were stratified by topological, structural, and taxonomic contexts.

- **Sub-cellular topology:** Residues were partitioned into cytoplasmic (Cyto), transmembrane helical (TMH), and extracellular (Extra) segments based on PDBTM annotations in concert with TOPDB and Phobius predictions^79,96^.
- **Quaternary interaction partners:** Interacting residues were categorized by the nature of the closest interacting chain based on minimum heavy-atom distance: partners from homomeric/heteromeric complexes of transmembrane proteins or complexes of soluble proteins.
- **Structural complexity and taxonomy:** Proteins were binned by the count of transmembrane *α*-helices (1, 2-3, 4-6, ≥ 7 segments) and taxonomic kingdoms, such aseukaryota, bacteria, archaea, and viruses. Supervised training exposure was mapped via SIFTS annotations^83^ to identify species- and Pfam-level representation^97^.

### Structural profiling of respiratory membrane supercomplexes

To validate macromolecular interface prediction on massive native assemblies, CHORUS was further benchmarked on large-scale respiratory complexes, including four porcine supercomplex cryo-EM structures comprising complexes I, III, and IV^98^ (PDB codes: 8UGH, 8UGJ, 8UGN, 8UGR) and two human complex-IV respirasome assembly intermediates^99^ (PDB codes: 9I6F, 9I7U). Transmembrane subunits were verified by mapping UniProt topological annotations (≥ 80% sequence identity, ≥ 70% helix coverage). Predictions were executed per full-length sequence in single-precision floating-point 32 on CPU without sequence padding or structural features. Interaction sites were determined by an inter-chain heavy-atom distance cutoff ≤ 5Å across both intra-complex and inter-complex interfaces. Performance was quantified across all resolved residues and annotated TM regions.

### Generalization to soluble protein interfaces

Model transferability outside the membrane domain was benchmarked across 7 non-membrane assemblies, including coiled-coil oligomers (cortexillin I, COMP, tetrabrachion) and enzyme-inhibitor complexes (eglin C, stefin B, Im9, ribonuclease inhibitor) (**Supplementary Tables 16-18**). Using the fixed threshold (*t*^∗^ = 0.5542) and a 5Å heavy-atom cutoff for native interfacial residues, CHORUS was compared against baseline predictions derived from Kyte-Doolittle hydropathy ranking^100^ across non-homologous soluble targets. Maximum global sequence identity (< 30%) was assessed against all 8,483 unique canonical chains for training using the BLOSUM62 alignment.

### Proteome-scale sequence selection and inference

To construct a comprehensive resource of transmembrane interaction sites, we extracted all reviewed *α*-helical transmembrane protein entries from UniProt (20 July 2026 release)^54^, which yielded 70,247 unique sequences comprising 33,722,948 residue positions across 80,619 records. Full-sequence inference was performed using the final ensemble model of CHORUS. For polypeptides exceeding 2,174 residues (557 records), predictions were generated using sliding windows of 2,174 residues with a 256-residue overlap. We anchored the terminal window to the end of the sequence of each chain. Probabilities for residue *i* covered by *n*_*i*_ windows were averaged across overlapping predictions as given below

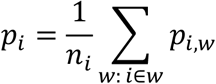

Still, binary interaction states were assigned using the calibrated threshold (*t*^∗^ = 0.5542). Sequence similarity between the Atlas entries and our training data was mapped via MMseqs2 (version: 18-8cc5c, sensitivity 7.5, *E* ≤ 1,000)^101^.

### Atlas database implementation and structural case studies

The Atlas is structured as a relational SQLite resource that links protein accessions, sequence metadata, and predicted interaction probabilities. An interactive interface supports sequence-level querying, threshold filtering, batch retrieval, and real-time mapping onto macromolecular coordinates via Mol* Viewer^102^. The structure lookup first used PDBe accession mappings and then an experimental-structure sequence search through RCSB. A gap-aware sequence alignment mapped the query sequence to the selected structural chain, using match, mismatch and gap scores of 3, -2 and -3, respectively. Only mapped residues received colours from the prediction profile, while unmapped positions were displayed separately. The webserver of the Atlas is made available at https://chorus.atlas.sunnylab.org.

Resource utility was exemplified on the odorant receptor coreceptor Orco^103^ with 478 amino acids from the clonal raider ant Ooceraea biroi (UniProt accession code: A0A026W182). Sequence-derived interaction probabilities were projected onto homologous cryo-EM coordinates (PDB code: 8V00, 77.1% sequence identity, 99.8% coverage), illustrating the spatial distribution of predicted interfacial sites within the intact transmembrane complex. Other examples are shown in **Supplementary Tables 20 and 21**.

### Mapping to biological annotations

Biological association analyses were performed using the family- and depth-controlled ensemble across 70,247 unique reviewed UniProt membrane-protein sequences (30,933,614 residues, deduplicated from 80,619 records to ensure equal statistical weighting). External functional features from BioLiP and CATH were strictly segregated from model training, neural network selection, probability calibration, and inference. Since experimental interface annotations are incomplete across the entire membrane proteome, these analyses only evaluate macro-scale concordance with established structural and functional annotations.

### BioLiP mapping and overlap analysis

To investigate functional coupling between PPI interfaces and protein ligand-binding (PLB) pockets, the CHORUS-predicted interaction sites were benchmarked against experimentally determined receptor contacts in the Q-BioLiP database (version: 10 February 2026)^104^. We downloaded the *qbiolip_PL_annotation.csv* document to obtain annotated protein-ligand binding details. Small-molecule and ion binding sites were mapped to UniProt accessions via SIFTS XML alignments (version: 29.26)^83^, excluding polynucleotide and peptide ligands. Residue correspondences required exact sequence identity and coordinate alignment, defining a high-confidence set of 100,003 verified ligand-binding residues across 2,971 non-redundant sequences (1,171,502 structurally observed residues).

Positions, which were documented in SIFTS but lack annotations ofPLB sites, were retained as background sites. Note that these positions represent an annotation background rather than experimentally established non-PBL sites.

We retained protein chains that contain at least one PLB site and at least one observed background residue, yielding 2,971 chains, 1,171,502 observed residues and all 100,003 PBL sites. For each protein sequence *s*, we denoted the number of observed sites by *N*_*s*_, the number of PBL sites by *M*_*s*_, the number of CHORUS-predicted interaction sites as by *K*_*s*_, and the number of sites overlapped between the PBL sites and the predicted interaction sites by *O*_*s*_ . While preserving the sequence-specific margins, the expected number of overlapping residues was expressed as

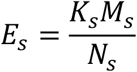

We compared the observed PLB capture fraction as follows

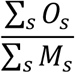

with the corresponding expectation under independence

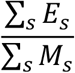

and summarized relative enrichment as

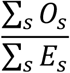

This formulation attempts to evaluate whether CHORUS-predicted interaction sites coincide preferentially with ligand-interacting residues beyond that expected from the sequence-specific abundance of the two site classes. The effect size was expressed as the observed-minus-expected capture percentage, defined as

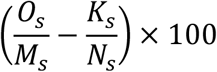

To quantify evolutionary lineage and species-level stratification, we also defined the differential PLB site occupancy within sequences, Δ_occupancy_, phrased as

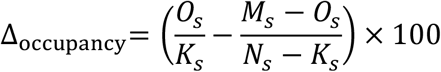

The first term 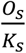 defines the empirical PLB site occupancy within the predicted interfaces, whereas the second term 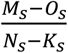 represents the background PLB site occupancy across the remaining non-interfacial residues of the polypeptide chain. The difference (also rephrased as 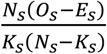 between these two ratios measures whether ligand-binding pockets are enriched (ΔOccupancy_*s*_ > 0), depleted (ΔOccupancy_*s*_ < 0), or mutually independent (ΔOccupancy_*s*_ ≈ 0) relative to the non-interfacial structural surface. Protein chains, which lack either predicted interfacial or non-interfacial observed residues (*n* = 2,652 eligible sequences retained) were excluded from this paired comparison to avoid degenerate denominators.

Two-sided 95% confidence intervals were estimated from 2,000 bootstrap resamples across complete sequences. Sub-cellular topological intervals were retrieved from UniProt (version: 9 August 2026). A total of 2,994 BioLiP-mapped sequences (1,174,365 validated observed residues) were topologically annotated across 80,599 UniProt accessions. Cytoplasmic and cytosolic annotations were designated as cytoplasmic, while extracellular, lumenal, and periplasmic annotations were grouped as extracellular. These annotations were assigned under a strict hierarchy of TOPO_DOM, INTRAMEM, and TRANSMEM from UniProt.

### FunFam mapping and conservation analysis

Functional domains across the 70,247 unique transmembrane protein sequences were assigned by scanning against the CATH (version 4.3.0)^105^ Functional Family (FunFam) hidden Markov model (HMM) library using HMMER (version 3.3.2)^106^ hmmsearch --cut_tc --notextw. Candidate matches were resolved into non-overlapping domain assignments using cath-resolve-hits v0.16.10^107^, requiring a minimum HMM coverage of 80% and a minimum bit score of 25, with a long-domain preference of 2.0. Each query was indexed by the digest of its complete uppercase sequence. Ambiguous residue identifiers were replaced with X for the HMM search only, while the original residue identifiers were retained in the source-coordinate output.

For each validated domain, query residues aligned to HMM match states were mapped through the profile MAP architecture onto standardized FunFam multiple sequence alignment coordinates. Insertions, deletions, and residues flanking the aligned domain boundaries were treated as unobserved conservation states rather than assigned zero-conservation penalties. In the presence of tandem homologous domains within a single polypeptide, positional observations across domain copies were first sequence-wise averaged internally. The global assignment identified 56,221 structural domains across 30,799 unique sequences, spanning 9,564 distinct FunFams and 10,180,315 domain-mapped residues of which 10,018,715 positions aligned directly to consensus HMM match states.

Residue conservation was quantified using the PET91 Scorecons algorithm^108^. Homologous sequences within each FunFam alignment were realigned to their corresponding profile HMMs using hmmalign. Residue conservation was computed across consensus match columns parameterized by the pinned PET91mod.mat2 substitution matrix. In this framework, higher scores denote heightened evolutionary constraints at the corresponding functional consensus position. Continuous Scorecons served as the primary variable, with a predefined threshold of ≥ 0.7 denoting highly conserved functional positions in accordance with established CATH criteria^109^. To guarantee statistical stability and eliminate artifacts arising from shallow alignments, conservation evaluations were restricted to checksum-verified CATH alignments exhibiting a Diversity of Positional Score (DOPS) ≥ 70, a minimum representation of three exact sequences in the current dataset and validated HMM match-state coordinates. No conservation metrics were imputed for uncharacterized families. From the initial cache of 621 official alignments, 580 occurred within the assigned domain cohort. Excluding 69 alignments lacking sufficient sequence support yielded a final high-confidence benchmarking corpus of 9,529 unique sequences, 12,093 resolved domains, and 511 FunFams spanning 114 homologous superfamilies, comprising 2,739,713 Scorecons-annotated residues.

Enrichment of predicted quaternary interfaces within highly conserved positions (Scorecons ≥ 0.7) was tested against the fixed-margin hypergeometric null model, substituting verified ligand-binding sites with highly conserved residue annotations. In parallel, continuous conservation shifts, ΔScorecons, were computed for each sequence by subtracting the mean Scorecons of predicted non-interfacial background residues from that of predicted interfacial residues, denoted as

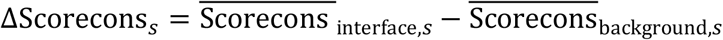

Of the 9,529 mapped sequences, 8,768 contained both predicted classes and were retained for paired evaluation (761 homogeneous sequences were excluded). Multi-compartment stratifications across cytoplasmic (2,878 sequences), extracellular/lumenal (1,802 sequences), transmembrane (7,374 sequences), and intramembrane microenvironments (319 sequences) utilized the identical UniProt topological hierarchy defined in the BioLiP benchmark, with corresponding high-conservation overlap cohorts comprising 3,445, 3,286, 8,349, and 434 sequences, respectively.

At the sub-domain and consensus-state levels, predicted probabilities and binary interaction classifications were averaged across tandem domain repeats and pooled across sequences occupying identical HMM match states. Within each FunFam, monotonic concordance between positional conservation and interface propensity was assessed via Spearman rank correlation across eligible states (supported by ≥ 3 exact sequences, ≥ 5 and ≥ 10 assessed as sensitivity controls), requiring at least ten scoreable states and non-zero variance. FunFam-level estimates were assigned equal weighting, with two-sided 95% confidence intervals derived from 10,000 bootstrap iterations. Domain-level differences in mean and median Scorecons, alongside shifts in the fraction of highly conserved positions, were summarized per FunFam prior to global aggregation. Unbiased probability decile trends were tracked across pooled-residue, equal-FunFam, and family-centered conservation baselines.

Taxonomic comparisons were performed using exact-sequence-level effects after duplicate domain observations had been collapsed within each sequence-FunFam combination. To distinguish conservation shifts occurring within the same FunFam from differences in the FunFam composition represented across taxa, we defined a Δ_*q,f*_ effect for sequence *q* within FunFam *f*, which were standardized against the remaining family members using leave-one-out centering:

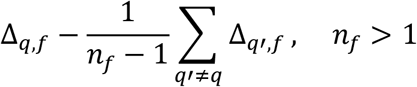

where *n*_*f*_ denotes the sequence count of FunFam *f*. Centered FunFam effects were averaged within each sequence across contributing domains prior to taxonomic grouping (*n* = 8,767 eligible sequences). Taxonomic lineages represented by ≥ 20 sequences were treated as primary comparative groups, and lineages with 10-19 sequences as exploratory. Statistical confidence bounds were estimated via 2,000 sequence-level bootstrap resamples, treating phylogenetic ranks as nested descriptive strata.

### Statistical analysis

We strictly retained the hierarchical distinction between residues, chain contexts and sequence-similarity clusters across all evaluations. The primary benchmark metric was the unweighted mean AP values across eligible contexts. To quantify uncertainty, the resampling unit for bootstrap analyses was the intact sequence cluster, thereby preserving dependencies among its associated contexts and residue-level predictions. For paired method comparisons, identical bootstrap clusters were drawn across models to compute paired metric differences for each replicate. Confidence intervals (95% CIs) correspond to the 2.5th and 97.5th percentiles of the empirical bootstrap distribution.

Specifically, ensemble controls, pairwise constituent comparisons and ablation analyses used 10,000 cluster bootstrap replicates (base random seed: 20260905, with contrast-specific offsets documented in the configuration repository), whereas length-stratified evaluations used 2,000 replicates. Candidate-selection benchmarks were performed over 10,000 replicates (seed: 20260822) with equal weighting assigned to cluster-level AP or Brier scores. For representation-performance regressions, validation clusters (supplying probe scores) and test clusters (supplying interaction-prediction scores) were resampled independently. Representation-level metrics were reconstructed and regressions refitted across 2,000 replicates (seed: 20260717). Reported confidence intervals were pointwise and unadjusted for various comparisons.

Biological subgroup distributions are reported as median cluster AP values with interquartile ranges (IQRs) and sample counts. The 57-candidate benchmark was evaluated using a single fitted model per candidate on a fixed partition of datasets. Variability across the three probe optimization seeds was quantified independently from resampling uncertainty of proteins and does not capture retraining variance of the interaction predictors.

### Software implementation and reproducibility

Data processing and analysis were implemented in Python, using TMKit^110^ and PyPropel^111^ for reusable sequence, structure and feature-processing operations, and Biopython^112^, NumPy^113^, Pandas^114^, and Scikit-learn^115^ supported sequence alignment, numerical analysis, and model evaluation. Downstream prediction backbones and linear probes were implemented in PyTorch^116^. To extract frozen representations, we used fair-esm 2.0.0 (ESM2), Transformers 4.57.6 (ESM-C), and Transformers 5.13.0 with SentencePiece 0.2.1 (ProtT5)^117^. To account for dependency variations across workflows, software environments were recorded per experiment rather than assumed identical across all processing stages.

Model fitting and inference were conducted on NVIDIA A100 80-GB GPUs, whereas full-set evaluation across the development cohort was executed on an A800 GPU. Detailed hyperparameters, including numerical precision, batch size, random seeds, and contextual windowing, were specified in the corresponding sections above. To ensure full reproducibility, the accompanying repository documented the source configurations, model checkpoints, data splits, residue mappings, model outputs, and exact evaluation commands for each *in silico* testing round. Complete parameter settings and the external dataset registry were retained within this reproducibility bundle. Repository links and access details are specified in the Data and Code Availability statements upon release of this work for publication.

## Acknowledgements

We thank the High-Performance Computing (HPC) platform at The Hong Kong University of Science and Technology (Guangzhou) for providing computational resources.

## Declarations

The authors declare no competing interests.

## Data availability

The CHORUS-predicted atlas of protein-protein interacting residues in membrane proteomes across kingdoms is accessible at https://chorus.atlas.sunnylab.org.

## Code availability

The CHORUS method is made publicly available as a standalone GitHub package at https://github.com/sunnylab-cb/chorus and as a webserver at https://chorus.server.sunnylab.org.

## Author contributions

J.S. conceived this project. J.Z. and S.H. collected the data, performed the analysis, developed the algorithm, and prepared the draft. J.S. provided supervision, investigated the literature, collected the data, interpreted the results, and revised the manuscript. All authors have given approval to the final version of the manuscript.

