## Extended Figures for "CHORUS: A proteome-scale atlas of membrane protein interaction interfaces predicted from sequence"

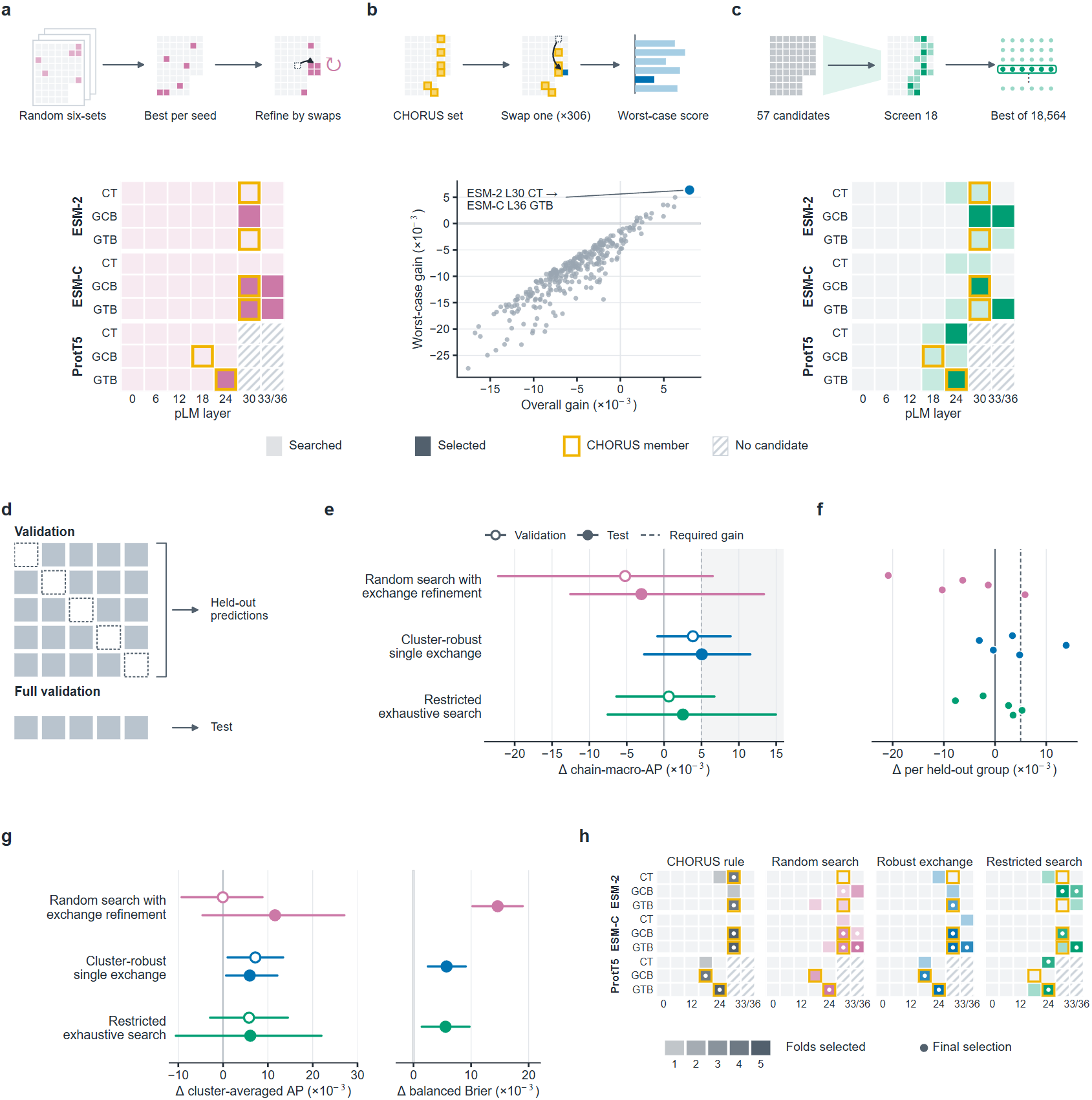

**Extended Data Fig. 1 | Illustration of in silico approaches for determining high-performance ensemble models using average-precision (AP). a-c**, Three schemes posited to reconstruct ensemble models. Each chooses six of the 57 single-layer candidates. The panels to the upper show schematics of search for ensemble models. The grids represent the candidate pool. The panels to the lower show the selection fitted on the full validation dataset. Dark cells in the matrix showed the models of choice. Gold outlines mark the individual models in CHORUS. Hatched cells have no candidate due to a prespecified setting for ProtT5 layers (Methods). **a**, Random-search-based replacement of a single model per round. For each of 3 seeds, the scheme retained the best among 299 ensemble sets (our predefined number) composed of 6 arbitrarily selected individual models according to the chain-macro AP metric. It then swapped single model while chain-macro AP values increased. **b**, Cluster-based replacement of a single model per round. Starting from the set of models in CHORUS, this scheme scored each of the 306 swaps (i.e., $6\times(57-6)$) using the worst-case gain across three cluster weightings and the cluster and cluster-group deletions (Methods). It kept the swap with the largest worst-case gain. Each scatter point indicates one swap with its gain in chain-macro AP over all validation protein clusters (x-axis) against its worst-case gain (y-axis). The selected swap colored in blue and 14 others had a positive worst-case gain. **c**, Restricted exhaustive search for ensemble models. Within each pLM family, the scheme screened the two candidates with the highest chain-macro AP values and the next four with the highest cluster-macro AP values, resulting in a total of 18 candidates. It then evaluated all 18,564 possible combinations of models and kept the one with the highest cluster-macro AP value. **d**, Evaluation settings. Within validation, the 65 protein clusters were divided into five groups. An ensemble model acquired by applying each of the three schemes was fitted on four validation groups, leaving the rest for validation. The ensemble model was also fitted on the full validation dataset and further evaluated on our test dataset composed of 55 protein clusters. **e**, Chain-macro AP performance of the three schemes, with paired cluster bootstrap 95% intervals (10,000 replicates). Open circles indicate the results on the validation dataset, while filled circles indicate the results on the validation dataset. Crucially, a scheme led to an improvement only if its gain reached the dashed line (+0.005) and its interval excluded zero. **f**, Chain-macro AP performance on each of the 5 held-out validation datasets. **g**, Cluster-macro AP performance (the left panel) on the validation and test datasets and class-balanced Brier scores (the right panel) on the test dataset. **h**, Comparison of ultimately chosen ensemble models across different schemes. Dots mark the models of choice.

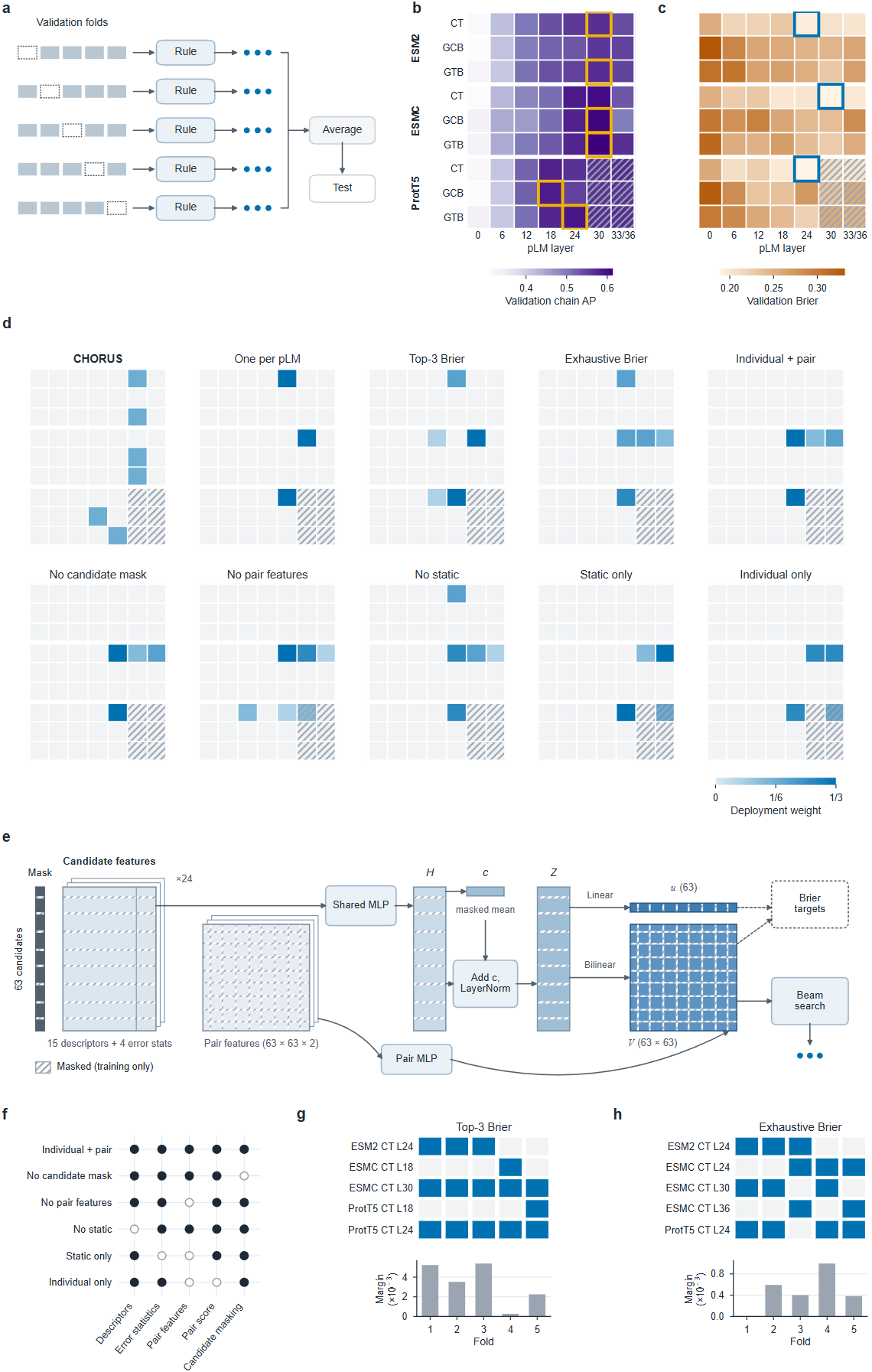

**Extended Data Fig. 2 | Ensemble-selection rules. a**, Sketch of model selection schemes trained on four folds (filled squares) and tested on different validation partitions (dashed squares) composed of protein chains with each pair sharing less than 30% sequence identity. **b,c,** Chain-macro AP values (**b**) and cluster-averaged Brier scores (**c**) on the validation dataset across the total of 63 models using pLM layers of various pLM methods (layer 33 for ESM2 and layer 36 for ESMC) and prediction heads. Hatched cells for ProtT5 are concatenated layers 0-24 and 6-24, which were made available only to the replacement rules. Gold outlines mark the models of choice used ultimately to assemble CHORUS, while blue outlines mark the model per pLM family with the lowest Brier scores (corresponding to the highest performance). **d**, Choices of models determined by various scoring schemes. The $\frac{1}{3}$ development weight is taken as the upper-bound threshold for model selection across all 5 folds (See **Supplementary Note 1**). **e**, A neural network-based scorer trained to reproduce the contribution of each candidate model and paired models. A shared two-layer perceptron encodes descriptors of each candidate and validation error statistics. The encodings are combined with their mean over the candidate set and layer normalized. A linear head is introduced to yield a list of Brier scores $u$ for individual models, while a bilinear head together with a perceptron on residual-error correlations is used to yield a matrix of Brier scores $v_{ij}$ for pairs of models. Beam search selects the combination of models that maximizes $\sum u_{i}+\sum v_{ij}$. During training, candidate models are randomly masked (crossed squares, 15% per epoch), which removes them from the set mean and the loss. **f**, Inputs and score terms retained (filled) or removed (open) in each neural variant. **g,h,** Fold-specific selections (top) and winning margins (bottom) of Top-3 Brier (Brier gap between the fourth- and third-ranked candidates, **g**) and Exhaustive Brier (Brier gap between the second-best and best of all 39,711 teams, **h**). Each fold uses different fitting data and the margins are small, so near-tied candidates exchange places between folds. Exhaustive Brier scores whole teams and can therefore include a predictor with a higher individual Brier (ESMC CT L24) that lowers the error of a model combination.

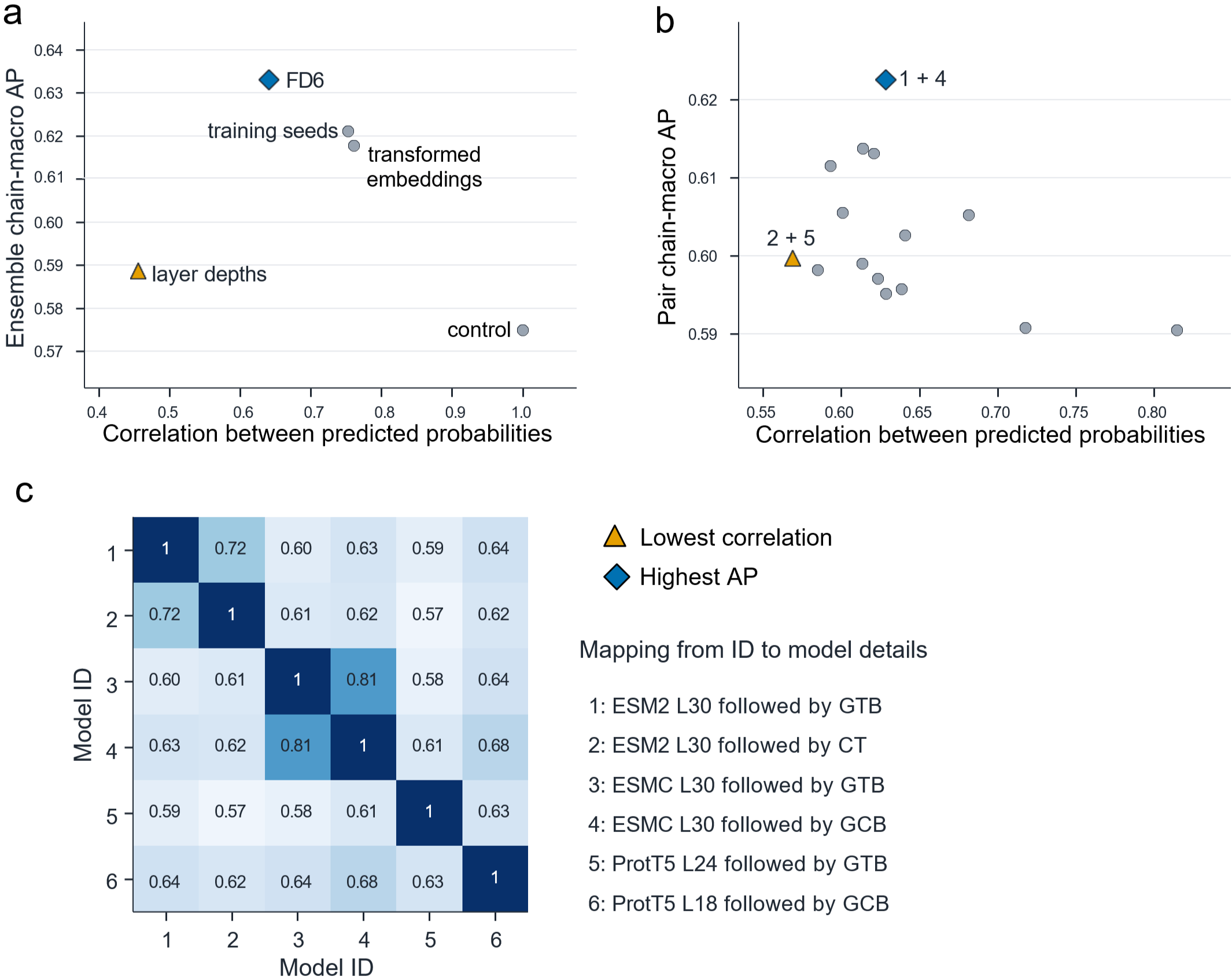

**Extended Data Fig. 3 | Prediction correlation and ensemble performance.** **a,** Relationship between the mean pairwise correlation of predicted probabilities of interacting residues and the chain-macro AP values with different ensemble models constructed using different training seeds, signed-permuted embeddings, ESM2 representation layers or identical predictions (control). Correlation was calculated as the mean Pearson correlation coefficient across all 15 pairs of predicted probabilities within each ensemble. The FD6 ensemble was built by merging six individual models headed only to GTB yet spanning three pLM families and two representation layers per family, distinct from the final CHORUS ensemble. The lowest-correlation ensemble (ESM2 depths) and highest-AP ensemble (FD6) are highlighted. **b,** Pairwise Pearson correlation between predictions versus chain-macro AP for all 15 pairwise combinations of the six individual models comprising CHORUS. Each pair was evaluated by equally averaging its two predicted probabilities. The pair 2 + 5 combination exhibited the lowest prediction correlation, whereas the pair 1 + 4 combination achieved the highest chain-macro AP value. **c,** Pairwise Pearson correlation matrix of residue-level predictions from the six CHORUS individual models. Correlations were calculated across pooled test residues, whereas the chain-macro AP values were computed by equally averaging across protein chains.

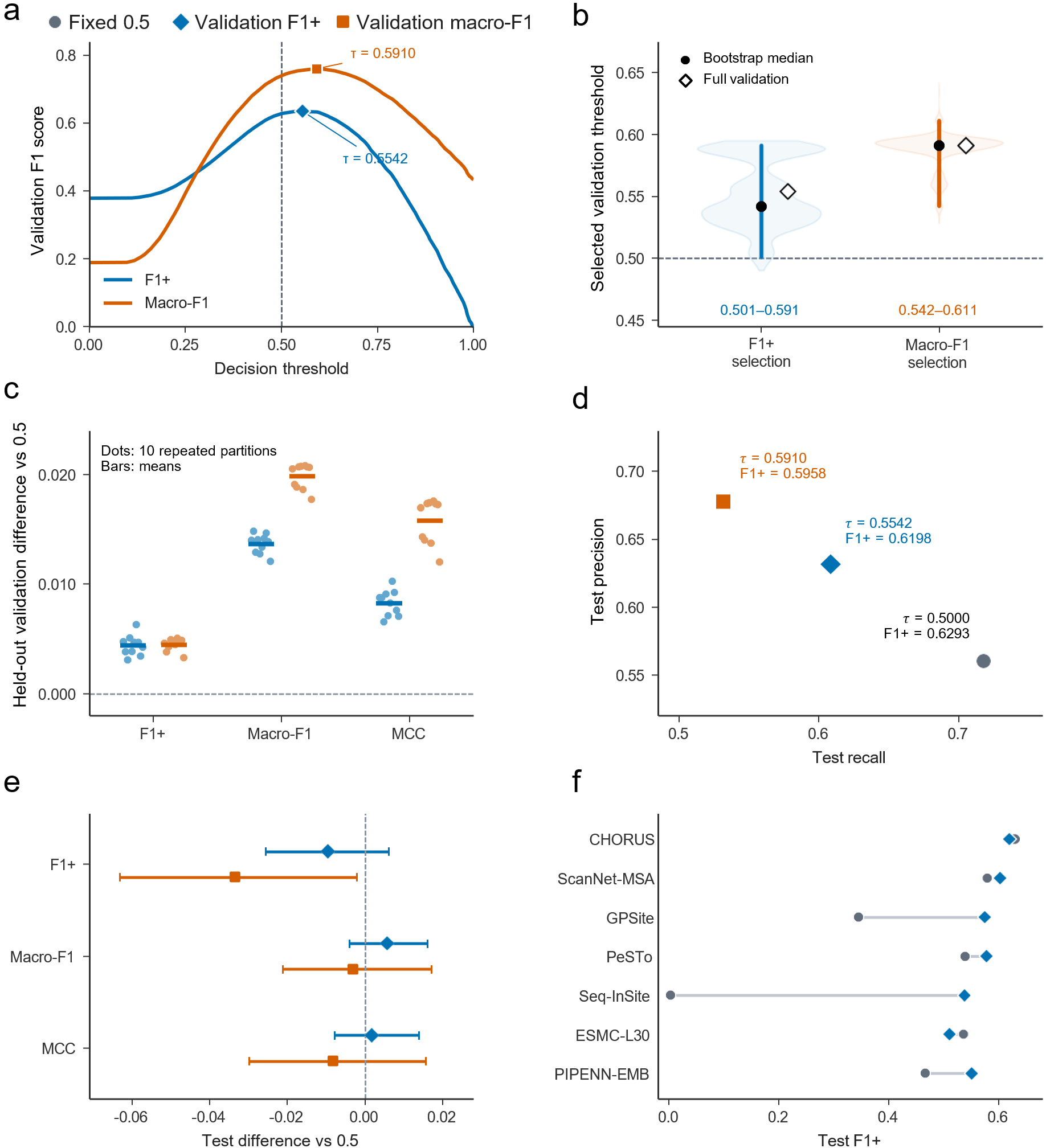

**Extended Data Fig. 4 | Selection of decision thresholds and sensitivity of predictive performance. a,** F1-scores of predicted interacting residues on the validation dataset (Validation F1+) and macro-F1 (Validation macro-F1) across various thresholds for CHORUS, with optimal thresholds of 0.5542 and 0.5910, respectively. **b,** Distribution of optimal thresholds from 2,000 bootstrap resamples of 65 clusters from the validation dataset. Black circles indicate medians. Diamonds indicate full-validation optima. Vertical bars denote central 95% ranges. **c,** Held-out validation performance difference relative to the fixed threshold of 0.5 across ten repeated five folds over protein clusters. Dots represent repetitions and horizontal bars indicate means. **d,** Test precision-recall points at the fixed and validation-selected thresholds, annotated with F1+. **e,** Paired test difference in F1+, macro-F1, and Matthews correlation coefficient (MCC) relative to the fixed threshold of 0.5. Error bars indicate percentile 95% confidence intervals from 2,000 cluster bootstrap resamples. **f,** Test F1+ under fixed and validation-selected thresholds for seven benchmark methods. CHORUS predictions were used in these analyses.

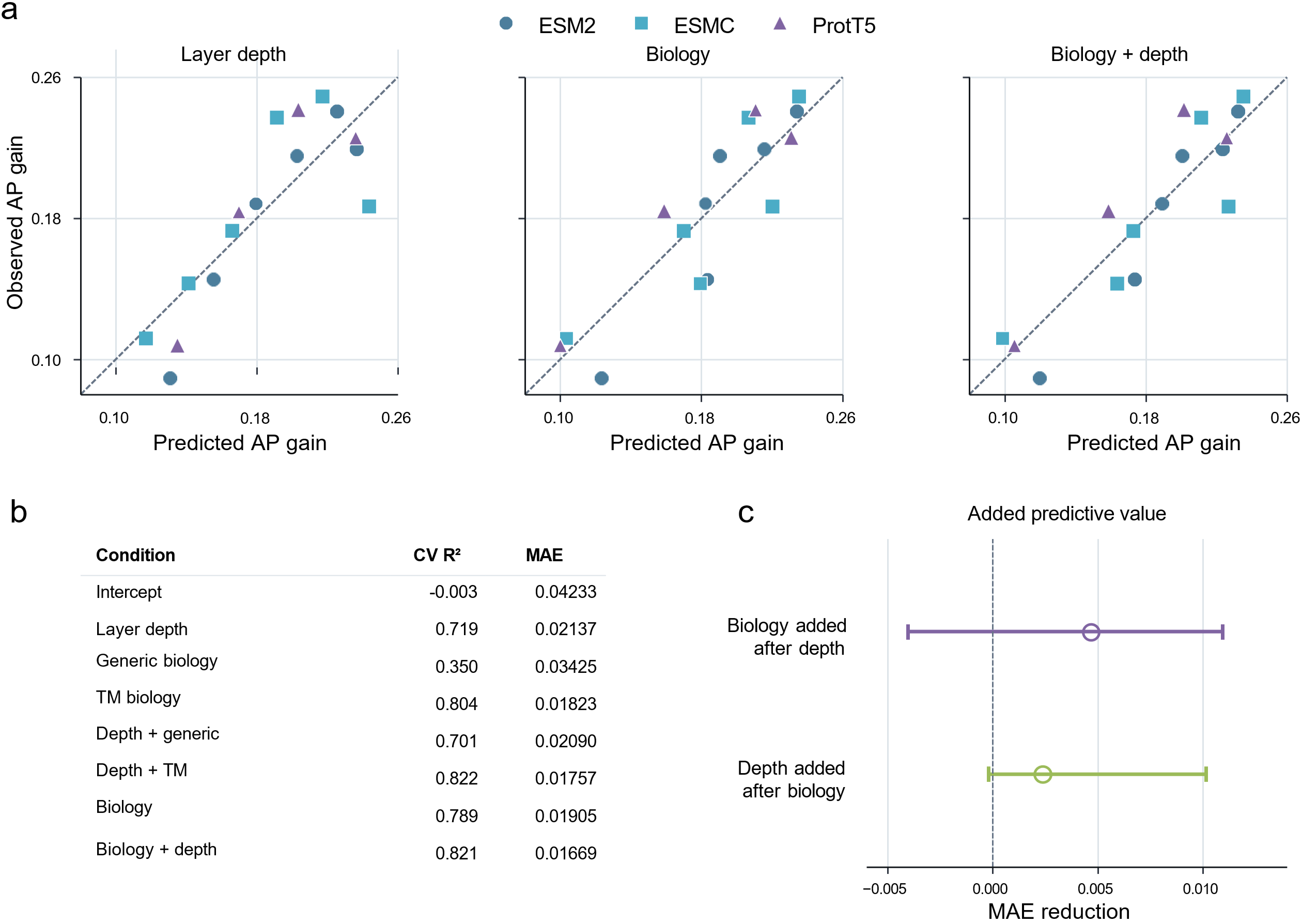

**Extended Data Fig. 5 | Representation depth and biological profiles predict layer-specific performance. a,** Comparison between observed and predicted AP gains across 16 pLM layers spanning from ESM2, ESMC, to ProtT5, evaluated using leave-one-pLM-out cross-validation. AP gains were calculated relative to the layer-0 baseline of each pLM and averaged across three prediction heads. **b,** Cross-validated $R^{2}$ and mean absolute error (MAE) for models incorporating representation depth, generic biology (local contacts and conservation), TM-specific biology (TM location, helix position, membrane depth, and helix neighbours), and their combinations (see **Methods**). **c,** Incremental MAE reduction from adding biological profiles to depth or depth to biological profiles. Points represent observed improvements, with 95% intervals from 2,000 cluster bootstrap resamples.

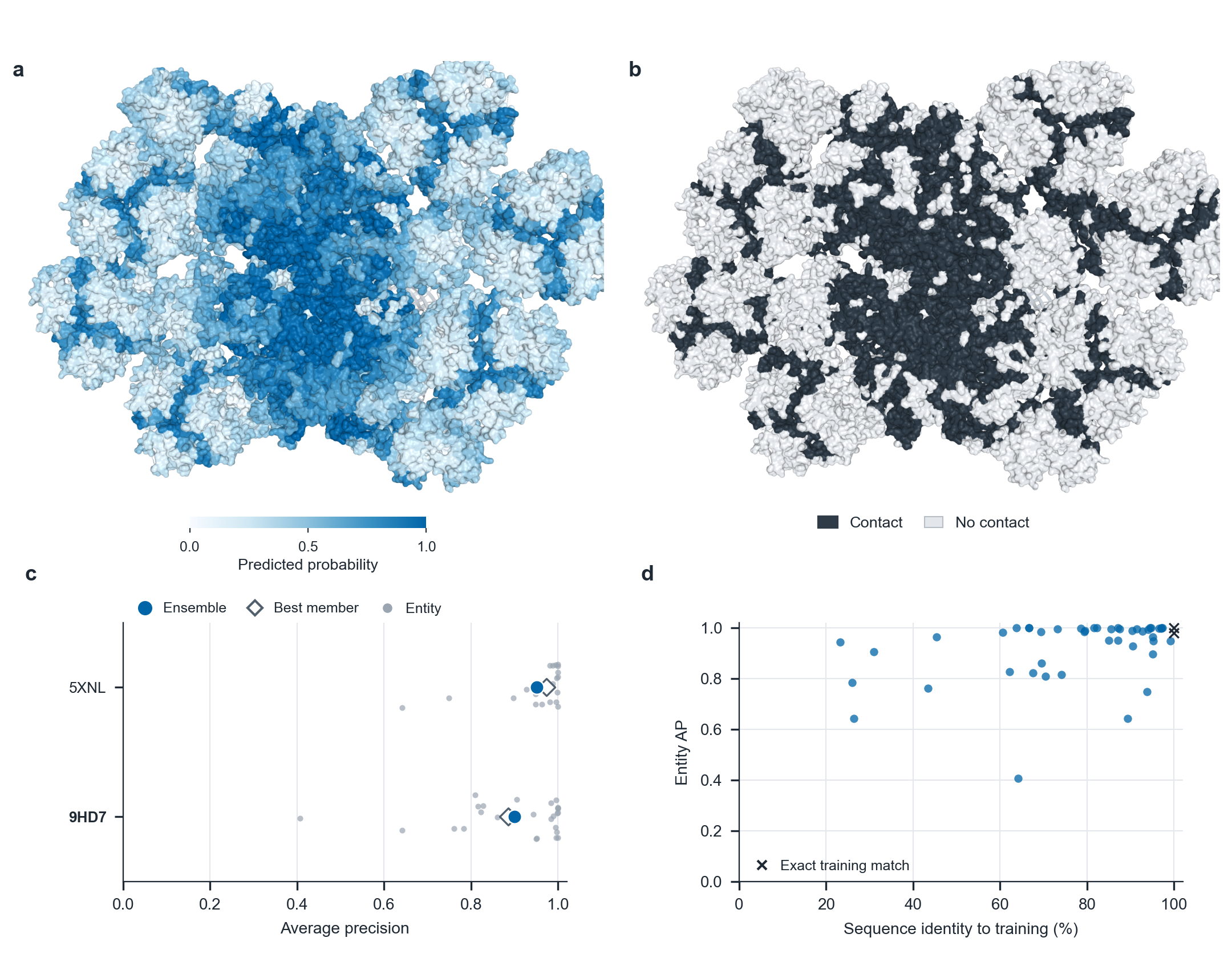

**Extended Data Fig. 6 | Analysis of interfaces on PSII-LHCII supercomplexes.** **a,b,** Structural representation of probabilities of residues predicted as interacting (**a**) and actual interaction interfaces colored in black (**b**) on chlorella ohadii PSII-LHCII supercomplex (PDB code: 9HD7). The interaction partners are visualized as grey ribbons. **c,** Performance between CHORUS and its constituent models/members for chlorella ohadii PSII-LHCII and pea PSII-LHCII (PDB code: 5XNL). **d,** Entity AP against sequence identity to training proteins. The cross mark represents the exact match with training data.

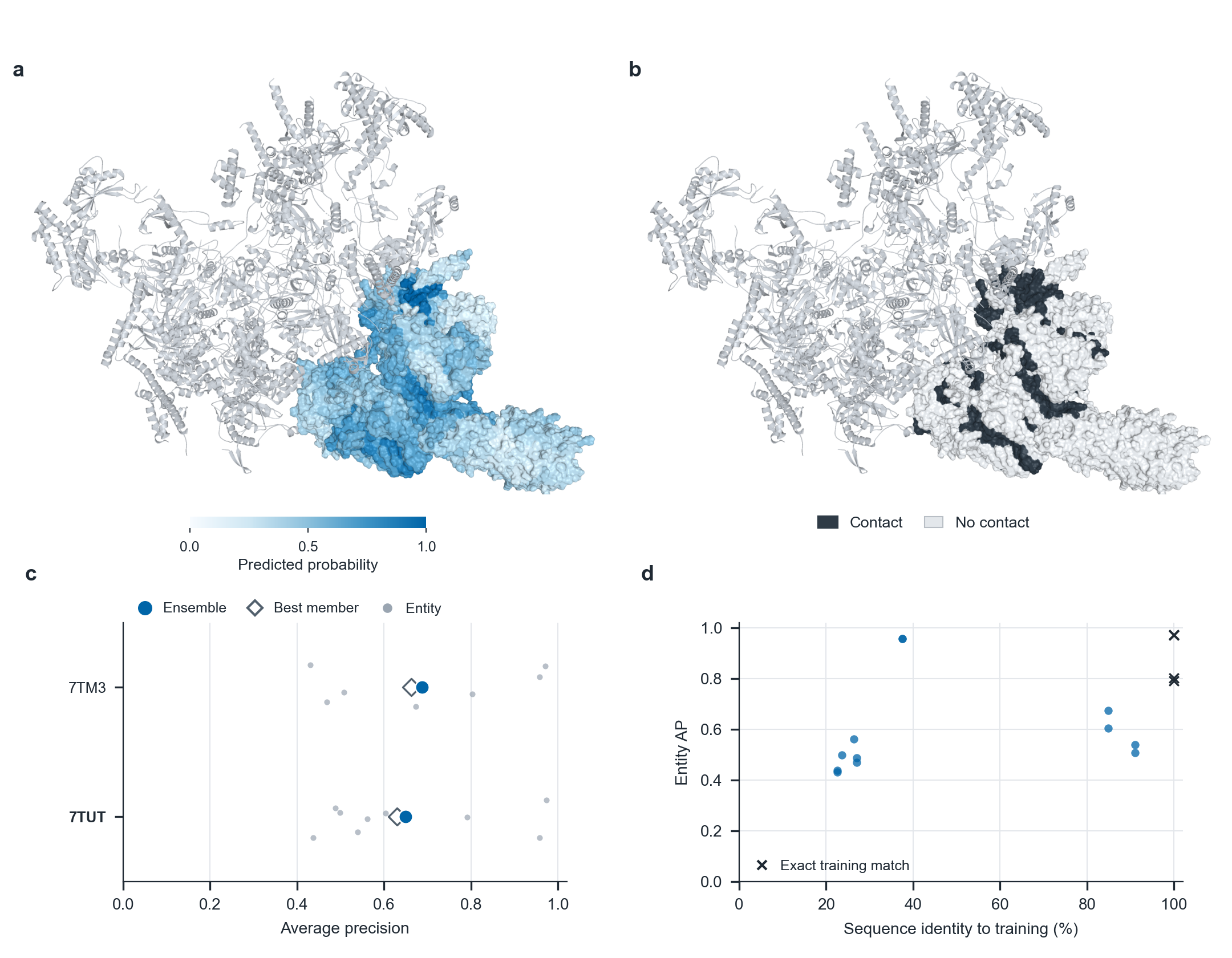

**Extended Data Fig. 7 | Analysis of interfaces on ER multipass translocon. a,b,** Structural representation of probabilities of residues predicted as interacting (**a**) and actual interaction interfaces colored in black (**b**) on ribosome-bound multipass translocon with a nascent rhodopsin intermediate (PDB code: 7TUT). The interaction partners are visualized as grey ribbons. **c,** Performance between CHORUS and its constituent models/members for two insertion intermediates. **d,** Entity AP against sequence identity to training proteins. The cross mark represents the exact match with training data.
