## Supplementary Information for "CHORUS: A proteome-scale atlas of membrane protein interaction interfaces predicted from sequence"

**Supplementary Note 1**

To understand the organization of sequence-level representations across pLM families, we compared the geometry of ESM-2, ESM-C and ProtT5 embeddings against a common protein-interaction reference (**Supplementary Figs. 2 and 3**). We observe that cross-model agreement among the 15 nearest reference-protein identities was limited, with mean Jaccard overlaps ranging from 0.060 to 0.107, indicating that the selected pLM representations retrieved substantially different local neighborhoods. Membrane-protein queries also showed preferential proximity to independently annotated membrane-protein anchors under the evaluated reference definitions. However, reciprocal reference-support analyses did not identify a consistent direction across all settings, and two-dimensional projections did not fully preserve the original high-dimensional neighborhoods. These analyses therefore characterize differences in representation-space organization rather than establish a direct mechanistic relationship between embedding geometry and residue-level interface prediction performance.

**Supplementary Note 2**

***Probability averaging***. For an input protein sequence $S$ of length $L$, let

$$\mathbf{H}^{\left( f\mathcal{,l} \right)}\left( S \right)\in\mathbb{R}^{L\times d_{f}}$$

denote the frozen representations with a dimension $d$ extracted from layer $\mathcal{l}$ of pLM $f$, where $f\in\left\{ ESM2,\mathrm{ESMC},ProtT5 \right\}$.

Let $g_{\theta}^{\left( a \right)}$ denote a trained model with a head/backbone $a\in\left\{ \mathrm{GCB},\mathrm{CT},\mathrm{GTB} \right\}$ and trainable parameters $\theta$. Its prediction for residue $r$ can be expressed as

$$p_{r}^{\left( f\mathcal{,l,}a \right)}=g_{\theta}^{\left( a \right)}\left( \mathbf{H}^{\left( f\mathcal{,l} \right)}\left( S \right) \right)_{r}$$

For feature-enabled GCB predictors, the network additionally incorporates the corresponding sequence-derived auxiliary features. All ensembles consisting of six individual models aggregate their predicted probabilities by the same unweighted mean

$$\hat{p}_{r}^{\mathrm{ensemble}}=\frac{1}{6}\sum_{j=1}^{6} p_{r}^{\left( j \right)}$$

CHORUS retains the top two individual models within each pLM family, ranked by the chain-macro AP performance on the validation dataset. The selected configurations are ESM2-L30/GTB, ESM2-L30/CT, ESMC-L30/GTB, ESMC-L30/GCB, ProtT5-L24/GTB, and ProtT5-L18/GCB. For the ensemble CHORUS model (denoted as $M_{\mathrm{CHORUS}}$), its ensemble predicted probability of residue $r$ is written as

$$\hat{p}_{r}^{\mathrm{CHORUS}}=\frac{1}{6}\sum_{m\in M_{CHORUS}} p_{r}^{(m)}$$

The selected combination spans pLM families, representation layers and prediction architectures. This comparator tests the released CHORUS construction, without isolating an independent causal effect of any one diversity source.

***Chain-count weighting***. The weighting scheme sets $w_{k,s}$ to $n_{k}$, which assigns the same effective weight to each AP-eligible chain context. Consequently, clusters containing more chain contexts contribute more heavily to the aggregate score. When no clusters are removed, this definition recovers the chain-macro AP

$$AP_{\mathrm{chain}}(S)=\frac{\sum_{k=1}^{K} n_{k} \overline{AP}_{k}\left( S \right)}{\sum_{k=1}^{K} n_{k}}$$

***Equal-cluster weighting***. The weighting scheme sets $w_{k,s}$ to $1$, which assigns equal aggregate weight to every retained cluster, irrespective of its size. It reduces the influence of clusters represented by many closely related chain contexts

$$AP_{\mathrm{cluster}}(S)=\frac{1}{K}\sum_{k=1}^{K} \overline{AP}_{k}\left( S \right)$$

for the complete validation partition.

***Square-root cluster-size weighting***. The weighting scheme sets $w_{k,s}$ to $\sqrt{n_{k}}$, which provides an intermediate weighting between chain-count and equal-cluster schemes. Larger clusters retain greater influence, but their weights grow sublinearly with the number of chain contexts

$$AP_{\mathrm{sqrt}}(S)=\frac{\sum_{k=1}^{K} \sqrt{n_{k}} \overline{AP}_{k}\left( S \right)}{\sum_{k=1}^{K} \sqrt{n_{k}}}$$

***Overall gain***. As shown in **Extended Data Fig. 1b**, the overall gain measures the difference in AP between CHORUS and a modified ensemble (i.e., a combination of six individual models), evaluated across all validation clusters based on the chain-macro AP metric, defined as

$$\Delta AP_{overall}=AP_{new}-AP_{CHORUS}$$

A positive overall gain indicates an improvement in chain-macro AP values relative to the original CHORUS ensemble on the validation data following replacement of an individual model.

***Worst-case gain***. For each round of model replacement, the improvement in AP was assessed under the three cluster-weighting schemes and the cluster-level and cluster-group deletion scenarios specified in the Methods. The worst-case gain is the smallest AP improvement observed across these evaluation conditions

$$\Delta{AP}_{worst}=\min_{w\in\mathcal{W}} [AP\left( S^{'};w \right)-AP(S_{CHORUS};w)]$$

where $S^{'}$ denotes an ensemble obtained through individual model replacement, and 𝒲 denotes the collection of prespecified cluster-weighting and deletion conditions. A positive worst-case gain indicates that the exchange improves AP under every evaluated condition.

**Maximin selection criterion**

This strategy can be formulated as a maximin robust optimization problem, given by

$$S^{*}=\underset{S^{'}\in\mathcal{N(}S_{CHORUS})}{arg max} \min_{w\in\mathcal{W}} \Delta AP\left( S^{'};w \right)$$

In this expression, $\mathcal{N(}S_{CHORUS})$ is the set of 306 possible individual-model exchanges, $\mathcal{W}$ contains the evaluated cluster-weighting and deletion conditions, and $\Delta AP\left( S^{'};w \right)$ is the difference in AP between the modified ensemble and CHORUS under condition $w$. The objective is not simply to maximize AP on the full validation partition, but to select the exchange with the largest improvement under the least favourable condition.

***Pair average precision (AP)***. For each pair of models, $a,b$, probabilities of residue $r$ being an interaction site in protein $i$, were first averaged with equal weights.

$$\hat{p}_{i,r}^{(a,b)}=\frac{p_{i,r}^{\left( a \right)}+p_{i,r}^{\left( b \right)}}{2}$$

Residues were then ranked in descending order of their averaged probabilities. AP was calculated against ground-truth labels.

$${AP}^{(a,b)}=\sum_{k=1}^{N} \left( R_{k}-R_{k-1} \right)P_{k}$$

Here, $N$ denotes the number of residues in a protein sequence, while $P_{k}$ and $R_{k}$ represent the precision and recall, respectively, at the $k$-th position in the ranked list.

The primary aggregate metric is the arithmetic mean across eligible chains $\mathcal{E}$.

$${AP}_{chain}^{(a,b)}=\frac{1}{|\mathcal{E}|}\sum_{i\mathcal{\in E}} {AP}_{i}$$

where $f$ denotes either an individual model or a two-model probability average. Each eligible chain contributes equally, regardless of length. Chains without reference contact-positive residues were omitted under the study’s AP-eligibility rule.

Each individual model outputs a probability for a residue $r$. Let $p_{i,r}^{(a)}\in\left[ 0,1 \right]$ be the predicted probability of residue $r$ being an interaction site of protein $i$ from individual model $j$. Let $y_{i,r}\in\{0,1\}$ be its ground-truth label. The pairwise gain is then calculated on the same independent-test protein chains.

$$t^{*}=arg\max_{t\in\{a,b\}} {AP}_{chain}^{(t)}$$

$$\Delta{AP}^{(a,b)}={AP}_{chain}^{(a,b)}-{AP}_{chain}^{(t^{*})}$$

The Pearson correlation was introduced to measure the similarity between probabilities of any pair of models in per-residue comparison manner, written by

$$\rho^{(a,b)}=\frac{\sum_{r\mathcal{\in R}} (p_{r}^{\left( a \right)}-\bar{p}^{(a)})(p_{r}^{\left( b \right)}-\bar{p}^{(b)})}{\sqrt{\sum_{r\mathcal{\in R}} {(p_{r}^{\left( a \right)}-\bar{p}^{(a)})}^{2}\sum_{r\mathcal{\in R}} {(p_{r}^{\left( b \right)}-\bar{p}^{(b)})}^{2}}}$$

We investigated whether averaging any pair of individual models out of six models in CHORUS improves the chain-macro AP performance. With six individual models, the set of unordered model pairs, $a,b$, is shown below

$$\mathcal{P=\{}\left( a,b \right):1\leq a<b\leq6\}, \left| \mathcal{P} \right|=\binom{6}{2}=15$$

Collecting repeated occurrences of the same candidate across folds yields a single weighted ensemble, with each candidate receiving weight proportional to its selection frequency.

$$w_{i}=\frac{1}{15}\sum_{f=1}^{5} \mathbf{1}\{i\in S_{f}\}$$

$$\hat{p}_{r}=\sum_{i\mathcal{\in U}} w_{i}p_{ir}$$

Under this weighting convention, all deployment weights sum to one, and no predictor can exceed one-third of the total weight

$$\sum_{i\mathcal{\in U}} w_{i}=1, 0<w_{i}\leq\frac{1}{3}$$

***Initialization diversity***. The same frozen ESM2 layer-30 representations were taken input into the GTB head, which were then trained with 6 different random seeds s = 0, 1, 2, 3, 4 and 5. The resulting predicted probability

$$\hat{p}_{r}^{\mathrm{seed}}=\frac{1}{6}\sum_{s=0}^{5} g_{\theta_{s}}^{\mathrm{GTB}}\left( H^{\left( ESM2,30 \right)} \right)_{r}$$

Thus, the pLM input and backbone design are fixed, while stochastic training and the resulting fitted model parameters vary.

**Orthogonal** **transformation of pLM embeddings.** We used the original ESM2 layer-30 embeddings and five fixed signed permutations. For variant $j$, define an orthogonal transformation $Q_{j}$ as the product of a permutation matrix $P_{j}$ and a diagonal sign matrix $D_{j}$:

$$\mathbf{Q}_{j}=\mathbf{P}_{j}\mathbf{D}_{j}, \mathbf{Q}_{j}^{T}\mathbf{Q}_{j}=\mathbf{I}$$

With $\mathbf{Q}_{0}=I$ for the original feature coordinates, the corresponding transformed embedding matrix is

$${\tilde{\mathbf{H}}}_{j}=\mathbf{H}^{\left( ESM2,30 \right)}\mathbf{Q}_{j}^{T}$$

The predictions of six separately fitted GTB networks are then averaged

$$\hat{p}_{r}^{\mathrm{te}}=\frac{1}{6}\sum_{j=0}^{5} g_{\phi_{j}}^{\mathrm{GTB}}\left( {\tilde{\mathbf{H}}}_{j} \right)_{r}$$

Because each signed permutation is orthogonal, both Euclidean distances and inner products are preserved.

$$\left\| \mathbf{Q}_{j}\mathbf{u-}\mathbf{Q}_{j}\mathbf{v} \right\|_{2}=\left\| \mathbf{u}-\mathbf{v} \right\|_{2}$$

$$\left( \mathbf{Q}_{j}\mathbf{u} \right)^{T}\left( \mathbf{Q}_{j}\mathbf{u} \right)=\mathbf{u}^{T}\mathbf{v}$$

Accordingly, this control changes the coordinate basis presented to the trainable network without supplying additional protein-sequence information. Differences in fitted predictions may arise because the network and its training procedure are not necessarily invariant to this reparameterization.

***Six ESM2 layers for representation depth diversity***. Six GTB predictors use frozen ESM2 representations extracted from layers 0, 6, 12, 18, 24, and 30, each connected to the GTB head. Their averaged probability is

$$\hat{p}_{r}^{\mathrm{depth}}=\frac{1}{6}\sum_{\mathcal{l\in L}} g_{\theta_{\mathcal{l}}}^{\mathrm{GTB}}{(\mathbf{H}^{\left( ESM2,\mathcal{l} \right)})}_{r}$$

This control tests whether mixing representation depths within ESM2 alone can reproduce the CHORUS ensemble performance, without mixing distinct pLM families or backbone architectures.

***Six identical predictions for zero-diversity duplication.*** Let $p_{r}^{(0)}$ denote the residue-level output of one fixed ESM2-L30/GTB predictor. Averaging six exact copies gives:

$$\hat{p}_{r}^{\mathrm{identical}}=\frac{1}{6}\sum_{j=1}^{6} p_{r}^{(0)}=p_{r}^{(0)}$$

The equality is exact: duplicating a fixed probability stream does not alter the prediction and therefore cannot generate an ensemble-averaging gain.

***Enlarged head structure with enlarged parameters downstream capacity scaling.*** The final control uses a single higher-capacity ESM2-L30/GTB model rather than an ensemble:

$$\hat{p}_{r}^{\mathrm{large}}=g_{\Theta_{\mathrm{large}}}^{\mathrm{GTB}}\left( \mathbf{H}^{\left( ESM2,30 \right)} \right)_{r}$$

Its architecture employs a projection width of 640, 960 BiGRU units per direction, two Transformer layers and eight attention heads. The downstream trainable parameter counts are

$$N_{\mathrm{large}}=27,691,521$$

$$N_{\mathrm{CHORUS}}\approx24.38\times{10}^{6}$$

Frozen pLM parameters are excluded from both parameter counts. This comparator tests whether expanding a single prediction network can reproduce the CHORUS ensemble performance.

**Supplementary Note 3**

**Predictive complementarity underlies the ensemble performance of CHORUS**

We examined the predictive complementarity and individual contributions of the six models comprising CHORUS (**Supplementary Tables 2** and **3**). Uncertainty in pairwise AP gains was estimated using 10,000 paired cluster-level bootstrap resamples on the independent test set. Of the 55 test clusters, 54 were retained for resampling but the remaining cluster was excluded from performance evaluation in that it contained proteins with no interacting residues. For each bootstrap replicate, 54 clusters were sampled uniformly with replacement, and the AP values for the pairwise individual model were computed on the complete sets of proteins associated with the sampled clusters. All 15 pairwise combinations improved the chain-macro AP performance relative to their better-performing constituent model, with gains ranging from 0.0155 to 0.0609 and positive cluster-resampled 95% confidence intervals (**Supplementary Table 6**). Notably, the least-correlated pair (models 2 and 5; $r=0.5690$) did not achieve the highest pairwise AP, whereas models 1 and 4 attained the best pairwise performance (AP: 0.6226) despite exhibiting a higher correlation ($r=0.6286$) (**Extended Data Fig. 3b,c**). This further demonstrates that prediction correlation alone does not determine the benefit of combining models. Leave-one-out ablations produced reductions in AP ranging from 0.0018 to 0.0082, although the confidence intervals for several individual-model removals included zero. Removing both models associated with a given pLM family resulted in larger point-estimate reductions of 0.0149-0.0161, with positive confidence intervals for the ESM2 and ESMC family removals (**Supplementary Table 5**). All 15 model pairs (i.e., $\binom{6}{2}$) exhibited positive AP gains, with individual 95% bootstrap confidence intervals entirely above zero. Together, these analyses support the value of combining complementary representations and prediction architectures, while suggesting that the ensemble advantage arises from their collective contributions rather than dependence on any single individual model.

**Supplementary Note 4**

**Evaluation of alternative model-selection strategies**

To gain an understanding of whether the current combination of individual models in CHORUS (**Supplementary Table 3**) suffices for an optimal choice or whether alternative approaches perform better, we developed several strategies to identify the optimal model ensemble from a large pool of individual models, including one-per-pLM, top-3-Brier, exhaustive-Brier, individual-pair, no candidate mask, no pair features, no static, static only, and individual only (**Supplementary Table 7**). The one-per-pLM strategy is built to select the candidate model with the lowest cluster-averaged Brier score within each pLM family (ESM2, ESMC and ProtT5) and combine the three selected models by unweighted averaging of their predicted probabilities. The top-3-Brier strategy is constructed to select the three candidate models with the lowest individual-model cluster-averaged Brier scores, irrespective of pLM family, and combine their predicted probabilities by unweighted averaging. The exhaustive-Brier strategy is set to evaluate all eligible three-model combinations using the cluster-averaged Brier score of their equally averaged predicted probabilities and select the combination with the lowest score. The individual-pair strategy is introduced to train a perceptron-based scorer to estimate individual-model and pairwise contributions to ensemble performance and use these estimates to guide beam search for a three-model combination. Note that the top-3-Brier strategy selects models according to their individual performance, whereas the exhaustive-Brier strategy selects a combination according to its joint predictive performance. The no-candidate-mask strategy disables random candidate masking during training while retaining all input features and scoring terms. This allows assessment of the contribution of candidate masking to selection performance. The no-pair-features strategy removes explicit pairwise error features while retaining pairwise scoring. The no-static-variant strategy removes model-configuration descriptors while retaining individual-model error statistics and pairwise error features. The static-only strategy uses model-configuration descriptors as its sole inputs while retaining both individual and pairwise scoring terms. The individual-only strategy removes explicit pairwise error features and pairwise scoring terms and selects combinations using learned individual-model scores to assess the contribution of explicit pairwise modeling.

Moreover, three strategies, including exhaustive search, beam search, and direct selection, were taken to perform the ensemble model search (**Supplementary Table 8**). The exhaustive search evaluates all 39,711 candidate combinations (i.e., $\binom{63}{3}$) to identify the optimum under a specified scoring function, whereas the beam search explores a restricted subset to reduce the computational cost without guaranteeing global optimality. Both search strategies rely on a two-layer perceptron neural network for model scoring. For simplicity, we refer to this trained network as a neural selector, which was used to disentangle the positive correlations among these 63 individual models (**Extended Data Fig. 2e**). The direct selection applies predefined rules without neural-selector-guided combinatorial search.

We systematically evaluated alternative model-selection strategies and neural-selector feature ablations (**Supplementary Table 9**). Exhaustive Brier-based selection achieved higher chain-macro AP (0.6264) than selecting one model per pLM (0.6093) or the three best individual models (0.6155). Within the full neural selector, removing pairwise error features reduced performance (AP = 0.6210 to 0.6065 under mean-seed settings), whereas removing static features had little effect. Nevertheless, the individual-only strategy achieved the highest observed chain-macro AP (0.6312), although its Brier scores were higher than those of the full selector at matched seeds. Removing candidate masking yielded identical reported results, and beam search matched exhaustive search across mean-seed configurations. These findings suggest that more complex selection strategies do not consistently improve predictive performance and that the preferred configuration depends on the evaluation metric.

**Supplementary Note 5**

**Predictive complementarity contributes to CHORUS ensemble performance**

To investigate whether the performance improvement of CHORUS could be explained by averaging multiple predictions, diversifying a single representation, or increasing model capacity, we constructed several controlled alternatives (**Supplementary Fig. 8**). CHORUS achieved a chain-macro AP value of 0.6434 on the held-out test set, outperforming ensembles based on different training seeds (0.6210), orthogonally transformed embeddings (0.6178), distinct ESM2 representation layers (0.5881), an enlarged single predictor (0.5567), and six identical predictions (0.5752). All paired performance differences were in favour of CHORUS, with 95% confidence intervals excluding zero (**Supplementary Table 12**). A complementary analysis further revealed that the complementarity of predicted probabilities alone was insufficient to explain ensemble performance. For instance, the FD6 ensemble, comprising the GTB heads spanning three pLM families and two representation layers per family, achieved a chain-macro AP value of 0.6330 at a mean prediction correlation of 0.6409. In contrast, the ensemble constructed from different ESM2 layers exhibited substantially lower prediction correlation (0.4561) but also lower AP (0.5886) (**Extended Data Fig. 3a**; **Supplementary Table 11**). These findings indicate that ensemble performance depends not merely on the complementarity of predicted probabilities, but also on the predictive quality and complementary information captured by the individual models in CHORUS.

Ensemble performance was not monotonically associated with inter-model prediction correlation (**Supplementary Table 2**). Although the ESM2-layer ensemble exhibited the lowest mean correlation (0.4561), its ensemble AP remained relatively low (0.5886). In contrast, FD6 achieved the highest ensemble AP (0.6330) despite higher prediction correlation (0.6409) and lower mean individual-model AP than the training-seed ensemble. These findings suggest that effective ensemble integration depends on predictive complementarity rather than prediction decorrelation alone. CHORUS consistently outperformed ensembles constructed using different training seeds, orthogonally transformed embeddings or ESM2 representation layers, as well as an enlarged single predictor and an identical-prediction control. CHORUS achieved a chain-macro AP value of 0.6434, with improvements of 0.0224-0.0868 over these alternatives. All paired 95% confidence intervals for the performance gains excluded zero. These results indicate that the predictive advantage of CHORUS cannot be reproduced by model replication, representation reparameterization, layer variation or increased model capacity alone.

**Supplementary Note 6**

**Biological information and representation depth jointly track predictive performance**

To investigate whether layer-dependent differences in the performance of predicting interaction sites in membrane proteins reflect biologically informative pLM representations, we developed six linear probes capturing membrane protein topology, helix organization (helix position and membrane depth), intrachain packing (helix neighbours and local contacts), and residue conservation (**Supplementary Fig. 9**). In 16 pLM-layer settings (ESM2, ESMC, and ProtT5), biological probe profiles and layer depth jointly predicted downstream AP gains (cross-validated $R^{2}=0.821$; **Extended Data Fig. 5**), with transmembrane-specific features outperforming generic biological features. Further analyses revealed substantial overlap between biological information and layer depth, with shared effects accounting for most of the explained performance variation (**Supplementary Fig. 10**). Additionally, sensitivity and feature-level analyses supported these predictive associations but did not establish independent biological contributions to model performance (**Supplementary Fig. 11**).

**Representation diversity and individual-model complementarity**

We further examined whether representation similarity and functional diversity could predict performance gains from combining individual models. Across 171 model pairs, incorporating representation dissimilarity modestly improved the prediction of pairwise AP gains over baseline descriptors (leave-one-expert-out $R^{2}=0.611$ versus $0.597$), whereas functional-diversity features provided no consistent additional benefit (**Supplementary Fig. 11e**). These findings offer insights into model complementarity without establishing the mechanisms underlying the CHORUS performance.

**Supplementary Figures**

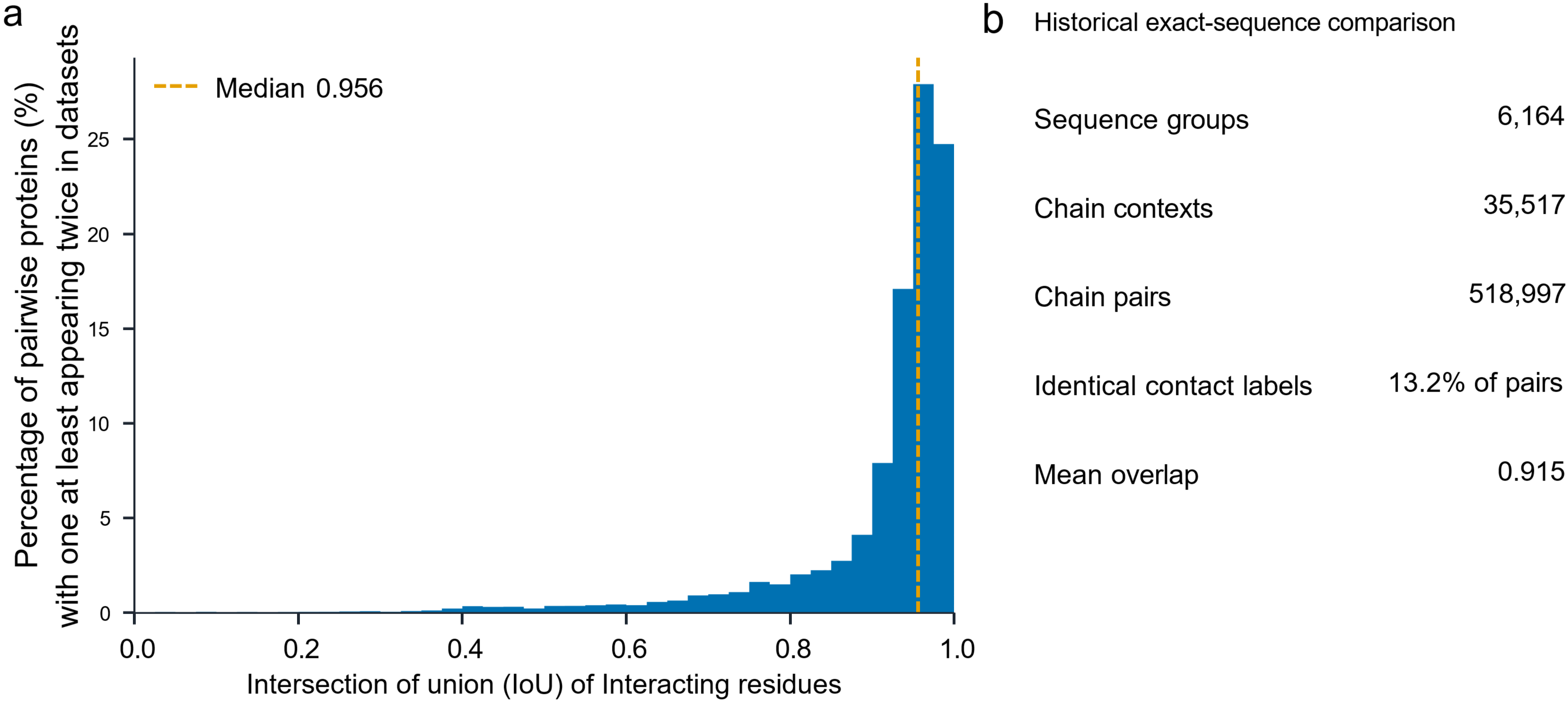

**Supplementary Fig. 1 |** **Variation in interacting residue positions among identical proteins across structural contexts. a,** Distribution of the overlap of interaction site labels (i.e., intersection over union, IoU) across 518,997 chain pairs (median IoU=0.956, dashed line). **b,** Summary of 6,164 sequence groups and 35,517 chain contexts. Only 13.2% of pairs exhibited identical interaction site labels (mean IoU=0.915). These data show that the same protein sequences appearing across different structural contexts (i.e., performing different cellular functions) often possess consistent interaction site labels. However, the low IoU profile indicates that some proteins exhibit different interaction patterns under different structural contexts.

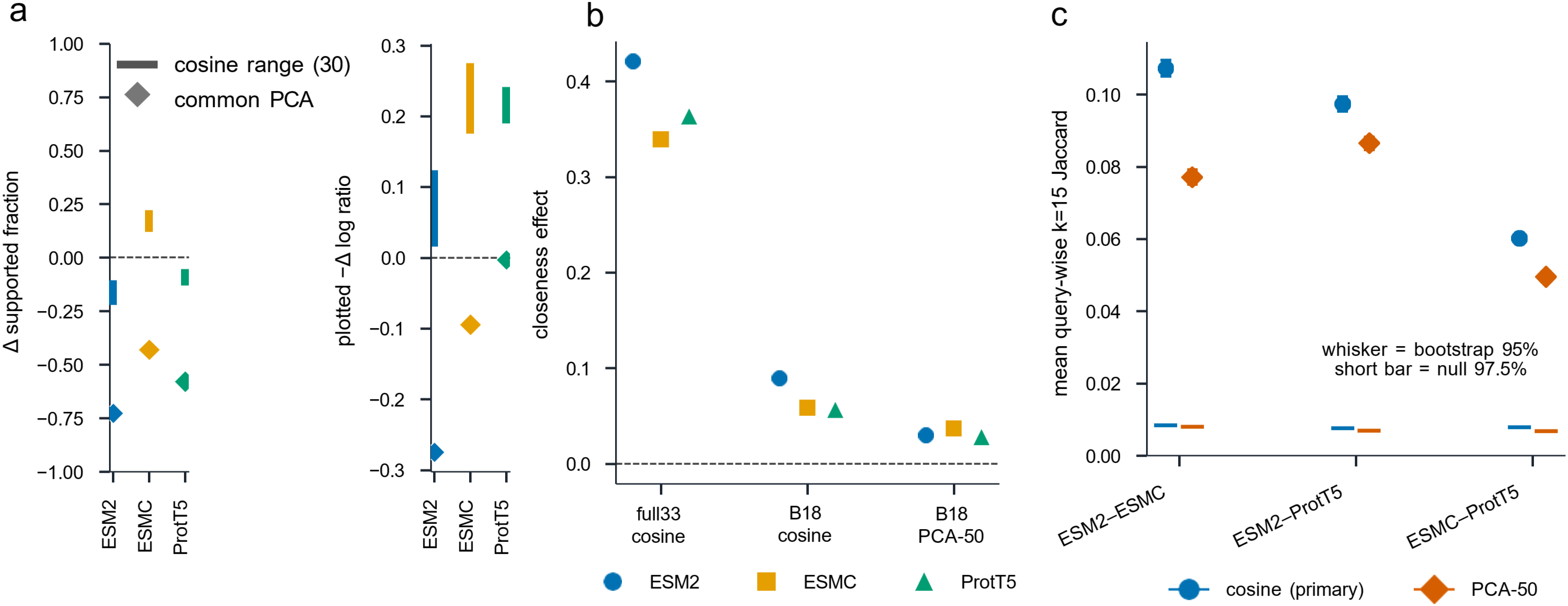

**Supplementary Fig. 2 | Reference-relative geometry of pLM representations.** Sequence-mean embeddings from ESM2, ESMC and ProtT5 were analyzed against 3,427 reference proteins, 33 membrane anchors, and 7,485 membrane-protein queries. **a,** Reciprocal neighborhood support measured by directional differences in supported fraction and log support ratio. **b,** Proximity to membrane anchors relative to matched null controls; positive values indicate closer proximity. **c,** Pairwise overlap of the 15 nearest reference neighbors across pLMs, with bootstrap 95% intervals and permutation-null thresholds.

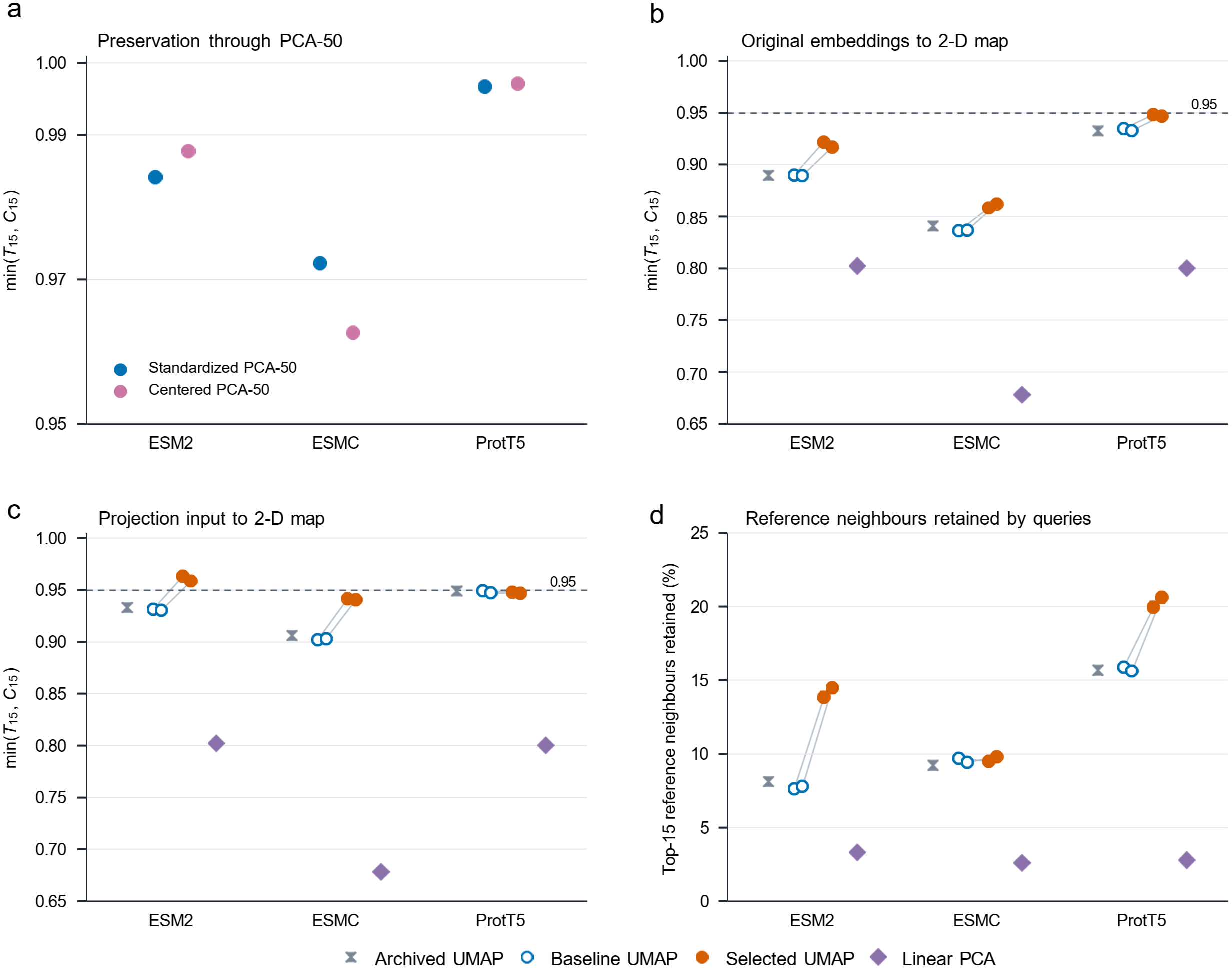

**Supplementary Fig. 3 | Projection fidelity and reference-neighbor retention. a,** Neighborhood preservation after standardized or centered PCA-50. **b,c,** Fidelity of two-dimensional projections relative to original embeddings (**b**) or projection inputs (**c**), measured by min(T15, C15), the lower of trustworthiness and continuity. Dashed lines indicate 0.95. **d,** Percentage of the 15 original reference neighbors retained for membrane-protein queries.

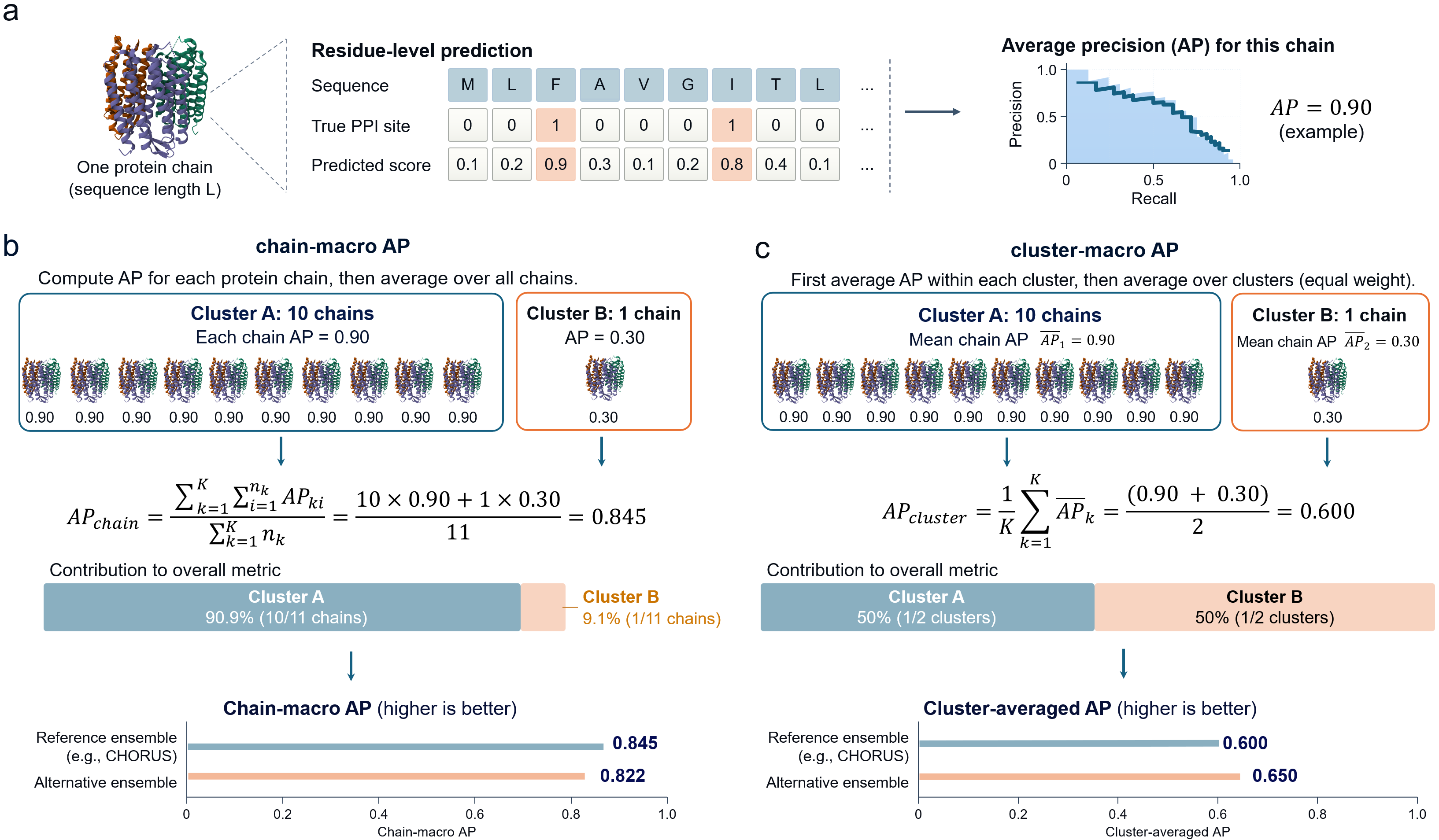

**Supplementary Fig. 4 |** **Schematic comparison of the chain-macro AP and cluster-averaged AP metrics.** **a,** Residue-level prediction scores are compared with binary interaction site labels to calculate average precision (AP) independently for each protein chain. **b,** Chain-macro AP is calculated by averaging chain-specific AP values across all eligible chains, assigning equal weight to each chain. Consequently, clusters containing more chains contribute proportionally more to the overall metric. **c,** Cluster-macro AP is calculated by first averaging chain-specific AP values within each cluster and then averaging across clusters, assigning equal weight to each cluster regardless of its size. The illustrative example contains two clusters comprising ten and one protein chains, respectively, yielding a chain-macro AP value of 0.845 and a cluster-macro AP value of 0.600. Comparison with an alternative ensemble demonstrates that these aggregation strategies can produce different model rankings.

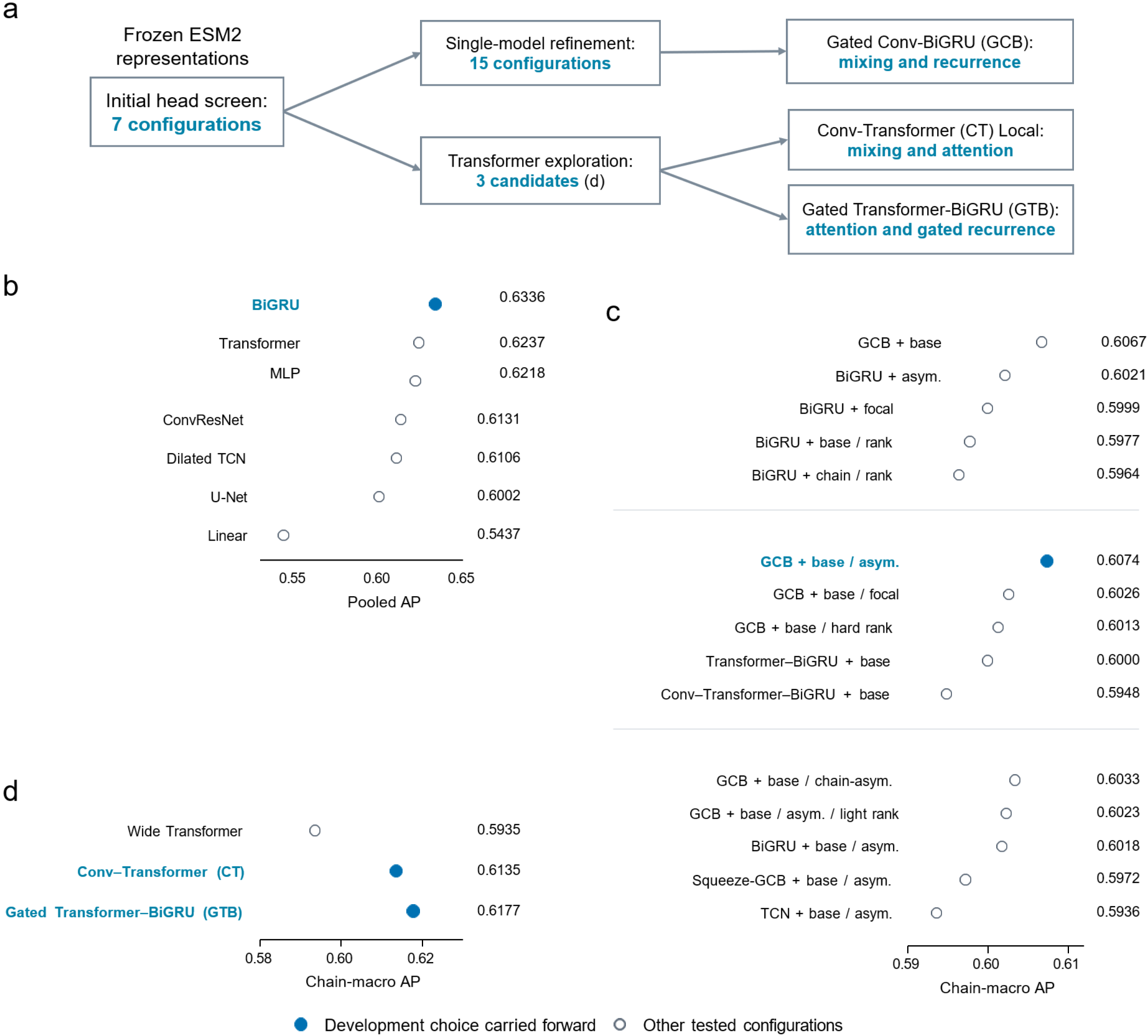

**Supplementary Fig. 5 | Exploration of optimal architectures for residue interaction prediction.** **a,** Workflow leading to gated Conv-BiGRU (GCB), Conv-Transformer (CT), and gated Transformer-BiGRU (GTB). Arrows indicate development lineage; the three Transformer candidates were trained in parallel. Intervening ensemble experiments are omitted. **b,** Seven heads evaluated using frozen ESM2 representations. Using the pooled average precision (AP) metric on validation data, BiGRU was identified as optimal. **c,** Fifteen configurations evaluated across three rounds of single-model refinement, each using a different seed. GCB with asymmetric focal loss was identified as the best configuration. The base term: a 28-dimensional non-MSA feature block containing topology-derived variables, subsequently replaced by ten sequence features. The asym term: asymmetric focal loss. The chain term: chain-balanced loss. The rank term: within-chain ranking. **d,** Comparison of Transformer, CT, and GTB using regenerated ESM2 layer-33 embeddings. CT and GTB were retained for subsequent architecture and layer screening. All benchmarks were made based on a strict-SI30 split containing 300 validation protein chains.

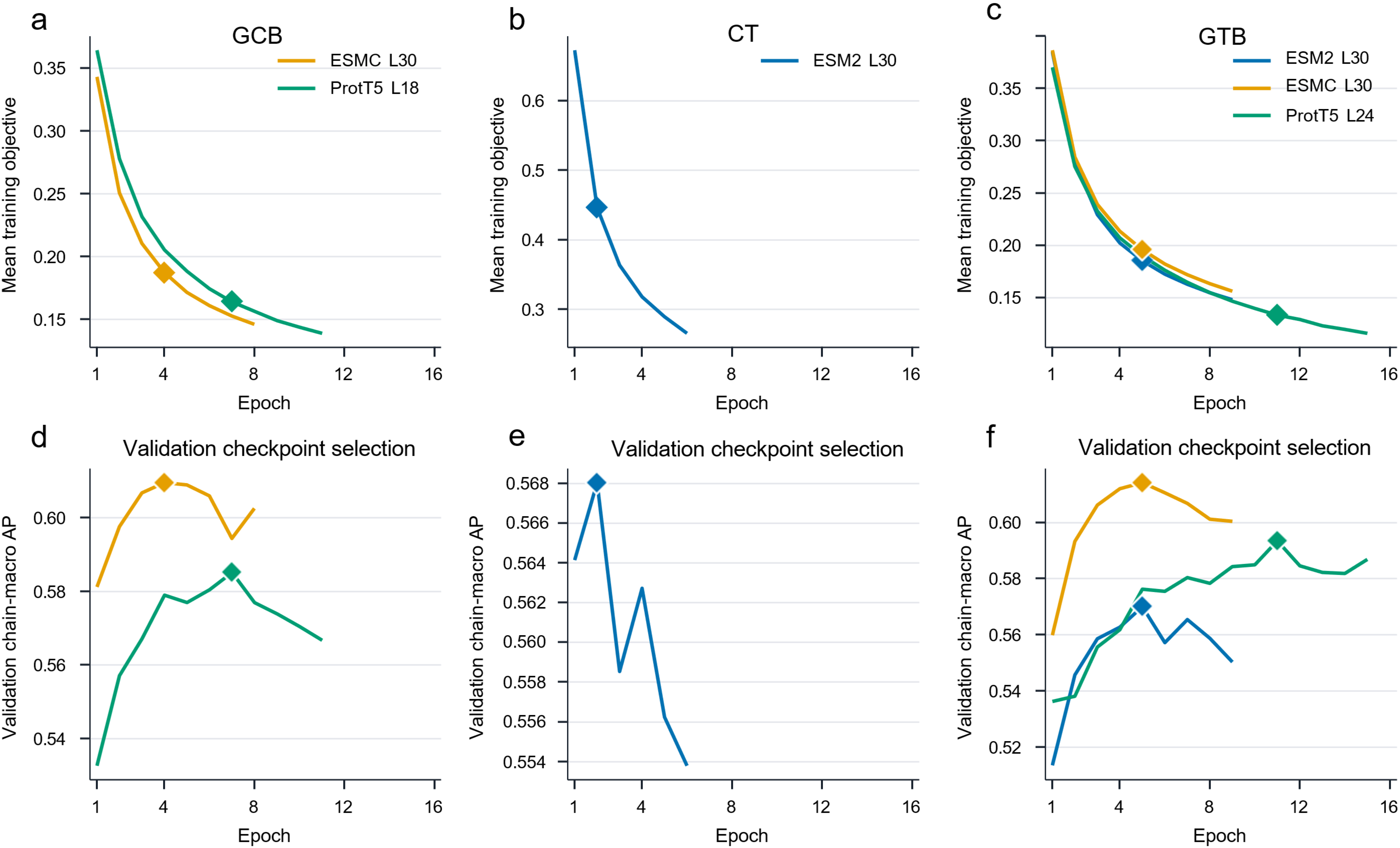

**Supplementary Fig. 6 | Training of the six individual models in CHORUS. a-c,** Mean training objectives for gated Conv-BiGRU (GCB), Conv-Transformer (CT) and gated Transformer-BiGRU (GTB). **d-f,** Corresponding validation chain-macro average precision (AP) across epochs. Colors indicate pLM families and representation layers. Diamonds mark checkpoints selected by the highest validation AP.

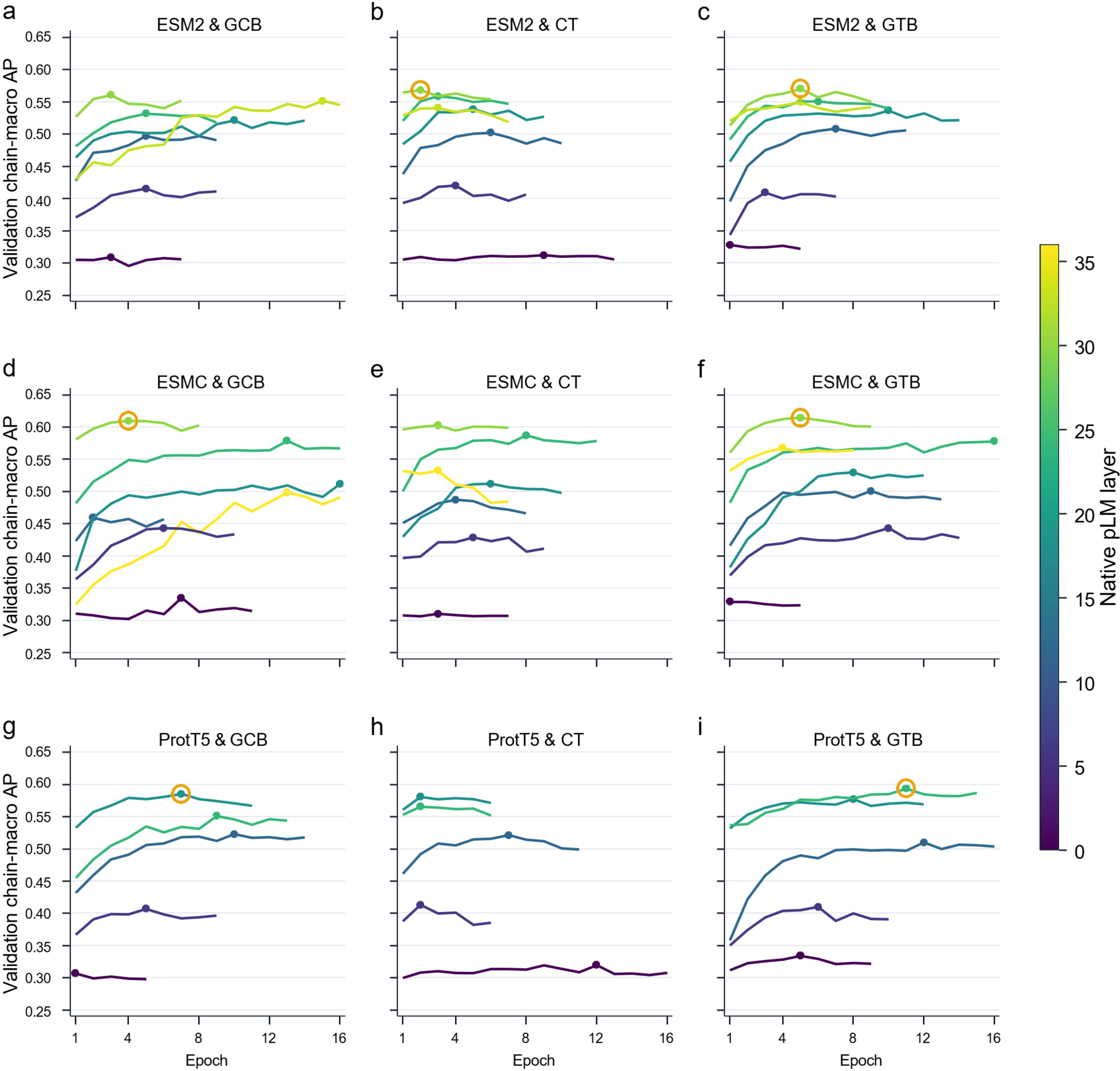

**Supplementary Fig. 7 | Validation of 57 candidate individual models with regard to training epochs. a-i,** Validation chain-macro AP across epochs for ESM2 (**a-c**), ESMC (**d-f**), and ProtT5 (**g-i**), each evaluated with GCB, CT and GTB heads. Each curve represents one candidate model, with colors indicating representation layers. Dots mark selected checkpoints, and orange rings identify the six individual models retained in CHORUS.

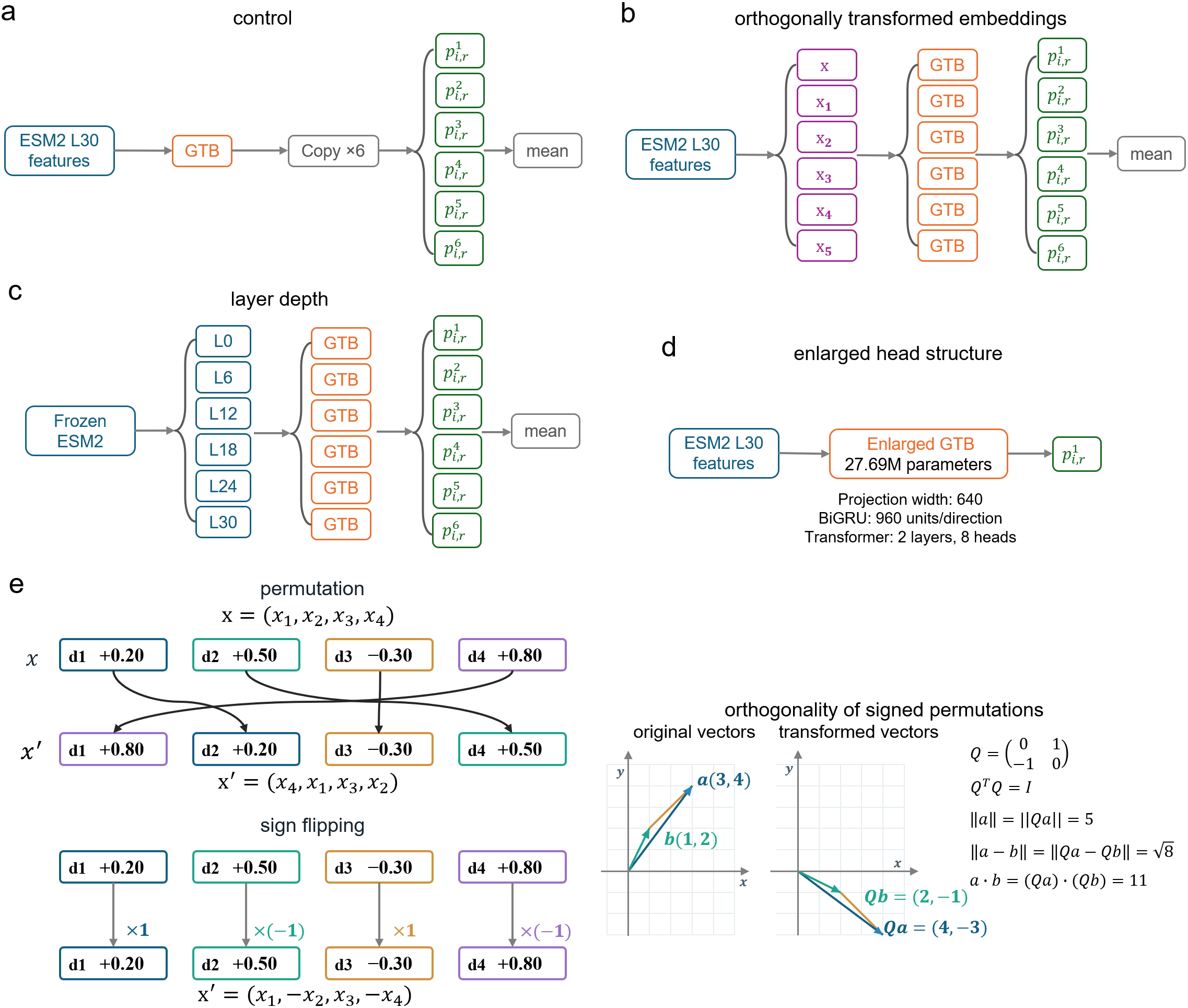

**Supplementary Fig. 8 |** **Construction of ensemble controls and orthogonal feature transformations.** **a,** Identical-prediction control in which a single ESM2 layer-30 GTB predictor is replicated six times before probability averaging, yielding an output identical to the original predictor. **b,** Transformed embeddings comprising six independently trained GTB models using the ESM2 layer-30 embeddings and five fixed signed permutations of the same representations. The transformation retains feature orthogonality. **c,** Representation-depth control, combining six GTB predictors trained on ESM2 representations extracted from layers 0, 6, 12, 18, 24 and 30. **d,** Enlarged single-model control using an ESM2 layer-30 GTB architecture with 27.69 million trainable downstream parameters, a projection width of 640, 960 BiGRU units per direction and two Transformer layers with eight attention heads. **e,** Illustration of feature transformation using two signed permutation techniques, including feature-dimension permutation and sign flipping, together with a two-dimensional example demonstrating that signed permutations are orthogonal transformations that preserve vector norms, pairwise Euclidean distances, and inner products. All six-predictor controls use unweighted averaging of predicted probabilities of interacting residues.

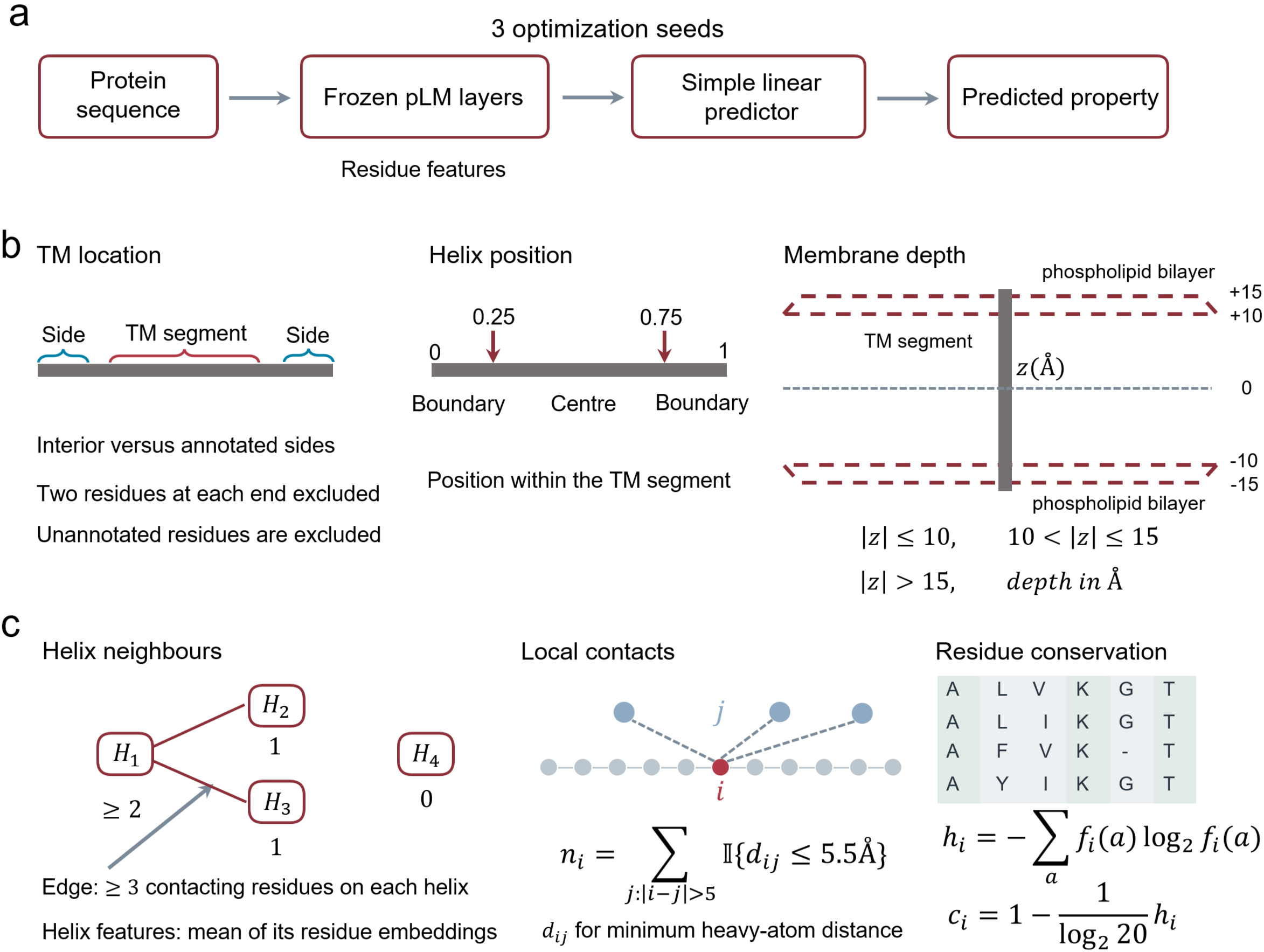

**Supplementary Fig. 9 | Biological probing of frozen pLM representations. a,** Schematic of linear probes trained on frozen pLM representations to recover independently derived biological properties. Probes were trained on training clusters and evaluated on validation clusters across three optimization seeds. **b,** Membrane protein-related properties, including membrane topologies, relative residue positions in helices, and membrane depth categorized by absolute distance from the membrane midplane ($\leq10$, $10-15$, and $>15Å$). **c,** Intrachain packing targets comprising TM helix-neighbour counts ($0, 1$ or$\geq2$) and nonlocal residue contacts defined by a minimum heavy-atom distance of $\leq5.5Å$ and sequence separation of $>5$ residues. Residue conservation scores were estimated from the entropy of query-aligned multiple sequence alignment (MSA) columns. Structural and MSA annotations were used exclusively to define probing properties and were not required for CHORUS inference.

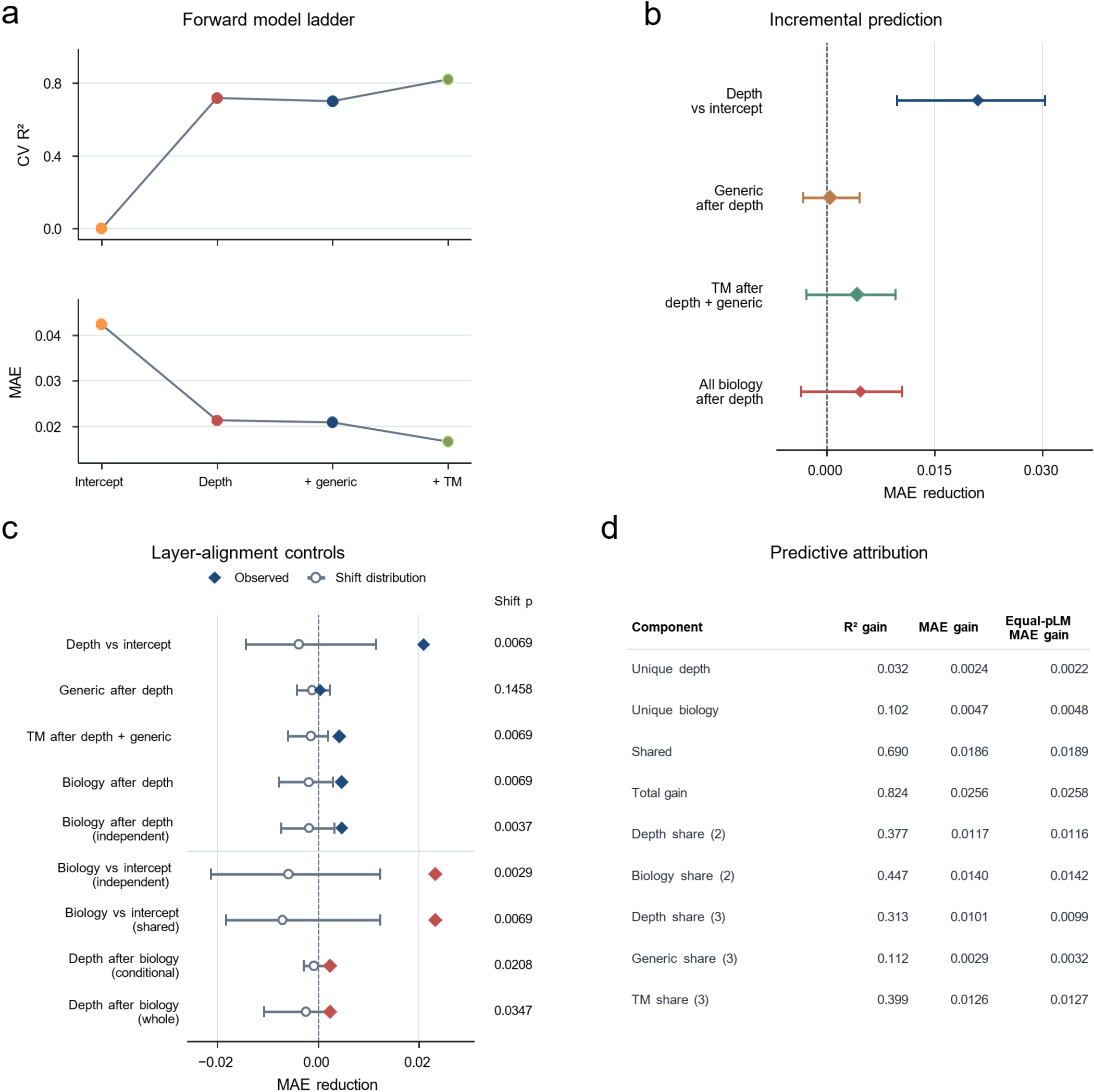

**Supplementary Fig. 10 | Incremental and shared predictive contributions of representation depth and biological information. a,** Cross-validated $R^{2}$ and MAE across a sequential model ladder incorporating layer depth, generic biology and TM-specific biology. **b,** Incremental MAE reductions associated with successive predictor additions. Diamonds indicate observed improvements, with 95% intervals from 2,000 cluster bootstrap resamples. **c,** Layer-shift controls comparing observed MAE reductions (diamonds) with null distributions generated by shifting biological profiles across layers (open circles and central 95% ranges). Reported P values quantify layer-specific alignment rather than independent predictive contributions. **d,** Commonality analysis and Shapley decomposition partitioning predictive improvements into unique and shared contributions of representation depth, generic biology and TM-specific biology. $R^{2}$ and MAE gains are calculated relative to the intercept-only model. All analyses describe retrospective predictive associations rather than causal effects.

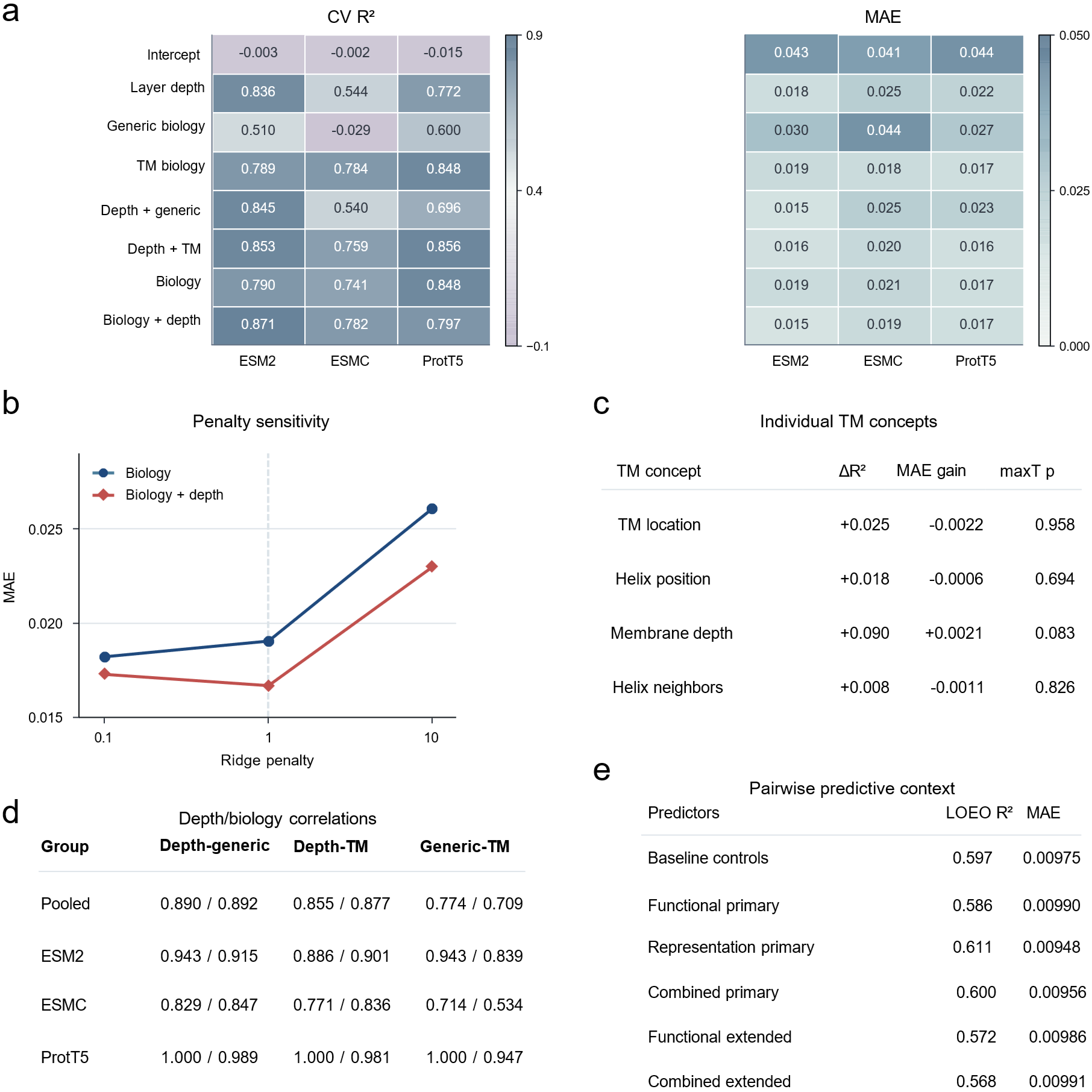

**Supplementary Fig. 11 | Sensitivity analyses of biological predictors and exploratory model complementarity. a,** Leave-one-pLM-out cross-validated $R^{2}$ and MAE for eight combinations of representation depth and biological predictors, evaluated separately across ESM2, ESMC and ProtT5. **b,** Sensitivity of predictive MAE to ridge regularization (penalties of 0.1, 1, and 10) for biological predictors with or without layer depth. **c,** Incremental $R^{2}$, MAE reduction and multiplicity-adjusted maxT P values for individual TM-related features added to the depth-plus-generic model. **d,** Spearman and Pearson correlations among representation depth, generic biology and TM-specific biological profiles, evaluated collectively and within each pLM family. **e,** Leave-one-expert-out prediction of pairwise AP gains across 171 combinations of 19 representation experts using baseline descriptors, functional-diversity metrics and representation-similarity features. Pairwise gains are defined relative to the better-performing individual expert. These exploratory comparisons do not establish the mechanisms underlying CHORUS ensemble performance.

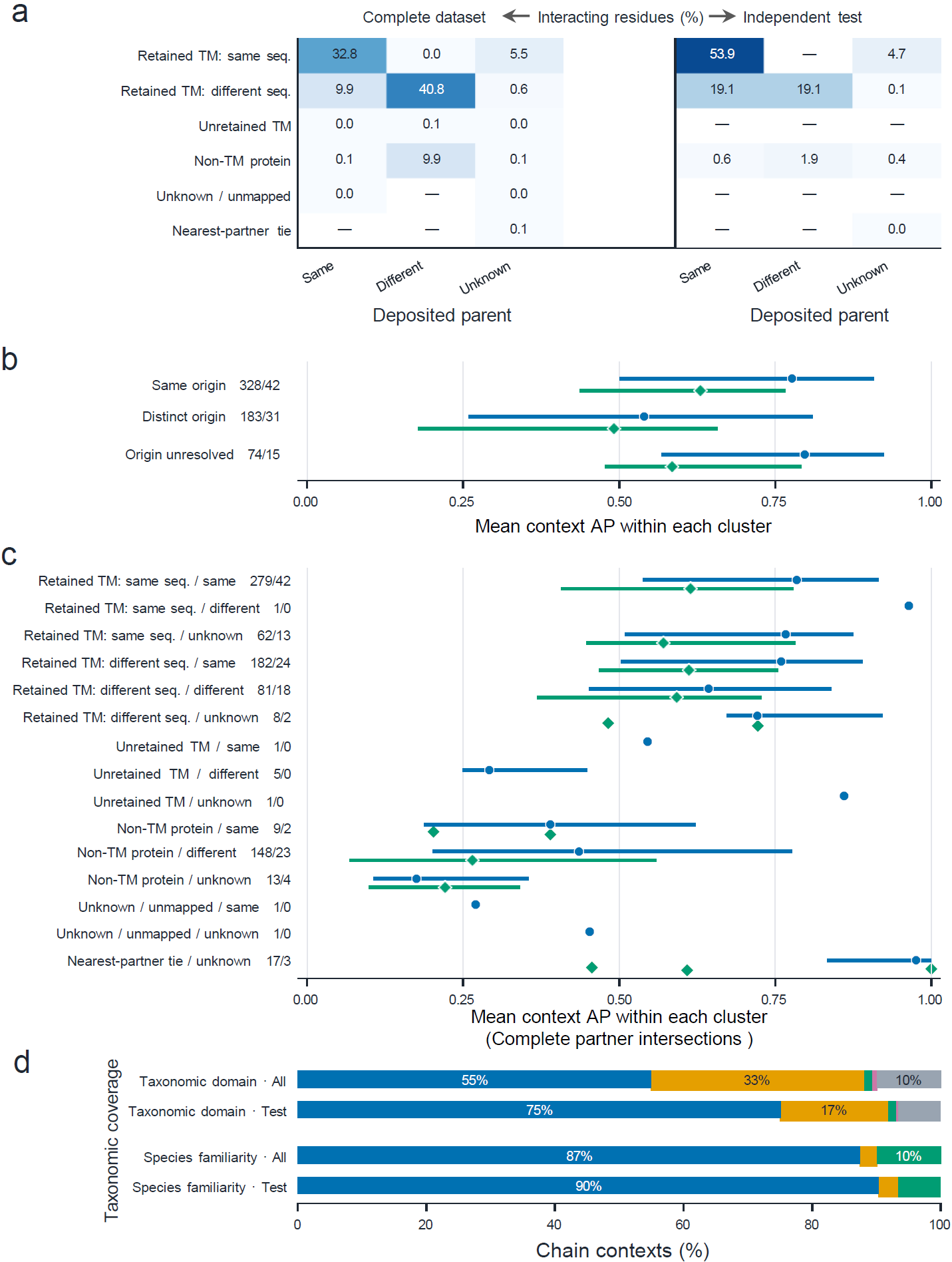

**Supplementary Fig. 12 | Biological composition of protein contacts and associated prediction performance. a,** Distribution of interacting residues by interacting-partner class and deposited protein-entity relationship in the complete dataset and independent test set. **b,c,** Chain-macro AP performance stratified by deposited parent-entity relationship (**b**) and its intersections with partner class (**c**). Points indicate medians and bars indicate interquartile ranges. Labels denote eligible cluster counts (blue for complete or green for independent test). **d,** Coverage of chain contexts by taxonomic domain and species representation during training.

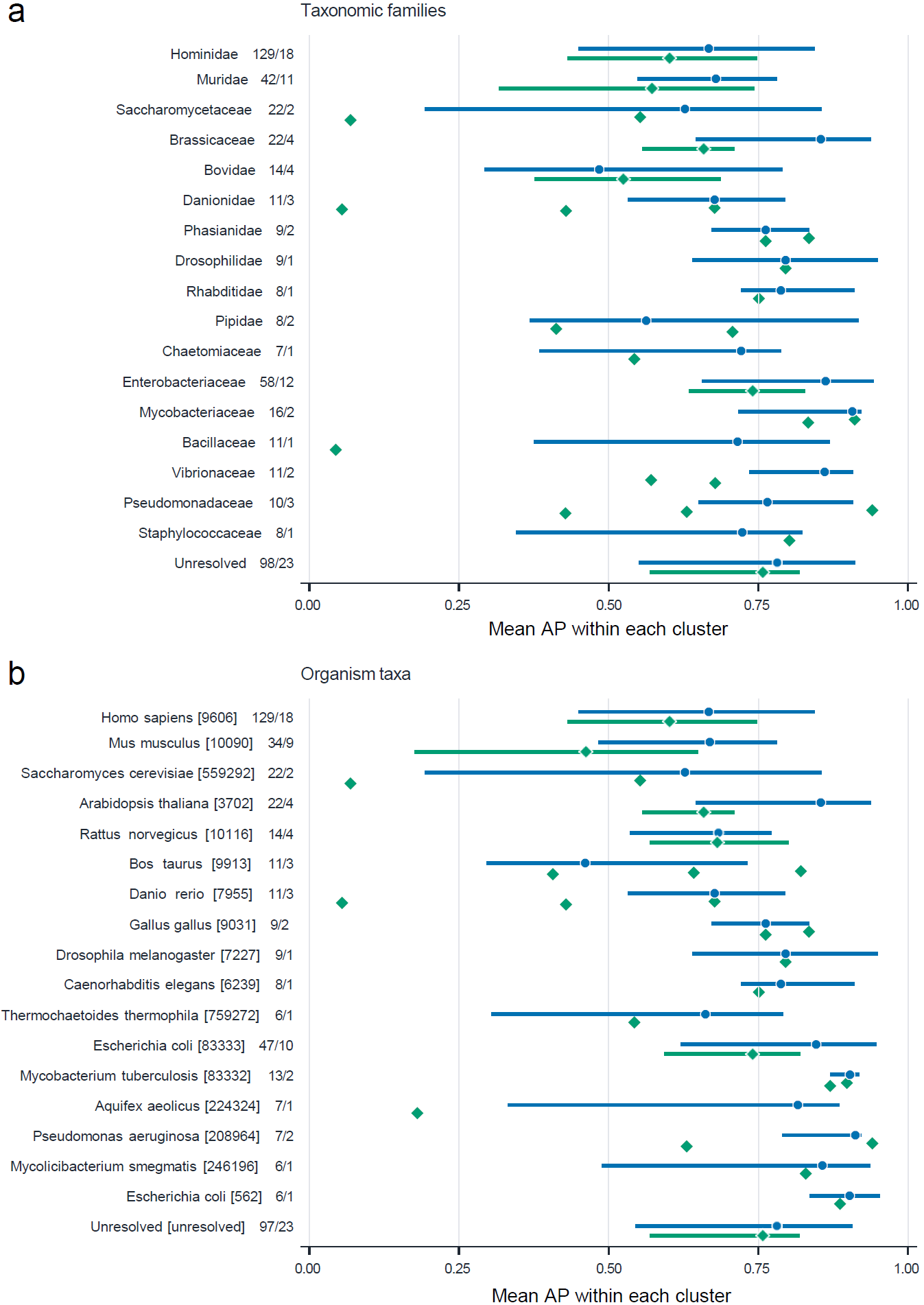

**Supplementary Fig. 13 | Taxonomic coverage and prediction performance. a,b,** Cluster-level average precision (AP) across 18 taxonomic families (**a**) and 18 organism taxa (**b**), selected by representation frequency independently of prediction performance. Blue and green indicate the complete dataset and independent test set, respectively. Points represent median cluster-averaged AP and bars indicate interquartile ranges; individual values are shown for groups with fewer than four eligible clusters. Labels indicate eligible cluster counts (complete/test). The complete dataset includes training, validation and test contexts and is shown for descriptive comparison.

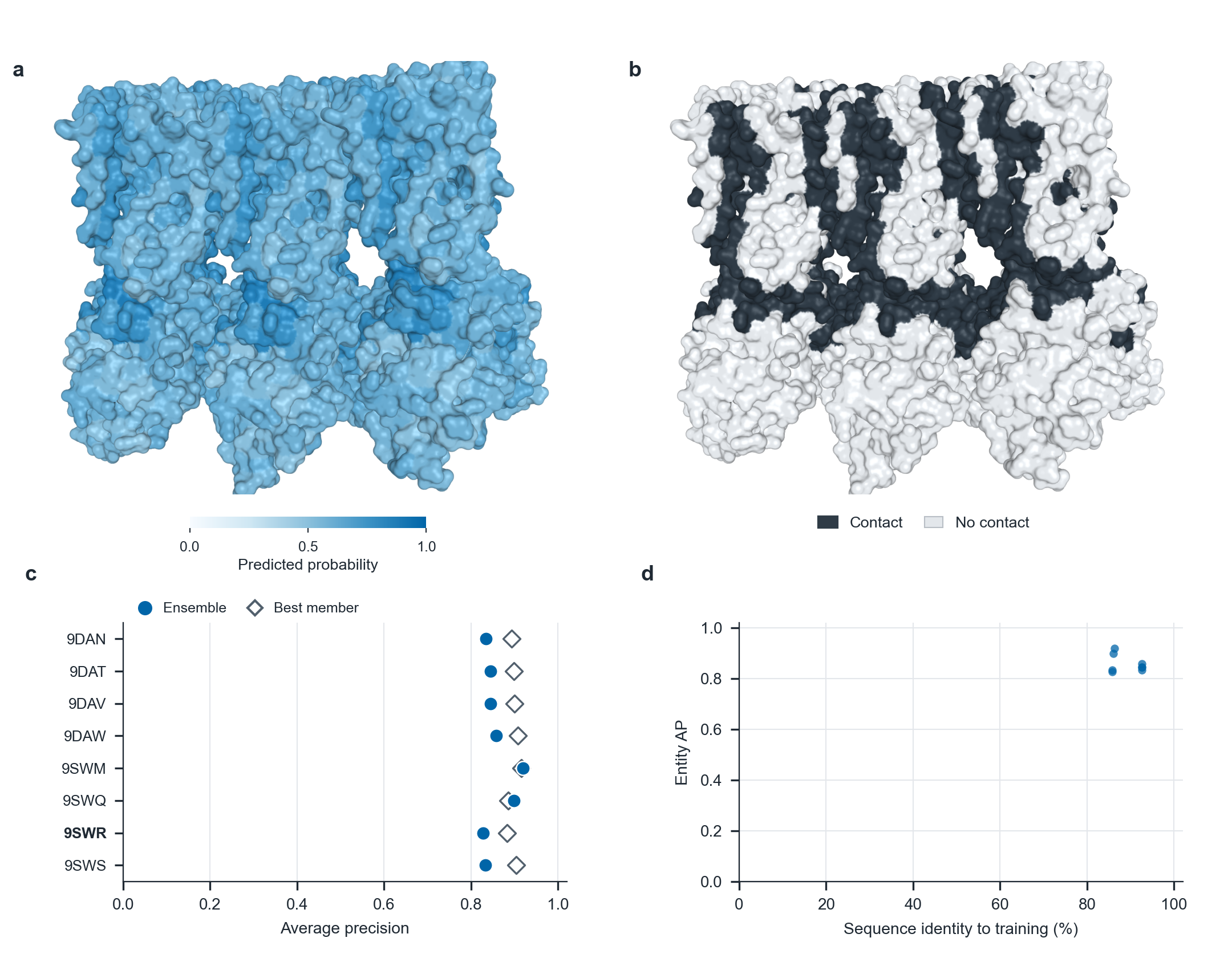

**Supplementary Fig. 14 | Analysis of interfaces on STING oligomers.** **a,b,** Structural representation of probabilities of residues predicted as interacting (**a**) and actual interaction interfaces colored in black using a distance cutoff of 5Å (**b**) on human STING hexamer (PDB code: 9SWR). The interaction partners are visualized in grey. **c,** Performance between CHORUS and its constituent models/members for eight STING structures. **d,** Entity AP against sequence identity to training proteins.

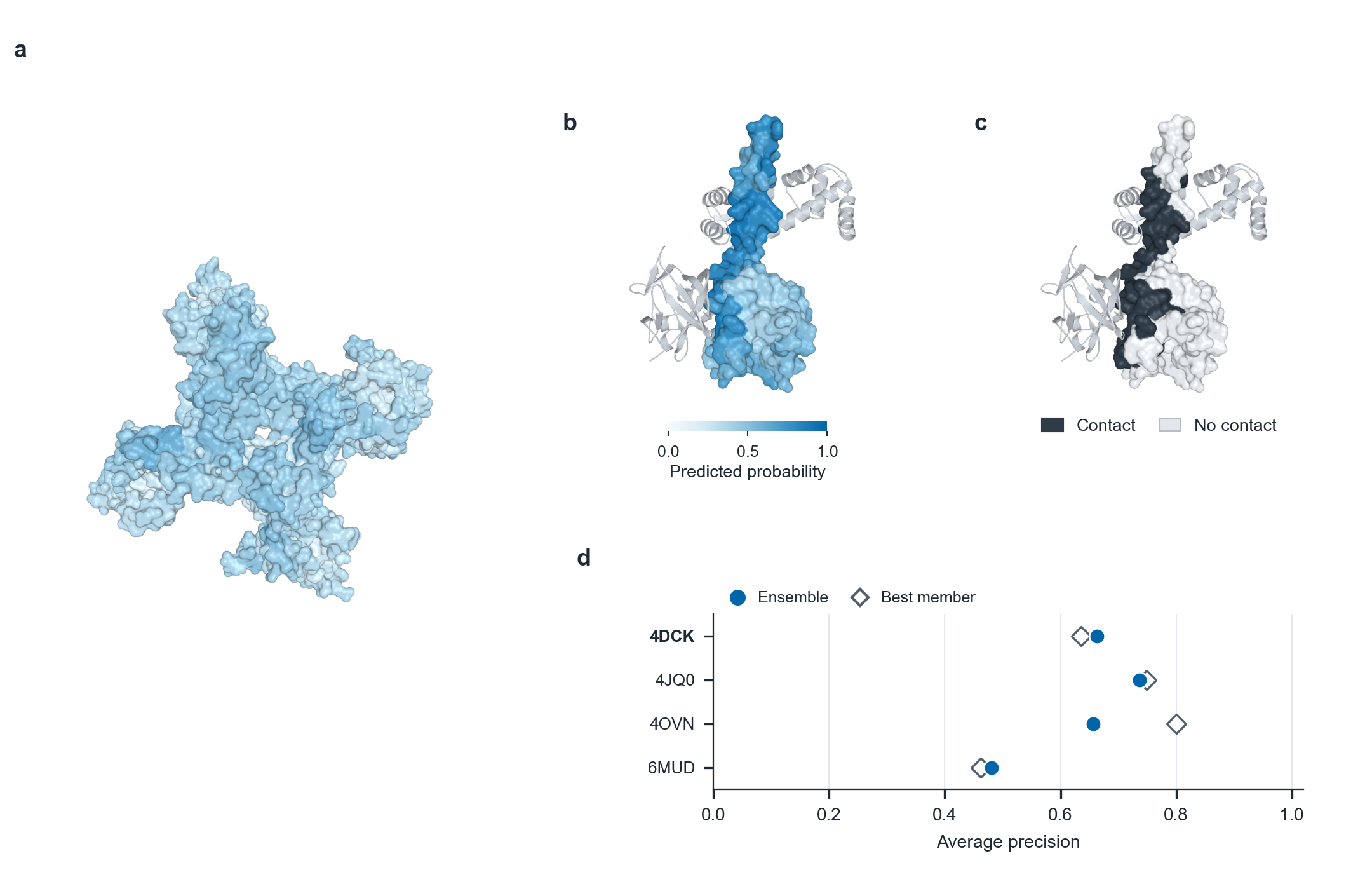

**Supplementary Fig. 15 | Analysis of interfaces on human cardiac sodium channel NaV1.5 (SCN5A).** **a,** Full channel structure (PDB code: 9P24) coloured by CHORUS-predicted probabilities. As a single-chain deposit, it has no interchain labels. **b,c,** Structural representation of probabilities of residues predicted as interacting (**b**) and actual interaction interfaces colored in black using a distance cutoff of 5Å (**c**) on C-terminal domain bound to FGF13 and calmodulin (PDB code: 4DCK). The interaction partner is visualized in grey. **d,** Performance between CHORUS and its constituent models/members for four C-terminal-domain complexes.

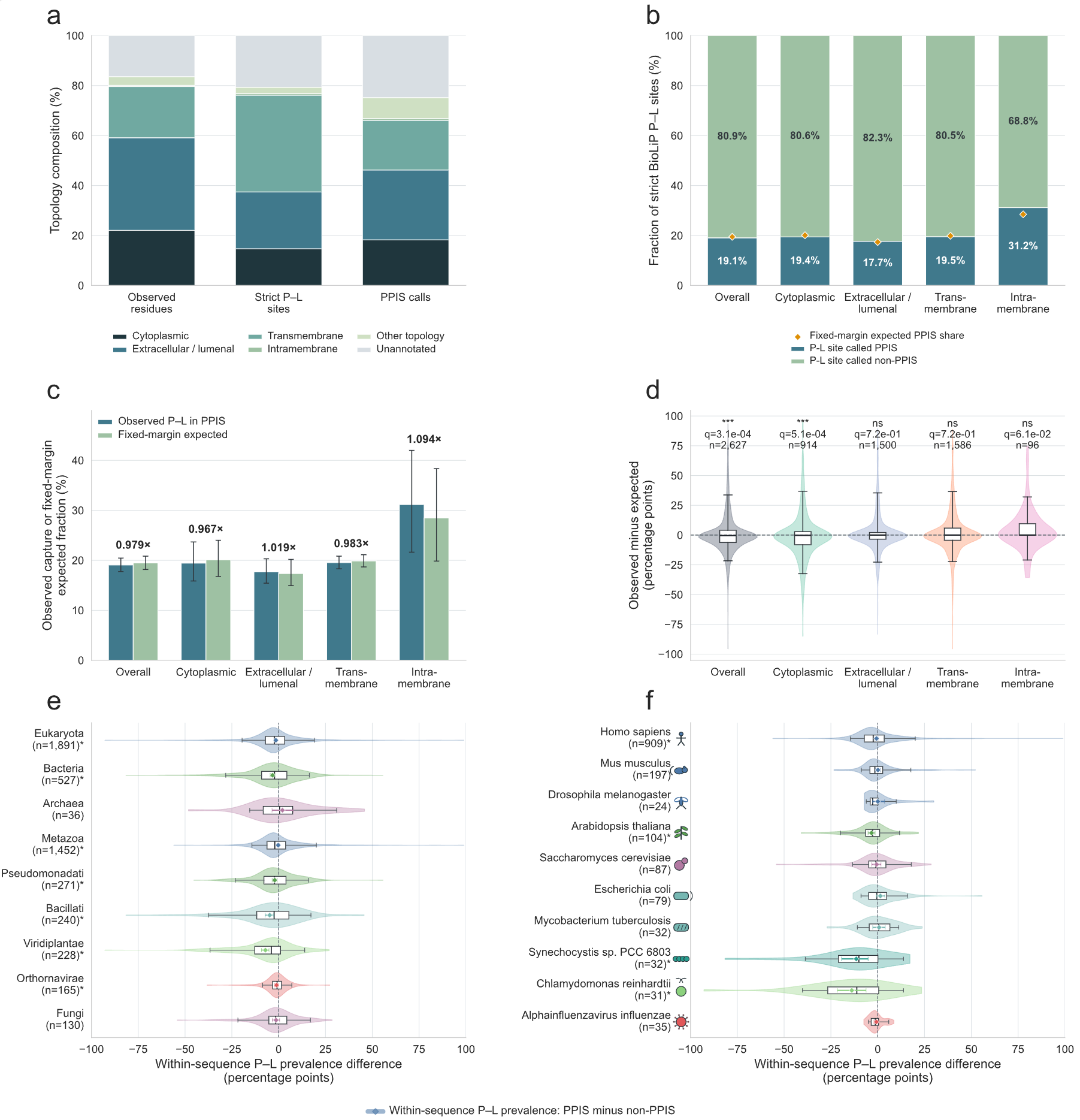

**Supplementary Fig. 16 | Analysis of relationships between predicted protein-protein interfaces and strict BioLiP protein-ligand-binding sites. a,** Structural topology composition. Proportion of mapped residues assigned to cytoplasmic, extracellular, transmembrane, intramembrane, other, or unannotated topologies across all evaluated positions, BioLiP-derived PLB sites, and predicted PPI sites. **b,** Fraction of PLB sites overlapping predicted PPI sites across membrane regions. Diamonds indicate fixed-margin expectations. **c,** Observed versus expected PLB-site capture by predicted PPI sites across membrane regions. Fold values indicate observed-to-expected ratios. Error bars depict 95% bootstrap confidence intervals. **d,** Sequence-level paired divergence. Paired percentage-point differences (observed minus expected capture fraction) across 2,627 exact sequences spanning the primary topology classes. Violin plots depict data distributions. Internal boxplots denote medians and interquartile ranges (IQR). False-discovery-rate (FDR)-adjusted $q$-values derive from paired two-sided Wilcoxon signed-rank tests. $n$ indicates the number of eligible sequences. **e,f,** Differential PLB-site occupancy across evolutionary lineages. For each protein, the occupancy was calculated as the difference between the fraction of predicted interacting residues annotated as PLB sites and that among non-interacting residues. A value of zero denotes equal PLB-site occupancy between predicted interacting and non-interacting residues, whereas positive and negative values indicate preferential occupancy within and outside interaction sites, respectively. Distributions are shown across major taxonomic strata (**e**) and selected representative species (**f**).

**Supplementary Tables**

**Supplementary Table 1** **|** Summary of datasets used to train CHORUS and valid as well as test its performance.

| **Partition** | **Contexts** | **Sequences** | **Clusters** | **Residues** | **Contacts** |
| --- | --- | --- | --- | --- | --- |
| Training | 30,675 | 8,483 | 299 | 7,106,648 | 2,279,484 |
| Validation | 5,546 | 1,538 | 65 | 2,244,041 | 523,268 |
| Independent test | 4,655 | 1,461 | 55 | 2,186,751 | 502,137 |

**Supplementary Table 2 |** Benchmark of various learning architectures to determine downstream heads for interaction site prediction. GCB: Conv-BiGRU. CT: Conv-Transformer. GTB: gated Transformer-BiGRU.

| **Configuration** | **Panel** | **Seed** | **Extra features** | **Loss** | **Rank weight** | **Val. AP** |
| --- | --- | --- | --- | --- | --- | --- |
| Linear | b | 0 | 0 | WBCE | 0.0 | 0.5437 |
| MLP | b | 0 | 0 | WBCE | 0.0 | 0.6218 |
| ConvResNet | b | 0 | 0 | WBCE | 0.0 | 0.6131 |
| Dilated TCN | b | 0 | 0 | WBCE | 0.0 | 0.6106 |
| U-Net | b | 0 | 0 | WBCE | 0.0 | 0.6002 |
| **BiGRU** | b | 0 | 0 | WBCE | 0.0 | **0.6336** |
| Transformer | b | 0 | 0 | WBCE | 0.0 | 0.6237 |
| BiGRU + focal | c | 1 | 0 | Focal | 0.0 | 0.5999 |
| BiGRU + asym. | c | 1 | 0 | Asym. | 0.0 | 0.6021 |
| BiGRU + chain / rank | c | 1 | 0 | Chain | 0.12 | 0.5964 |
| BiGRU + base / rank | c | 1 | 28 | WBCE | 0.12 | 0.5977 |
| GCB + base | c | 1 | 28 | Chain | 0.0 | 0.6067 |
| GCB + base / focal | c | 2 | 28 | Focal | 0.0 | 0.6026 |
| **GCB + base / asym.** | c | 2 | 28 | Asym. | 0.0 | **0.6074** |
| GCB + base / hard rank | c | 2 | 28 | Chain | 0.08 | 0.6013 |
| Transformer-BiGRU + base | c | 2 | 28 | WBCE | 0.0 | 0.6000 |
| CT-BiGRU + base | c | 2 | 28 | WBCE | 0.0 | 0.5948 |
| GCB + base / chain-asym. | c | 3 | 28 | Chain-asym. | 0.0 | 0.6033 |
| GCB + base / asym. / light rank | c | 3 | 28 | Asym. | 0.03 | 0.6023 |
| Squeeze-GCB + base / asym. | c | 3 | 28 | Asym. | 0.0 | 0.5972 |
| BiGRU + base / asym. | c | 3 | 28 | Asym. | 0.0 | 0.6018 |
| TCN + base / asym. | c | 3 | 28 | Asym. | 0.0 | 0.5936 |
| Wide Transformer | d | 0 | 0 | Chain | 0.0 | 0.5935 |
| **CT** | d | 0 | 0 | Chain | 0.05 | **0.6135** |
| **GTB** | d | 0 | 0 | Asym. | 0.03 | **0.6177** |

**Supplementary Table 3 |** Information about individual models in CHORUS.

| **ID** | **Combination** |
| --- | --- |
| 1 | ESM2 / GTB / L30 |
| 2 | ESM2 / CT / L30 |
| 3 | ESMC / GTB / L30 |
| 4 | ESMC / GCB / L30 |
| 5 | ProtT5 / GTB / L24 |
| 6 | ProtT5 / GCB / L18 |

**Supplementary Table 4 |** Comparison of performance using embeddings of different layers of pLMs across three neural network architectures (*i.e.*, heads/backbones).

| **pLM** | **Backbone** | **Native layer(s)** | **Input width** | **Epoch** | **Val. AP** | **Test AP** |
| --- | --- | --- | --- | --- | --- | --- |
| ESM2 | CT | 0 | 1,280 | 9 | 0.3119 | 0.3103 |
| ESM2 | CT | 6 | 1,280 | 4 | 0.4199 | 0.3998 |
| ESM2 | CT | 12 | 1,280 | 6 | 0.5020 | 0.4693 |
| ESM2 | CT | 18 | 1,280 | 5 | 0.5383 | 0.5277 |
| ESM2 | CT | 24 | 1,280 | 3 | 0.5584 | 0.5377 |
| **ESM2** | **CT** | **30** | **1,280** | **2** | **0.5680** | **0.5571** |
| ESM2 | CT | 33 | 1,280 | 3 | 0.5404 | 0.5304 |
| ESM2 | GCB | 0 | 1,290 | 3 | 0.3089 | 0.3183 |
| ESM2 | GCB | 6 | 1,290 | 5 | 0.4153 | 0.4074 |
| ESM2 | GCB | 12 | 1,290 | 5 | 0.4970 | 0.4442 |
| ESM2 | GCB | 18 | 1,290 | 10 | 0.5215 | 0.4901 |
| ESM2 | GCB | 24 | 1,290 | 5 | 0.5318 | 0.5238 |
| ESM2 | GCB | 30 | 1,290 | 3 | 0.5605 | 0.5500 |
| ESM2 | GCB | 33 | 1,290 | 15 | 0.5515 | 0.5343 |
| ESM2 | GTB | 0 | 1,280 | 1 | 0.3278 | 0.3328 |
| ESM2 | GTB | 6 | 1,280 | 3 | 0.4093 | 0.4220 |
| ESM2 | GTB | 12 | 1,280 | 7 | 0.5081 | 0.4838 |
| ESM2 | GTB | 18 | 1,280 | 10 | 0.5366 | 0.5080 |
| ESM2 | GTB | 24 | 1,280 | 6 | 0.5506 | 0.5452 |
| **ESM2** | **GTB** | **30** | **1,280** | **5** | **0.5701** | **0.5752** |
| ESM2 | GTB | 33 | 1,280 | 5 | 0.5493 | 0.5535 |
| ESMC | CT | 0 | 1,152 | 3 | 0.3099 | 0.3006 |
| ESMC | CT | 6 | 1,152 | 5 | 0.4284 | 0.4153 |
| ESMC | CT | 12 | 1,152 | 4 | 0.4870 | 0.4510 |
| ESMC | CT | 18 | 1,152 | 6 | 0.5117 | 0.4760 |
| ESMC | CT | 24 | 1,152 | 8 | 0.5871 | 0.5506 |
| ESMC | CT | 30 | 1,152 | 3 | 0.6024 | 0.5571 |
| ESMC | CT | 36 | 1,152 | 3 | 0.5320 | 0.5148 |
| ESMC | GCB | 0 | 1,162 | 7 | 0.3348 | 0.3201 |
| ESMC | GCB | 6 | 1,162 | 6 | 0.4431 | 0.4422 |
| ESMC | GCB | 12 | 1,162 | 2 | 0.4595 | 0.4580 |
| ESMC | GCB | 18 | 1,162 | 16 | 0.5118 | 0.4820 |
| ESMC | GCB | 24 | 1,162 | 13 | 0.5787 | 0.5451 |
| **ESMC** | **GCB** | **30** | **1,162** | **4** | **0.6095** | **0.5806** |
| ESMC | GCB | 36 | 1,162 | 13 | 0.4984 | 0.4743 |
| ESMC | GTB | 0 | 1,152 | 1 | 0.3290 | 0.3251 |
| ESMC | GTB | 6 | 1,152 | 10 | 0.4429 | 0.4238 |
| ESMC | GTB | 12 | 1,152 | 9 | 0.5007 | 0.4659 |
| ESMC | GTB | 18 | 1,152 | 8 | 0.5293 | 0.5061 |
| ESMC | GTB | 24 | 1,152 | 16 | 0.5778 | 0.5607 |
| **ESMC** | **GTB** | **30** | **1,152** | **5** | **0.6142** | **0.5545** |
| ESMC | GTB | 36 | 1,152 | 4 | 0.5675 | 0.5164 |
| ProtT5 | CT | 0 | 1,024 | 12 | 0.3197 | 0.2981 |
| ProtT5 | CT | 6 | 1,024 | 2 | 0.4128 | 0.4177 |
| ProtT5 | CT | 12 | 1,024 | 7 | 0.5210 | 0.4999 |
| ProtT5 | CT | 18 | 1,024 | 2 | 0.5806 | 0.5657 |
| ProtT5 | CT | 24 | 1,024 | 2 | 0.5654 | 0.5580 |
| ProtT5 | GCB | 0 | 1,034 | 1 | 0.3067 | 0.3071 |
| ProtT5 | GCB | 6 | 1,034 | 5 | 0.4072 | 0.4154 |
| ProtT5 | GCB | 12 | 1,034 | 10 | 0.5228 | 0.4987 |
| **ProtT5** | **GCB** | **18** | **1,034** | **7** | **0.5852** | **0.5554** |
| ProtT5 | GCB | 24 | 1,034 | 9 | 0.5511 | 0.5175 |
| ProtT5 | GTB | 0 | 1,024 | 5 | 0.3341 | 0.3443 |
| ProtT5 | GTB | 6 | 1,024 | 6 | 0.4097 | 0.4394 |
| ProtT5 | GTB | 12 | 1,024 | 12 | 0.5099 | 0.5011 |
| ProtT5 | GTB | 18 | 1,024 | 8 | 0.5771 | 0.5518 |
| **ProtT5** | **GTB** | **24** | **1,024** | **11** | **0.5935** | **0.5506** |
| ProtT5 | CT | 0,6,12,18,24 | 5,120 | 5 | 0.5669 | – |
| ProtT5 | CT | 6,12,18,24 | 4,096 | 3 | 0.5726 | – |
| ProtT5 | GCB | 0,6,12,18,24 | 5,130 | 9 | 0.5654 | – |
| ProtT5 | GCB | 6,12,18,24 | 4,106 | 5 | 0.5671 | – |
| ProtT5 | GTB | 0,6,12,18,24 | 5,120 | 4 | 0.5708 | – |
| ProtT5 | GTB | 6,12,18,24 | 4,096 | 8 | 0.5729 | – |

**Supplementary Table 5 |** Ablation experiments of individual models. Removed IDs are derived from Supplementary Table 2.

| **Removed IDs** | **Retained** | **Remaining AP** | **Full minus remaining [95% CI]** |
| --- | --- | --- | --- |
| 1 | 5 | 0.6380 | 0.0054 [0.0013, 0.0087] |
| 2 | 5 | 0.6407 | 0.0027 [-0.0037, 0.0103] |
| 3 | 5 | 0.6417 | 0.0018 [-0.0005, 0.0049] |
| 4 | 5 | 0.6361 | 0.0073 [0.0040, 0.0098] |
| 5 | 5 | 0.6352 | 0.0082 [0.0035, 0.0120] |
| 6 | 5 | 0.6393 | 0.0042 [-0.0058, 0.0117] |
| 1, 2 (ESM2 family) | 4 | 0.6273 | 0.0161 [0.0088, 0.0236] |
| 3, 4 (ESMC family) | 4 | 0.6281 | 0.0154 [0.0101, 0.0214] |
| 5, 6 (ProtT5 family) | 4 | 0.6286 | 0.0149 [-0.0003, 0.0270] |

**Supplementary Table 6 |** Predictive complementarity of pairwise models in CHORUS. For each model pair, predicted probabilities were averaged with equal weights. The resulting chain-macro AP value was compared with that of the better-performing constituent model. The improvement was quantified as the difference between pairwise ensemble AP and the individual-model AP.

| **Pair** | **Selected model** | **Correlation** | **Pair AP** | **AP gain [95% CI]** |
| --- | --- | --- | --- | --- |
| 1 + 2 | 1 | 0.7177 | 0.5908 | 0.0155 [0.0040, 0.0290] |
| 1 + 3 | 3 | 0.6009 | 0.6056 | 0.0510 [0.0295, 0.0677] |
| 1 + 4 | 4 | 0.6286 | 0.6226 | 0.0419 [0.0254, 0.0545] |
| 1 + 5 | 5 | 0.5931 | 0.6115 | 0.0609 [0.0387, 0.0730] |
| 1 + 6 | 6 | 0.6407 | 0.6027 | 0.0472 [0.0392, 0.0553] |
| 2 + 3 | 3 | 0.6133 | 0.5990 | 0.0445 [0.0286, 0.0616] |
| 2 + 4 | 4 | 0.6208 | 0.6131 | 0.0325 [0.0204, 0.0473] |
| 2 + 5 | 5 | 0.5690 | 0.5997 | 0.0490 [0.0309, 0.0668] |
| 2 + 6 | 6 | 0.6235 | 0.5971 | 0.0417 [0.0304, 0.0547] |
| 3 + 4 | 3 | 0.8146 | 0.5904 | 0.0359 [0.0241, 0.0439] |
| 3 + 5 | 3 | 0.5850 | 0.5982 | 0.0437 [0.0250, 0.0578] |
| 3 + 6 | 3 | 0.6387 | 0.5957 | 0.0412 [0.0164, 0.0617] |
| 4 + 5 | 4 | 0.6139 | 0.6138 | 0.0331 [0.0225, 0.0410] |
| 4 + 6 | 4 | 0.6815 | 0.6052 | 0.0246 [0.0085, 0.0368] |
| 5 + 6 | 5 | 0.6287 | 0.5951 | 0.0445 [0.0218, 0.0561] |

**Supplementary Table 7 |** Definitions of model selection strategies.

| **Method** | **Modification relative to the full scorer** | **Purpose** |
| --- | --- | --- |
| One per pLM | Selects the candidate with the lowest cluster-averaged Brier score within each pLM family (ESM2, ESMC and ProtT5), yielding a three-model ensemble. | Assesses selection constrained to one model per pLM family. |
| Top-3 Brier | Selects the three candidates with the lowest individual-model cluster-averaged Brier scores, irrespective of pLM family. | Provides a baseline based solely on individual-model performance rankings. |
| Exhaustive Brier | Evaluates all eligible three-model combinations and selects the combination whose averaged predictions yield the lowest cluster-averaged Brier score. | Provides an exhaustive-search benchmark for optimizing joint ensemble performance. |
| Individual + pair | Retains all input features, individual and pairwise scoring terms, and candidate masking. | Serves as the reference configuration. |
| No candidate mask | Disables random candidate masking during training. | Assesses the contribution of candidate masking to selection performance. |
| No pair features | Removes explicit pairwise error features while retaining pairwise scoring. | Assesses the value of explicitly representing error relationships between candidate models. |
| No static | Removes model-configuration descriptors while retaining error-derived features. | Assesses whether error-derived information is sufficient without model-configuration information. |
| Static only | Removes individual-model error statistics and explicit pairwise error features while retaining model descriptors and pairwise scoring. | Assesses selection based solely on model-configuration information. |
| Individual only | Removes explicit pairwise features and pairwise scoring terms. | Assesses the contribution of modeling interactions between candidate models beyond their individual performance. |

**Supplementary Table 8** **|** Search strategies for ensemble model selection.

| **Search** | **Search** **strategy** | **Note** |
| --- | --- | --- |
| Exhaustive | Exhaustively evaluates all 39,711 possible model combinations. | Identifies the globally optimal combination under the specified scoring function. |
| Beam | Explores a restricted subset of promising model combinations. | Reduces computational cost but does not guarantee global optimality. Using a selector trained using neural network. |
| Direct | Selects models according to a predefined selection rule. | Does not require combinatorial search using the neural selector. |

**Supplementary Table 9** | Performance comparison of model-selection strategies and feature ablations. The number of distinct models corresponds to the number of models that overlap among the three choices within each of the 5 folds.

| **ID** | **Selection strategy** | **Seed** | **Search** | **Number of distinct models** | **Cluster AP** | **Cluster Brier** | **Chain AP** |
| --- | --- | --- | --- | --- | --- | --- | --- |
| **S01** | **Exhaustive training with Brier search** | **–** | **Direct** | **5** | **0.5712** | **0.1700** | **0.6264** |
| S02 | Individual + pair | 15 | Exhaustive | 6 | 0.5622 | 0.1693 | 0.6191 |
| S03 | Individual + pair | 16 | Exhaustive | 6 | 0.5655 | 0.1694 | 0.6224 |
| S04 | Individual + pair | 17 | Exhaustive | 6 | 0.5645 | 0.1698 | 0.6199 |
| **S05** | **Individual + pair** | **Mean** | **Beam** | **4** | **0.5637** | **0.1708** | **0.6210** |
| S06 | Individual + pair | Mean | Exhaustive | 4 | 0.5637 | 0.1708 | 0.6210 |
| S07 | No candidate mask | 15 | Exhaustive | 6 | 0.5622 | 0.1693 | 0.6191 |
| S08 | No candidate mask | 16 | Exhaustive | 6 | 0.5655 | 0.1694 | 0.6224 |
| S09 | No candidate mask | 17 | Exhaustive | 6 | 0.5645 | 0.1698 | 0.6199 |
| S10 | No candidate mask | Mean | Beam | 4 | 0.5637 | 0.1708 | 0.6210 |
| S11 | No candidate mask | Mean | Exhaustive | 4 | 0.5637 | 0.1708 | 0.6210 |
| S12 | No pair features | 15 | Exhaustive | 8 | 0.5619 | 0.1719 | 0.6165 |
| S13 | No pair features | 16 | Exhaustive | 5 | 0.5570 | 0.1745 | 0.6086 |
| S14 | No pair features | 17 | Exhaustive | 6 | 0.5531 | 0.1715 | 0.6125 |
| S15 | No pair features | Mean | Beam | 6 | 0.5552 | 0.1735 | 0.6065 |
| S16 | No pair features | Mean | Exhaustive | 6 | 0.5552 | 0.1735 | 0.6065 |
| S17 | No static features | 15 | Exhaustive | 7 | 0.5669 | 0.1685 | 0.6220 |
| S18 | No static features | 16 | Exhaustive | 6 | 0.5662 | 0.1712 | 0.6222 |
| S19 | No static features | 17 | Exhaustive | 5 | 0.5580 | 0.1717 | 0.6170 |
| S20 | No static features | Mean | Beam | 5 | 0.5632 | 0.1710 | 0.6209 |
| S21 | No static features | Mean | Exhaustive | 5 | 0.5632 | 0.1710 | 0.6209 |
| **S22** | **One per pLM** | **–** | **Direct** | **3** | **0.5568** | **0.1756** | **0.6093** |
| S23 | Static features only | 15 | Exhaustive | 5 | 0.5703 | 0.1740 | 0.6263 |
| S24 | Static features only | 16 | Exhaustive | 5 | 0.5618 | 0.1761 | 0.6210 |
| S25 | Static features only | 17 | Exhaustive | 5 | 0.5748 | 0.1757 | 0.6298 |
| S26 | Static features only | Mean | Beam | 4 | 0.5716 | 0.1744 | 0.6242 |
| S27 | Static features only | Mean | Exhaustive | 4 | 0.5716 | 0.1744 | 0.6242 |
| S28 | Top 3 individual Brier | – | Direct | 5 | 0.5603 | 0.1737 | 0.6155 |
| S29 | Individual only | 15 | Exhaustive | 6 | 0.5716 | 0.1717 | 0.6263 |
| S30 | Individual only | 16 | Exhaustive | 8 | 0.5728 | 0.1713 | 0.6308 |
| S31 | Individual only | 17 | Exhaustive | 6 | 0.5743 | 0.1738 | 0.6312 |
| **S32** | **Individual only** | **Mean** | **Beam** | **4** | **0.5751** | **0.1731** | **0.6267** |
| S33 | Individual only | Mean | Exhaustive | 4 | 0.5751 | 0.1731 | 0.6267 |

**Supplementary Table 10** | Performance comparison of model-selection strategies and feature ablations.

| **Rule** | **Validation ΔAP [95% CI]** | **Test ΔAP [95% CI]** | **Test ΔBrier (balanced)** |
| --- | --- | --- | --- |
| Random + one-swap | −0.0052 [−0.0222, +0.0065] | −0.0030 [−0.0126, +0.0134] | +0.0146 (worse) |
| CJ-MTR-1 | +0.0039 [−0.0009, +0.0089] | +0.0050 [−0.0027, +0.0115] | +0.0057 (worse) |
| RESS-18 | +0.0006 [−0.0064, +0.0067] | +0.0025 [−0.0076, +0.0150] | +0.0056 (worse) |

**Supplementary Table 11 |** Performance of various ensemble models.

| **Ensemble** | **Mean correlation** | **Mean AP** | **Ensemble AP** |
| --- | --- | --- | --- |
| FD6 | 0.6409 | 0.5564 | 0.6330 |
| Training seeds | 0.7531 | 0.5697 | 0.6211 |
| Orthogonally transformed embeddings | 0.7613 | 0.5681 | 0.6177 |
| ESM2 layers | 0.4561 | 0.4770 | 0.5886 |
| Control | 1.0000 | 0.5749 | 0.5749 |

**Supplementary Table 12** **|** Comparison of predictive performance between CHORUS and those ensemble models built upon altered training seeds, different layer choices, orthogonally transformed embeddings, and enlarged model sizes. Control represents selected one model copied 6 times.

| **Predictor** | **Chain-macro AP [95% CI]** | **CHORUS gain [95% CI]** |
| --- | --- | --- |
| CHORUS | 0.6434 [0.5804, 0.7127] | — |
| Training seeds | 0.6210 [0.5584, 0.6931] | 0.0224 [0.0041, 0.0329] |
| Orthogonally transformed embeddings | 0.6178 [0.5523, 0.6935] | 0.0256 [0.0044, 0.0399] |
| ESM2 layers | 0.5881 [0.5206, 0.6678] | 0.0553 [0.0362, 0.0667] |
| Enlarged model structure | 0.5567 [0.4747, 0.6523] | 0.0868 [0.0428, 0.1187] |
| Control | 0.5752 [0.5136, 0.6511] | 0.0682 [0.0434, 0.0837] |

**Supplementary Table 13** | Benchmark implementations and evaluated inputs.

| **Predictor** | **Implementation and input qualification** | **Structure inputs evaluated** |
| --- | --- | --- |
| ScanNet-MSA^1^ | Released checkpoint with shared-source A3M sensitivity | Experimental, ESMFold, ColabFold |
| GPSite^2^ | Released five-fold ensemble with shared-source ProtT5/ESMFold | ESMFold |
| PeSTo^3^ | Released checkpoint with protein-interface output channel with corrected residue export | Experimental, ESMFold, ColabFold |
| Seq-InSite^4^ | Released checkpoint | No target coordinates |
| PIPENN-EMB^5^ | Project-trained embedding-compatible implementation; not the released PPI ensemble | No target coordinates |
| ScanNet-noMSA^1^ | Released checkpoint without MSAs | Experimental, ESMFold, ColabFold |
| ALLSites^6^ | Released architecture retrained with ESM2-3B layer-36 features; checkpoint selected by pooled validation AP | No target coordinates |
| GraphPPIS-slow^7^ | Released checkpoint with shared-source PSSM/HHM sensitivity | Experimental, ESMFold, ColabFold |
| EquiPPIS^8^ | Released checkpoint with shared-source input | Experimental, ESMFold, ColabFold |
| Gated-GPS^9^ | Project mean of five released folds with shared-source input | Experimental, ESMFold, ColabFold |
| RGN^10^ | Released checkpoint with shared-source input | Experimental, ESMFold, ColabFold |
| GraphPPIS-fast^7^ | Released checkpoint and structure-only | Experimental, ESMFold, ColabFold |
| Spatom^11^ | Released checkpoint with shared-source input | Experimental, ESMFold, ColabFold |
| DeepTMInter^12^ | Released package v0.0.1 with MSA-derived and coevolutionary features; PDBTM topology replaces Phobius | No target coordinates |
| EnsemPPIS^13^ | Released checkpoint with shared-source ProtBERT1024 | No target coordinates |
| DeepPPISP^14^ | project-retrained recovered architecture; not official weights | No target coordinates |
| PITHIA^15^ | Released checkpoint with input/output compatibility fix | No target coordinates |
| DeepProSite^16^ | Released five-fold mean with shared-source inputs; partial coverage | ESMFold |
| MaSIF-site^17^ | Released all_feat_3l with target-only structure and partial surface coverage | Experimental |
| EGRET^18^ | Released checkpoint with shared-source input and partial coverage | Experimental, ESMFold, ColabFold |

**Supplementary Table 14** | Summary of external datasets and related sources.

| **Datasets** | **Source and evaluation definition** |
| --- | --- |
| CompData101, IndepData30, TestData36 | Released and/or found from DeepTMInter^12^. Interacting residues were determined with an interchain non-hydrogen atom distance < $4Å$, $5.5Å$, and $6Å$. Previously published membrane datasets. The separate retraining experiment excluded similar chains (Table 6a); the released-model evaluation retained development overlap (Table 6b). |
| Dset355, Dset448, Dset336 | Released and/or found from DELPHI^19^ and SCRIBER^20^. Native binary labels use inter-partner atom distance below the sum of van der Waals radii plus 0.5 Å. Dset336 is the positive-protein subset of Dset448. |
| Dset500 | Released and/or found from PITHIA^15^. Released binary protein-interface labels retained. |
| Btest31, UBtest31, Test315, Test60 | Released and/or found from GraphPPIS^7^. Native labels require surface relative solvent accessibility > 5% and binding-induced absolute accessible-surface-area loss > 1 Å 2 . Btest31 and UBtest31 are related bound/unbound views. |
| Test70 | DeepPPISP^14^. Native legacy solvent-accessibility-derived binary labels retained. |
| ScanNet Test | ScanNet^1^. Interchain heavy-atom distance ≤ 4 Å, including the source’s homologous-interface union across structures with ≥ 95% sequence identity. |
| Spatom Test709; Spatom Test80 | Spatom^11^. Original unbound residue labels retained for the Dockground and DBD5.5 sets, respectively. Test80 is an author-reproduction analysis. |
| DB5 and DB5.5 chain-complete and full-case views | Protein-protein docking benchmarks^21,22^. Bound-complex protein/polymer non-hydrogen contacts at < 6 Å were mapped to unbound coordinate residues by the frozen sequence-and-coordinate alignment. Nonprotein cofactors, glycans, nucleotides and ions were excluded from the contact calculation. Full-case views are subsets, not additional independent cohorts. |
| transient53 | MaSIF surface-label source^17^. Vertex interface labels were mapped to target-chain residues using the nearest atom coordinate and area weighting. A residue was positive when interface area exceeded 5 Å 2 and its fraction exceeded 0.04. The 2,720 residues without an assigned vertex were masked. |

**Supplementary Table 15 |** CHORUS evaluation on external datasets Released-model evaluation on complete external collections.

| **Dataset** | **AP-eligible / all contexts** | | **Evaluated residues** | **Positive fraction** | **Chain AP** | **Pooled AP** | **Pooled AUROC** |
| --- | --- | --- | --- | --- | --- | --- | --- |
| **Previously published transmembrane-protein datasets (training/validation overlap not excluded)** | | | | | | | |
| CompData101 | 101/101 | | 19,543 | 0.428 | 0.920 | 0.918 | 0.927 |
| IndepData30 | 30/30 | | 6,013 | 0.413 | 0.881 | 0.918 | 0.931 |
| TestData36 | 36/36 | | 6,870 | 0.371 | 0.935 | 0.937 | 0.948 |
| **Previously published transmembrane-protein datasets** | | | | | | | |
| CompData101 | 101/101 | 19,543 | | 0.428 | 0.770 | 0.752 | 0.803 |
| IndepData30 | 30/30 | 6,013 | | 0.413 | 0.741 | 0.744 | 0.821 |
| TestData36 | 36/36 | 6,870 | | 0.371 | 0.817 | 0.844 | 0.900 |
| **General-protein datasets** | | | | | | | |
| Btest31 | 31/31 | | 6,569 | 0.132 | 0.348 | 0.293 | 0.734 |
| DB5.5 chain-complete | 311/331 | | 68,323 | 0.102 | 0.253 | 0.133 | 0.627 |
| DB5 chain-complete | 263/281 | | 58,110 | 0.105 | 0.272 | 0.139 | 0.628 |
| Dset336 | 336/336 | | 84,941 | 0.186 | 0.482 | 0.470 | 0.784 |
| Dset355 | 247/355 | | 95,940 | 0.120 | 0.473 | 0.384 | 0.793 |
| Dset448 | 336/448 | | 116,500 | 0.136 | 0.482 | 0.410 | 0.792 |
| Dset500 | 500/500 | | 137,270 | 0.159 | 0.346 | 0.319 | 0.712 |
| ScanNet Test | 1070/1079 | | 288,199 | 0.172 | 0.407 | 0.375 | 0.743 |
| Spatom Test709 | 709/709 | | 222,398 | 0.082 | 0.286 | 0.177 | 0.718 |
| Spatom Test80 (reproduction) | 80/80 | | 23,090 | 0.088 | 0.264 | 0.160 | 0.697 |
| Test315 | 315/315 | | 65,331 | 0.143 | 0.349 | 0.288 | 0.709 |
| Test60 | 60/60 | | 13,144 | 0.158 | 0.390 | 0.328 | 0.727 |
| Test70 | 70/70 | | 11,791 | 0.198 | 0.419 | 0.367 | 0.710 |
| transient53 | 53/53 | | 14,227 | 0.104 | 0.266 | 0.164 | 0.634 |
| UBtest31 | 31/31 | | 6,654 | 0.126 | 0.328 | 0.263 | 0.721 |
| **Additional nested docking views** | | | | | | | |
| DB5.5 full-case subset | 103/109 | | 21,093 | 0.116 | 0.316 | 0.184 | 0.658 |
| DB5 full-case subset | 91/95 | | 18,287 | 0.121 | 0.332 | 0.193 | 0.662 |

**Supplementary Table 16 |** Sequence and structural coverage of evaluated soluble proteins.

| **Protein (PDB)** | **Construct aa** | **Observed instances** | **Max. training SI** | **AP / prevalence** |
| --- | --- | --- | --- | --- |
| Cortexillin I (1D7M) | 101 | 202 | 0.287 | 1.713 |
| COMP (1VDF) | 46 | 230 | 0.278 | 1.143 |
| Tetrabrachion (1FE6) | 52 | 205 | 0.286 | 1.330 |
| Eglin C (1CSE) | 70 | 63 | 0.271 | 3.468 |
| Subtilisin (1CSE) | 274 | 274 | 0.258 | 1.686 |
| Stefin B (1STF) | 98 | 98 | 0.291 | 4.082 |
| Papain (1STF) | 212 | 212 | 0.264 | 1.232 |
| Im9 (1EMV) | 86 | 83 | 0.256 | 1.863 |
| Colicin E9 DNase (1EMV) | 134 | 131 | 0.250 | 4.210 |
| Ribonuclease inhibitor (1A4Y) | 460 | 920 | 0.251 | 4.627 |
| Angiogenin (1A4Y) | 123 | 246 | 0.260 | 1.231 |

**Supplementary Table 17 |** Evaluation on soluble interfaces in terms of primary endpoint and operating-threshold behaviour.

| **Protein (PDB)** | **Copies** | **Prevalence** | **AP** | **AUROC** | **F1,+** | **MCC** | **Called +** | **Hydropathy AP/AUROC** |
| --- | --- | --- | --- | --- | --- | --- | --- | --- |
| Cortexillin I (1D7M) | 2 | 0.554 | 0.950 | 0.943 | 0.794 | 0.484 | 0.842 | 0.638 / 0.531 |
| COMP (1VDF) | 5 | 0.870 | 0.993 | 0.956 | 0.961 | 0.668 | 0.913 | 0.949 / 0.767 |
| Tetrabrachion (1FE6) | 4 | 0.742 | 0.987 | 0.960 | 0.852 | 0.000 | 1.000 | 0.842 / 0.606 |
| Eglin C (1CSE) | 1 | 0.222 | 0.771 | 0.942 | 0.583 | 0.494 | 0.540 | 0.217 / 0.445 |
| Subtilisin (1CSE) | 1 | 0.117 | 0.197 | 0.662 | 0.170 | 0.112 | 0.055 | 0.112 / 0.488 |
| Stefin B (1STF) | 1 | 0.194 | 0.791 | 0.940 | 0.692 | 0.634 | 0.337 | 0.224 / 0.590 |
| Papain (1STF) | 1 | 0.137 | 0.169 | 0.469 | 0.067 | 0.173 | 0.005 | 0.139 / 0.536 |
| Im9 (1EMV) | 1 | 0.265 | 0.494 | 0.697 | 0.506 | 0.316 | 0.783 | 0.287 / 0.551 |
| Colicin E9 DNase (1EMV) | 1 | 0.160 | 0.675 | 0.869 | 0.609 | 0.529 | 0.191 | 0.149 / 0.477 |
| Ribonuclease inhibitor (1A4Y) | 2 | 0.089 | 0.412 | 0.891 | 0.000 | 0.000 | 0.000 | 0.079 / 0.339 |
| Angiogenin (1A4Y) | 2 | 0.305 | 0.375 | 0.618 | 0.206 | 0.033 | 0.130 | 0.256 / 0.390 |

**Note:** The hydropathy AP/AUROC column represents reference ranking in which residues were ordered by their raw Kyte-Doolittle hydropathy values (Methods for details). More hydrophobic residues ranked higher, with no window averaging, sign selection, or tuning. It used the same observed residues with 5$Å$ primary-partner labels.

**Supplementary Table 18 |** AP/AUROC performance calculated at different distance-based cutoffs on soluble protein interfaces.

| **Protein (PDB)** | **4 Å** | **5 Å** | **5.5 Å** | **7 Å** |
| --- | --- | --- | --- | --- |
| Cortexillin I (1D7M) | 0.911 / 0.924 | 0.950 / 0.943 | 0.954 / 0.931 | 0.992 / 0.788 |
| COMP (1VDF) | 0.958 / 0.880 | 0.993 / 0.956 | 0.994 / 0.943 | 0.997 / 0.867 |
| Tetrabrachion (1FE6) | 0.975 / 0.948 | 0.987 / 0.960 | 0.985 / 0.946 | 0.987 / 0.812 |
| Eglin C (1CSE) | 0.770 / 0.946 | 0.771 / 0.942 | 0.771 / 0.942 | 0.830 / 0.901 |
| Subtilisin (1CSE) | 0.183 / 0.696 | 0.197 / 0.662 | 0.197 / 0.662 | 0.270 / 0.618 |
| Stefin B (1STF) | 0.640 / 0.921 | 0.791 / 0.940 | 0.779 / 0.931 | 0.841 / 0.929 |
| Papain (1STF) | 0.140 / 0.445 | 0.169 / 0.469 | 0.177 / 0.486 | 0.256 / 0.499 |
| Im9 (1EMV) | 0.494 / 0.697 | 0.494 / 0.697 | 0.485 / 0.660 | 0.556 / 0.593 |
| Colicin E9 DNase (1EMV) | 0.637 / 0.865 | 0.675 / 0.869 | 0.688 / 0.878 | 0.729 / 0.878 |
| Ribonuclease inhibitor (1A4Y) | 0.386 / 0.901 | 0.412 / 0.891 | 0.424 / 0.895 | 0.477 / 0.828 |
| Angiogenin (1A4Y) | 0.347 / 0.679 | 0.375 / 0.618 | 0.388 / 0.611 | 0.488 / 0.627 |

**Supplementary Table 19 |** Evaluation on respiratory membrane proteins. Taking 8UGN as a case, in Complex I, CHORUS precisely delineated the inter-subunit interfaces of the proton-pumping core subunits ND1 (chain H, AP = 0.998) and ND5 (chain L, AP = 0.993), as well as the transmembrane bridging subunit NDUFA11 (chain Y, AP = 0.836). In Complex III, the central catalytic core Cytochrome b (chain VA) achieved near-perfect interface recovery (AP = 0.989). In Complex IV, the core catalytic subunits COX1 (chain PB, AP = 0.985) and COX3 (chain RB, AP = 0.990) matched native contact maps with extraordinary fidelity.

| **PDB** | **Entities / copies** | **Contact fraction** | **Residues** | **AP** | **AUROC** | **F1** | **MCC** | **Best-member AP** | **ΔAP** |
| --- | --- | --- | --- | --- | --- | --- | --- | --- | --- |
| 8UGH | 38/46 | 0.687 | 7,203 | 0.983 | 0.967 | 0.949 | 0.816 | 0.983 | +0.0003 |
| 8UGJ | 38/57 | 0.683 | 8,762 | 0.984 | 0.971 | 0.948 | 0.816 | 0.983 | +0.0004 |
| 8UGN | 38/78 | 0.687 | 12,337 | 0.984 | 0.970 | 0.948 | 0.817 | 0.982 | +0.0019 |
| 8UGR | 38/92 | 0.689 | 14,400 | 0.979 | 0.962 | 0.942 | 0.792 | 0.978 | +0.0011 |
| 9I6F | 11/11 | 0.620 | 1,556 | 0.945 | 0.936 | 0.890 | 0.743 | 0.941 | +0.0034 |
| 9I7U | 11/11 | 0.645 | 1,563 | 0.966 | 0.958 | 0.921 | 0.776 | 0.970 | -0.0038 |

**Supplementary Table 20 |** Evaluation across diverse membrane-protein systems.

| **Case** | **PDB** | **Entities/copies** | **Residues** | **AP** | **AUROC** | **F1** | **MCC** |
| --- | --- | --- | --- | --- | --- | --- | --- |
| photosynthetic PSII–LHCII | 5XNL | 21/50 | 8,238 | 0.9504 | 0.9719 | 0.8865 | 0.7393 |
|  | 9HD7 | 26/60 | 9,486 | 0.9004 | 0.9212 | 0.8382 | 0.6439 |
| STING | 9DAN | 1/4 | 1,272 | 0.8339 | 0.8763 | 0.7334 | 0.4818 |
|  | 9DAT | 1/4 | 1,272 | 0.8452 | 0.8791 | 0.7421 | 0.4867 |
|  | 9DAV | 1/4 | 1,272 | 0.8445 | 0.8793 | 0.7531 | 0.5024 |
|  | 9DAW | 1/4 | 1,286 | 0.8574 | 0.8873 | 0.7494 | 0.4964 |
|  | 9SWM | 1/2 | 642 | 0.9190 | 0.9297 | 0.6291 | 0.3676 |
|  | 9SWQ | 1/2 | 642 | 0.8981 | 0.9068 | 0.6229 | 0.3197 |
|  | 9SWR | 1/6 | 1,898 | 0.8274 | 0.8780 | 0.6038 | 0.2965 |
|  | 9SWS | 1/4 | 1,264 | 0.8332 | 0.9060 | 0.6049 | 0.3328 |
| PINK1–TOM–VDAC | 9EIH | 6/20 | 1,878 | 0.7888 | 0.8592 | 0.6667 | 0.4455 |
|  | 9EII | 6/10 | 937 | 0.8012 | 0.8766 | 0.6888 | 0.4847 |
|  | 9EIJ | 6/11 | 985 | 0.7532 | 0.8315 | 0.6504 | 0.4232 |
| ER multipass translocon | 7TM3 | 7/7 | 1,739 | 0.6873 | 0.8454 | 0.5895 | 0.4088 |
|  | 7TUT | 9/9 | 2,025 | 0.6497 | 0.8133 | 0.5629 | 0.3547 |
| SCN5A CTD complexes | 4DCK | 1/1 | 153 | 0.6634 | 0.7288 | 0.5289 | 0.2566 |
|  | 4JQ0 | 1/1 | 142 | 0.7371 | 0.8221 | 0.6226 | 0.4363 |
|  | 4OVN | 1/1 | 146 | 0.6569 | 0.8361 | 0.5051 | 0.3813 |
|  | 6MUD | 1/1 | 128 | 0.4803 | 0.9023 | 0.4000 | 0.3974 |

**Supplementary Table 21 |** Summary of coverage and scope for the CHORUS atlas.

| **Quantity** | **Count** |
| --- | --- |
| UniProt records | 80,619 |
| Unique sequences | 70,247 |
| Duplicate-sequence records | 10,372 |
| Residues in unique sequences | 30,933,614 |
| Residues after expansion to records | 33,722,948 |
| Complete-sequence records | 80,062 |
| Overlapping-window records | 557 |
| Records longer than the supervised training maximum | 189 |
| Maximum sequence length (residues) | 14,507 |
| Predicted-positive residue instances | 8,973,552 |
| Exact supervised-training sequence matches | 176 |
| Retained match: 50% to below 100% | 13,709 |
| Retained match: 30% to below 50% | 6,703 |
| Retained match: below 30% | 49,417 |
| No retained search hit | 242 |
